# Reassessment of body temperature and thermoregulation strategies in Mesozoic marine reptiles

**DOI:** 10.1101/2024.07.26.605303

**Authors:** Nicolas Séon, Peggy Vincent, Lene Liebe Delsett, Eve Poulallion, Guillaume Suan, Christophe Lécuyer, Aubrey Jane Roberts, François Fourel, Sylvain Charbonnier, Romain Amiot

## Abstract

Ichthyosauria, Plesiosauria and Metriorhynchidae were apex predators in Mesozoic oceanic trophic networks. Previous stable oxygen isotope studies suggested that several taxa belonging to these groups were endothermic and for some of them homeothermic organisms. However, these conclusions remain contentious owing to the associated uncertainties regarding the δ^18^O value and oxygen isotope fractionation relative to environmental sea water. Here, we present new hydroxylapatite phosphate δ^18^O values (δ^18^O_p_) of Ichthyosauria, Plesiosauria and Metriorhynchidae (Middle Jurassic to Early Cretaceous) recovered from mid- to high-paleolatitudes to better constrain their thermophysiology and investigate the presence of regional heterothermies. The intra-skeletal δ^18^O_p_ variability failed to reveal distinct heterothermic patterns within any of the specimens, indicating either intra-body temperature homogeneity or an overriding diagenetic overprint of the original biological δ^18^O_p_ bone record. Body temperature estimates have then been reassessed from new and published δ^18^O_p_ values of well-preserved isolated teeth, recently revised Mesozoic latitudinal δ^18^O oceanic gradient and ^18^O-enrichment factor of fully aquatic air-breathing vertebrates. Our results confirm that Ichthyosauria were homeothermic endotherms (31°C to 41°C), while Plesiosauria were likely poikilothermic endotherms (27°C to 34°C). The new body temperature estimates of the Metriorhynchidae (25°C to 32°C) closely follow ambient temperatures and point to poikilothermic strategy with no or little endothermic abilities. These results improve our understanding of Mesozoic marine reptile thermoregulation and indicate that due to their limited body temperature variations, the δ^18^O_p_ values from Ichthyosauria fossil remains could be used as valuable archives of Mesozoic oceans δ^18^O_sw_ values that may help improve palaeoenvironmental and palaeoclimatic reconstructions.

**Non-technical abstract:** Some marine reptiles from the Mesozoic such as ichthyosaurs, plesiosaurs and metriorhynchids, were capable of reaching elevated body temperatures and for some of them to maintain it few degrees above that of their marine environment, a characteristic similar to that observed in modern cetaceans. Nevertheless, the estimation of their body temperature from the chemical oxygen signature of their fossil remains (bones and teeth) is accompanied by uncertainties associated with the chemical oxygen signature of the surrounding water and the mineralisation processes of the bones and teeth. In this study, new data were collected from four ichthyosaurs, three plesiosaurs and one metriorhynchid in order to gain a deeper understanding of the mechanisms by which these marine reptiles were able to maintain a body temperature higher than that of their environment. The chemical signatures of oxygen in the bones and teeth of the specimens did not exhibit any discernible patterns indicative of specific zones of heat production or loss, as observed in modern marine vertebrates. Concurrently, we reassessed the estimated body temperature of these marine reptiles, thereby corroborating the hypothesis that ichthyosaurs were homeothermic endotherms. Conversely, our novel estimates suggest that plesiosaurs were likely poikilothermic endotherms, whereas metriorhynchids were probably also poikilothermic endotherms but with a limited capacity for heat production. Finally, the narrow range of body temperatures maintained by ichthyosaurs indicates that the oxygen chemical signature of fossilised remains could serve as a valuable marker for reconstructing variations in the oxygen isotope composition of the Mesozoic oceans, paving the way to enhance our understanding of the environment and climate of this period in Earth’s history.

## Introduction

During the Mesozoic era (251.9 to 66.0 million years ago, Ma), marine reptiles such as ichthyosaurs, plesiosaurs and metriorhynchids were distributed worldwide and played a key role in the trophic networks. Paleobiogeographic (Kear 2006a; Bardet et al. 2014; Vavrek et al. 2014; Delsett et al. 2016; Rogov et al. 2019; Zverkov et al. 2021), osteo-histological (Ichthyosauria: de Buffrénil & Mazin 1989; de Buffrénil & Mazin 1990; Anderson et al. 2019; Plesiosauria: Wiffen et al. 1995; Delsett and Hurum 2012; Fleischle et al. 2018), geochemical (Bernard et al. 2010; Séon et al. 2020; Leuzinger et al. 2023), and modelling studies (Brice and Grigg 2023) all indicate that Ichthyosauria Blainville, 1835 and Plesiosauria Blainville, 1835 were endothermic and probably homeothermic organisms, much like extant Cetacea Brisson, 1762. They would have been able to produce enough body heat to raise their body temperature above that of the environment in which they lived (Bernard et al. 2010; Séon et al. 2020; Leuzinger et al. 2023). High and constant body temperature in Ichthyosauria would have been facilitated by their fusiform morphology favouring heat retention, and the presence of a layer of fibroadipose tissue surrounding the trunk (Lindgren et al. 2018; Delsett et al. 2022). Evidence for adipose tissue is lacking for Plesiosauria for which very few specimens preserving soft tissues have been found (Vincent et al. 2017). Thermophysiological studies of Metriorhynchidae Fitzinger, 1843, a group of fully aquatic marine mesozoic crocodylomorphs, have not led to a consensus. The fossil occurrences confined to tropical paleo-latitudes (Bardet et al. 2014) and the osteo-histological findings from studies by Hua and de Buffrénil (1996) and de Buffrénil et al. (2021) suggested that Metriorhynchidae had an ectothermic poikilothermic thermoregulatory strategy similar to that of modern crocodylomorphs. Conversely, the oxygen isotope analyses of Séon et al. (2020) suggested that Metriorhynchidae were able to raise their body temperature few degrees above that of the ambient environment by metabolic heat production but could not maintain a constant body temperature, indicating they were poikilothermic endotherms like extant tunas (Block and Finnerty 1994; Graham and Dickson 2004).

Previous estimates of body temperature reconstruction of Ichthyosauria, Plesiosauria, and Metriorhynchidae, calculated from the phosphate oxygen isotope composition of the hydroxylapatite of their bones and teeth (δ^18^O_p_), assumed a constant global oceanic δ^18^O value (δ^18^O_sw_) of −1 ± 1‰ (Bernard et al. 2010; Séon et al. 2020; Leuzinger et al. 2023; **Supplementary information 1** for details about isotope-based body temperature estimations), an assumption over-simplified given latitudinal gradient in δ^18^O_sw_ values recorded in Jurassic and Cretaceous seas (Alberti et al. 2017, 2020; Letulle et al. 2022). Furthermore, recent work has shown that the oxygen isotope compositions of body water from extant air-breathing fully marine species [*Orcinus orca* (Linnæus, 1758), *Tursiops truncatus* (Montagu, 1821), Séon et al. 2023; and *Caretta caretta* (Linnæus, 1758), Séon 2023] are less ^18^O-enriched relative to their drinking water than that of semi-aquatic vertebrates (crocodiles, turtles) used previously to reconstruct body temperatures of extinct marine reptiles (Bernard *et al*., 2010, Séon *et al*., 2020; Leuzinger *et al*., 2023).

Previous osteo-histological and geochemical studies were exclusively based on isolated teeth or bone remains (de Buffrénil & Mazin 1990; Bernard et al. 2010; Fleischle et al. 2018; Séon et al. 2020) and thus provided only partial information on the thermoregulation of Ichthyosauria, Plesiosauria and Metriorhynchidae. A key aspect of thermophysiology is to define how constant body temperature is at the level of vital organs (homeotherm vs. poikilotherm: Clarke and Pörtner 2010; Furukawa et al. 2015; Lovegrove 2017) and the distribution of body temperature within the body, known as regional heterothermies (Irving and Hart 1957; Folkow and Blix 1987; Favilla et al. 2022). An investigation into the mechanisms employed by Ichthyosauria, Plesiosauria and Metriorhynchidae to regulate their body temperature allows for an understanding of their adaptation to their environment and the explanation of their stratigraphic occurrence in the fossil record in view of fluctuating environmental temperatures during the Mesozoic (Takashima et al. 2006; Dera et al. 2011; Wierzbowski et al. 2013). Such climate changes do not seem to have affected their diversity (Bardet 1994; Martin et al. 2014; Stubbs and Benton 2016). Moreover, defining precisely their thermoregulatory strategies opens the way to investigate their behaviour. For instance, homeothermic endotherms are typically active organisms, as they are fully independent of the temperature of their environment. Nevertheless, this requires a significant energy input (Clarke & Pörtner 2010). Conversely, some species that are considered as regional endotherms can produce heat locally, which can result in temperature heterogeneities, or regional heterothermies (Carey 1982; Block 1986; Dickson and Graham 2004; Graham and Dickson 2004). In swordfish, for example, the heat production located close to the eyes improves visual acuity in foraging cold environments of great depth (Block 1987; Fritsches et al. 2005), whereas heat production in locomotory muscles of tunas and lamnid sharks enables them to swim faster or migrate over longer distances than poikilothermic ectotherms organisms (Blank et al. 2007; Bernal et al. 2012; Watanabe et al. 2015; Harding et al. 2021). The mapping of intra-individual temperature variations, and thus regional heterothermies, through the use of oxygen isotopes enables the identification of heat-producing zones within an organism, in addition to the delineation of thermal windows, the preferred sites for heat loss, which serve to regulate body temperature (Séon et al. 2022, 2024).

In this study, we have performed an oxygen isotope analysis of 247 bones and teeth from eight complete or sub-complete specimens, enabling the first evaluation of possible regional heterothermies within the bodies of ichthyosaurs, plesiosaurs and metriorhynchids. We have also reassessed body temperature estimates of Ichthyosauria, Plesiosauria and Metriorhynchidae using new and published δ^18^O_p_ teeth data, a ^18^O-enrichment of fully aquatic animals and considering a global δ^18^O_sw_ gradient for the Jurassic and Cretaceous seas.

### Institutional abbreviations

MHNLM *Muséum d’Histoire Naturelle Le Mans*

MPV *Musée Paléontologique de Villers-sur-mer*

PMO *Palaeontological collections of the Natural History Museum, University of Oslo, Norway*

## Material and Methods

### Sampled specimens

Four Ichthyosauria, three Plesiosauria and one Metriorhynchidae specimens have been sampled for their intra-skeletal δ^18^O_p_ variability, the taxonomic affiliation, the collection number, the size estimate, the stratigraphic age, the locality and the type of sampled material of each marine reptile specimen are reported in **Table 1**. In addition to these complete and sub-complete specimens, two teeth of Metriorhynchidae indet. specimens from the Marnes de Dives Formation (Late Callovian, Vaches Noires Cliffs, France) were sampled for their δ^18^O_p_ values to reassess metriorhynchid body temperatures (**Table 1**), as well as a pachycormid fish tooth belonging to *Hypsocormus* Wagner 1863 recovered from the same stratigraphical layer of the specimen of Ichthyosauria indet. from “Les Ardilles” (Kimmeridgian, Auxerre, France) to determine the oceanic paleotemperature of this deposit. Overall preservation and appearance of the studied specimens varies between the localities. The Ichthyosauria indet. specimen from “Les Ardilles” (France) is slightly deformed and marked by numerous fractures secondarily filled with calcite (Mazin and Pavy 1995). The skeletal remains of the specimen of *Metriorhynchus* aff. *superciliosus* from the Vaches Noires Cliffs (France) seem to have been little affected by compaction, except for the skull, which shows a few cracks with surface recrystallization (Le Mort et al. 2022). The ichthyosaurs (PMO 222.655, PMO 222.667, PMO 222.669) and plesiosaurs (PMO 212.662, PMO 222.663) from the Slottsmøya Member (Spitsbergen) are partly eroded, probably by the action of suspended particles and sediments transported by bottom currents (Martill 1985; Reisdorf et al. 2012) and show recrystallizations of calcite and barite in the pores of the skeletal elements (Kihle et al. 2012; Delsett et al. 2016). The skeletal remains are also for the most part highly fractured, partly due to local faults within the deposit and to frost weathering. For instance, the specimen of *Keilhauia nui* (PMO 222.655) was crossed by large fractures (Delsett et al. 2017). Its skeletal elements are particularly fractured and have a sandy appearance.

### Sampling method

Approximately 40 mg of each skeletal element and tooth were grounded into fine powder using a Dremel^TM^ micro-drill equipped with a diamond-studded drill bit; cortical bone was selected to maximise the chances of preserving a biological isotopic record. Skeletal elements of complete and sub-complete specimens have been grouped in five categories, as follows: “Skull” groups all the bone forming the skull and the mandible, “Cervical region” groups the cervical vertebra only in Plesiosauria, “Dorsal region” groups the dorsal vertebrae, ribs and skeletal elements forming the pectoral and pelvic girdles, “Caudal region” groups the caudal vertebrae and the “Appendicular region” groups humerus, femur, radius, ulna, tibia, fibula, metacarpals, metatarsals, phalanges, and accessory bone elements located in the limbs. For the specimen of *Colymbosaurus svalbardensis* (PMO 222.663), the “Appendicular region” has been split into anterior right limb (ARL), anterior left limb (ALL), posterior right limb (PRL) and posterior left limb (PLL) since most of the elements were found in articulation (Delsett et al. 2016). For Ichthyosauria and *Metriorhynchus* aff. *superciliosus* specimens, cervical vertebrae were included in the “Dorsal region”. A total of 247 skeletal elements and teeth were sampled, averaging about 20 and 40 samples per specimen. In addition to the bone and teeth samples, a sample of the recrystallisation mineral observed in one dorsal vertebra from the Ichthyosauria indet. Specimen from “Les Ardilles” was taken.

### Biogenic hydroxylapatite P_2_O_5_ content and phosphate group oxygen isotope analysis

Cortical bone and teeth powders (n = 247) were prepared using the wet chemical procedure detailed by Crowson et al. (1991) and modified by Lécuyer et al. (1993), which involves the isolation of phosphate ions (PO_4_ ^3-^) from the bioapatite which are then precipitated as silver phosphate crystals (Ag_3_PO_4_). For each sample, 20–30 mg of cortical bone or tooth enamel powder was dissolved in 2 ml of 2M HF. The CaF_2_ residue was separated by centrifugation and the solution was neutralized by adding 2.2 ml of 2M KOH. Amberlite™ IRN78 anion-exchange resin beads were added to the solution to isolate the PO_4_^3-^ ions. After 24 h, the solution was removed, and the resin was rinsed with deionized water and eluted with 27.5 ml of 0.5M NH_4_NO3. After 4 h, the Amberlite™ IRN78 anion-exchange resin beads were removed from the solution. Then, 0.5 ml of NH_4_OH and 15 ml of an ammoniacal solution of AgNO_3_ were added and the solutions were placed in a thermostated bath at 70°C for 7 h, allowing the slow and quantitative precipitation of Ag_3_PO_4_ crystals. The Ag_3_PO_4_ crystals were filtered, dried and cleaned. For each sample, the P_2_O_5_ content, a proxy for estimating the overall preservation of the mineralized skeletal elements bioapatite in fossil specimens (Nemliher et al. 2004), have been estimated. It was calculated from the recovered mass of silver phosphate (Ag_3_PO_4_) after the chemical purification of the samples and phosphate chemical yields obtained for the NIST SRM 120c standards (P_2_O_5_= 33.34 wt%), assuming stoichiometric conversion of hydroxylapatite phosphate ions into silver phosphate (Lécuyer et al. 2003).

For isotope composition measurements, five aliquots of 300 ± 20 μg of Ag_3_PO_4_ for each sample were mixed with 400 ± 50 μg of graphite in silver foil capsules. Oxygen isotope ratios were measured using a high temperature vario PYRO cube^TM^ elemental analyser (EA), equipped with the “purge and trap” technology (Fourel et al. 2011) and interfaced in continuous flow mode to an IsoPrime^TM^ isotope ratio mass spectrometer (Elementar UK Ltd Cheadle, UK). To calibrate the measurements, the NBS127 (barium sulfate precipitated from sea water from Monterey Bay, California, USA, Sigma-Aldrich Co.), and a silver phosphate precipitated from the international standards NIST SRM 120c (natural Miocene phosphorite from Florida, Sigma-Aldrich Co.) were used. Their respective δ^18^O_p_ value was set at the certified value of +9.3‰ V-SMOW (Hut 1987; Halas and Szaran 2001) and fixed at +21.7‰ V-SMOW (Vienna Standard Mean Ocean Water) according to Lécuyer et al. (1993), Chenery et al. (2010) and Halas et al. (2011), for correction of instrumental mass fractionation during CO isotope analysis. Along with the silver phosphate samples derived from bones and teeth hydroxylapatite, silver phosphates precipitated from standard NIST SRM 120c were repeatedly analysed (δ^18^O_p_ = 21.7 ± 0.3‰, n = 59) to ensure that no isotope fractionation occurred during the wet chemistry. Values are reported as delta permil values expressed with respect to V-SMOW.

### Oxygen and carbon isotope analysis of biogenic hydroxylapatite carbonate and carbonate content

Seventy-one (n = 71) powders were pre-treated to measure the oxygen and carbon isotope compositions of the carbonate group of the bone hydroxylapatite (Plesiosauria: n = 40, Ichthyosauria: n = 29, *Metriorhynchus* aff. *superciliosus*: n = 1 and n = 1 from the pachycormid fish tooth of *Hypsocormus* sp.). The protocol used was that of Koch et al. (1997) and consisted in removing organic matter through a chemical reaction between bone powder and 0.4 mL of 2% sodium hypochlorite (NaOCl) for 24 h (0.4 mL NaClO per 10 mg of bone powder). The solution was centrifuged and rinsed three times with Ultrapure^TM^ water, before 0.1 M acetic acid (CH_3_COOH) was added for 24 h to remove any secondary carbonates that may have precipitated during the organic removal procedure. The solution was again centrifuged and rinsed three times. Then, the powders were dried at 50°C for 48 h before being collected. For each sample, three replicates of 2 mg were weighed into 3.7 mL round-bottomed glass vials and sealed (LABCO UK Exetainer®). Oxygen (δ^18^O_c_) and carbon (δ^13^C_c_) isotope composition of the carbonate group of bone and teeth bioapatite were measured using an isoFLOW-type preparation system (Elementar GmbH-Germany) connected in continuous flow to a precisION isotope ratio mass spectrometer (Elementar UK^TM^). Each of the pre-treated samples was reacted with saturated anhydrous phosphoric acid (H_3_PO_4_) prepared according to the protocol of McCrea (1950). The reaction took place at a constant temperature of 90°C. The CO_2_ generated during acid digestion of the carbonate sample is then transferred to the mass spectrometer. Carbonate content (CO_3_^2-^ %wt) of the bioapatite samples were measured based on the peak area of CO_2_ detected by the mass spectrometer. Isotope measurements are corrected for instrumental drift and calibrated using two calcite isotopic standards: a Carrara marble (laboratory standard) with values of δ^18^O_V-PDB_ = −1.84 ‰ and δ^13^C_V-PDB_ = +2.03 ‰ (Fourel et al. 2015), the NBS18 (international standard) whose values are δ^18^O_V-PDB_ = −23.2 ‰; δ^13^C_V-PDB_ = −5.01 ‰ (Friedman et al. 1982; Hut 1987; Coplen et al. 2006) and NIST SRM 120c (δ^18^O_V-PDB_= −1.13‰, δ^13^C_V-PDB_ = −6.27‰, (Passey et al. 2007). For bone and tooth apatite, the acid fractionation factor α(CO_2_–apatite carbonate) of 1.00773 determined for the NIST SRM 120c phosphate rock reference material was selected (Passey et al. 2007). Values are reported as delta permil values expressed with respect to V-SMOW for oxygen and V-PDB (Vienna Pee Dee belemnite) for carbon.

### Raman spectroscopy

Characterization of the mineralogical composition of the samples has been carried out using an XploRA Raman microscope equipped with a diode-pumped Nd:YAG laser at 532 nm. For each sample analysed (n = 84), 10 spectra of 10-second were acquired at x100 magnification. The scattered light was detected in the range of 150 and 2,000 cm^-1^. The first-order band of a pure silicon reference material at 520.7 cm^-1^ was used to calibrate the spectrometer before each measurement. The full width at half maximum (FWHM), and the position of the ν1(PO_4_^3-^) fully symmetric stretching band have been measured to obtain information on the mineralogical composition and crystal structure of bioapatite (Pucéat et al. 2004; Thomas et al. 2007, 2011; Dal Sasso et al. 2018; Barthel et al. 2020; Kral et al. 2022).

### Body temperature estimates

Body temperatures of Ichthyosauria, Plesiosauria and Metriorhynchidae have been re-estimated from published tooth δ^18^O_p_ values (Anderson et al. 1994; Bernard et al. 2010; Séon et al. 2020) as well as newly measured values from two teeth belonging to specimens of Metriorhynchidae indet. from “Les Vaches Noires Cliffs” (Late Callovian, Villers-sur-Mer, France), and the teeth from the three specimens of ichthyosaur referred to *Palvennia hoybergeti* (PMO 222.669), *Keilhauia* sp. (PMO 222.667) and Ichthyosauria indet. from “Les Ardilles”. The equation published by Lécuyer et al. (2013), adapted for air-breathing vertebrates, has been used and an average ^18^O-enrichment of +0.8 ± 0.9‰, for air-breathing fully aquatic marine vertebrates was applied (Séon et al. 2023). Paleolatitudes were reconstructed using the software developed by van Hinsbergen et al. (2015) and then used for the estimation of the δ^18^O_sw_ values from the equation published by Alberti et al. (2020). For each locality, sea surface temperature was calculated from the δ^18^O_p_ values of contemporary fish using equation of Lécuyer et al. (2013) considering that body temperature (Tb) is equal to sea water temperature (T_sw_), and that δ^18^O_bw_ ≈ δ^18^O_sw_ (Kolodny et al. 1983). The associated error in temperature calculation is equal to 3°C (based on the slope of 4.5 of the Lécuyer et al. (2013) equation used). As no contemporary fish tooth has been analysed for the Slottsmøya member fossil locality, the environmental temperature estimate has been done from the oxygen isotope composition of one brachiopod shell from Hammer et al. (2011) using the oxygen isotope fractionation equation provided by Letulle et al. (2023).

### Statistical analyses

We used descriptive statistics to explore body temperature re-estimates and intra-skeletal δ^18^O_p_ variations in Mesozoic marine reptiles. The normality and homoscedasticity (the uniformity of the error associated with the variance for each of the values) of the δ^18^O_p_ values could not be verified. Considering the variable number of samples for each set (between 1 and 14 samples), the non-parametric Mann-Whitney-Wilcoxon test was used to compare median values between two sets of observations, each corresponding to a skeletal region. Statistical tests were performed using R software with a significance threshold set at *p*-value < 0.05. In order to ascertain the degree of correlation between two quantitative variables, Pearson’s correlation coefficient *r* was employed.

## Results

### Hydroxylapatite P_2_O_5_ content and phosphate oxygen isotope composition (δ^18^O_p_)

A total of two hundred and eighteen (n = 218) δ^18^O_p_ values were acquired from four ichthyosaurs, three plesiosaurs and one *Metriorhynchus* aff. *superciliosus* specimen in order to map regional heterothermies and reassess body temperatures. Almost all samples from each specimen could be analysed, except a few samples that failed to yield silver phosphate after wet chemistry preparation, especially from the specimen of *Keilhauia nui* (PMO 222.655). P_2_O_5_ content expressed in weight percentage (wt%) and measured oxygen isotope compositions from the phosphate group of each bone and tooth of all the specimens are reported in **Supplementary Table 1** and synthetised in **Table 2**. P_2_O_5_ content for all specimens ranged from 2.0% to 40.5% (**Table 2**). No significant differences were observed between the fossil deposits (**Supplementary Figure 1A**), nor between specimens if we exclude the *Keilhauia nui* specimen (PMO 222.655) for which the percentage of P_2_O_5_ is significantly lower (Mann-Whitney-Wilcoxon test, *p-*value < 0.001; **Supplementary Figure 1B**). Globally, there are no significant differences in P_2_O_5_ content along skeletal region, except for the Ichthyosauria indet. and *Colymbosaurus svalbardensis* specimens (Mann-Whitney-Wilcoxon test, *p-*value < 0.05; **Supplementary Figure 2**). Finally, no strong relationship between P_2_O_5_ content and δ^18^O_p_ values is observed in Ichthyosauria and in Plesiosauria specimens (Pearson correlation test, *r* = 0.32; **Supplementary Figure 3**).

The intra-skeletal variability of δ^18^O_p_ is illustrated in **Figure 1, 2** and **3**, respectively for Ichthyosauria, Plesiosauria and Metriorhynchidae. The *Hypsocormus* tooth recovered from the same stratigraphical layer as the specimen of Ichthyosauria indet. from “Les Ardilles” yielded a δ^18^O_p_ value of 18.1± 0.2 ‰ and the δ^18^O_p_ value of the Metriorhynchidae teeth VN6 and VN7 equal to 21.4 ± 0.2 ‰ and 19.8 ± 0.2 ‰ respectively (**Supplementary Table 2**). Intra-skeletal variability in δ^18^O_p_ values is observed for all the specimens of Ichthyosauria, the specimen of *Kimmerosaurus* sp. and the specimen of *Metriorhynchus* aff. *superciliosus* but these differences are not significative between body regions (Mann–Whitney–Wilcoxon test, *p*-value > 0.05, **Fig. 4**). However, a trend seems to emerge in specimen PMO 222.667 of *Keilhauia* sp. and *Palvennia hoybergeti* specimen PMO 222.669, in which the teeth have overall higher δ^18^O_p_ values than those of the bones, but the statistical significance of this observation could not be tested because of the small number of samples for each set (n < 5 for teeth). The only significant differences in δ^18^O_p_ values along body regions observed are in the *Colymbosaurus svalbardensis* (PMO 222.663) specimen. Mann-Whitney-Wilcoxon statistical test results (*p*-value < 0.05, **Fig. 4**) clearly indicate that skeletal elements in the anterior part of the skeleton (dorsal vertebrae, ribs, left and right forelimbs) have significantly higher δ^18^O_p_ values than skeletal elements in the posterior part (left and right hindlimbs and caudal vertebrae).

**Figure 1:**
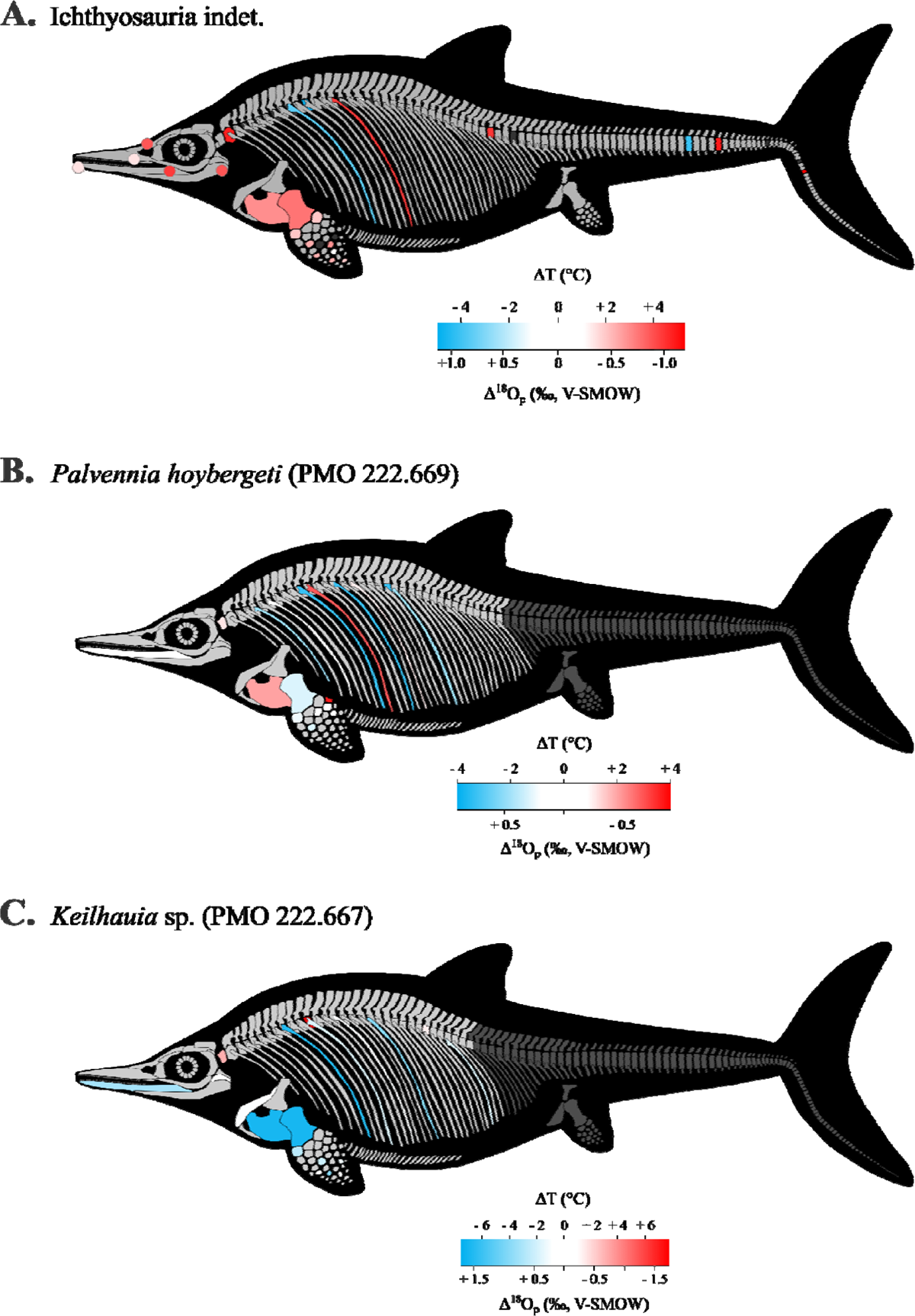
Regional heterothermies in ichthyosaurs. Phosphate oxygen isotope and temperature variability within the skeleton from **A.** the specimen of Ichthyosauria indet., **B.** the specimen (PMO 222.669) of *Palvennia hoybergeti* and **C.** the specimen (PMO 222.667) of *Keilhauia* sp. Color in bones correspond to the δ^18^O_p_ difference between bone and the mid-range value (δ^18^O_p-max_ + δ^18^O_p-min_ / 2) of the skeleton. For paired skeletal elements, the mean value is illustrated. Available skeletal elements are shown in light grey, while unavailable elements and skeletal elements with potentially altered δ^18^O_p_ values are shown in dark grey. Skeletal elements (e.g., ‘limb bones’, **Supplementary Table 1**) with unknown precise location are not illustrated while the location of vertebrae, ribs and phalanges which have not been found in articulation have been established arbitrarily. The representation of the organisms is not to scale.

**Figure 2:**
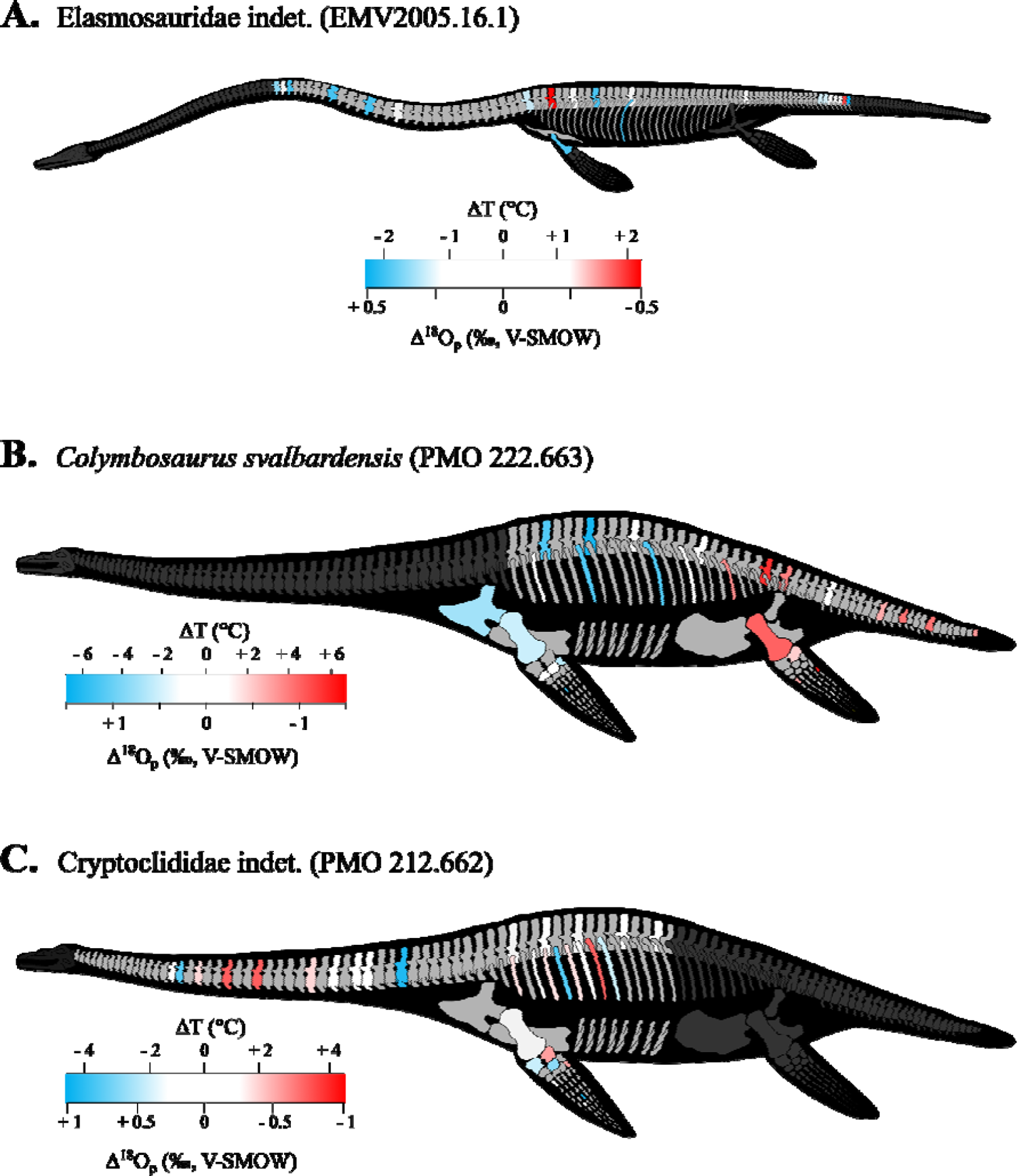
Regional heterothermies in plesiosaurs. Phosphate oxygen isotope and temperature variability within the skeleton from **A.** the specimen (MHNLM.2005.16.1) of Elasmosauridae indet., **B.** the specimen (PMO 222.663) of *Colymbosaurus svalbardensis* and **C.** the specimen (PMO 212.662) of Cryptoclididae indet. Colour in bones correspond to the δ^18^O_p_ difference between bone and the mid-range value (δ^18^O_p-max_ + δ^18^O_p-min_ / 2) of the skeleton. For paired skeletal elements, the mean value is illustrated. Available skeletal elements are shown in light grey, while unavailable elements and skeletal elements with potentially altered δ^18^O_p_ values are shown in dark grey. Skeletal elements (e.g., ‘limb bones’, **Supplementary Table 1**) with unknown precise location are not illustrated while the location of vertebrae, ribs and phalanges, which have not been found in articulation, have been established arbitrarily. The representation of the specimens is not to scale.

**Figure 3:**
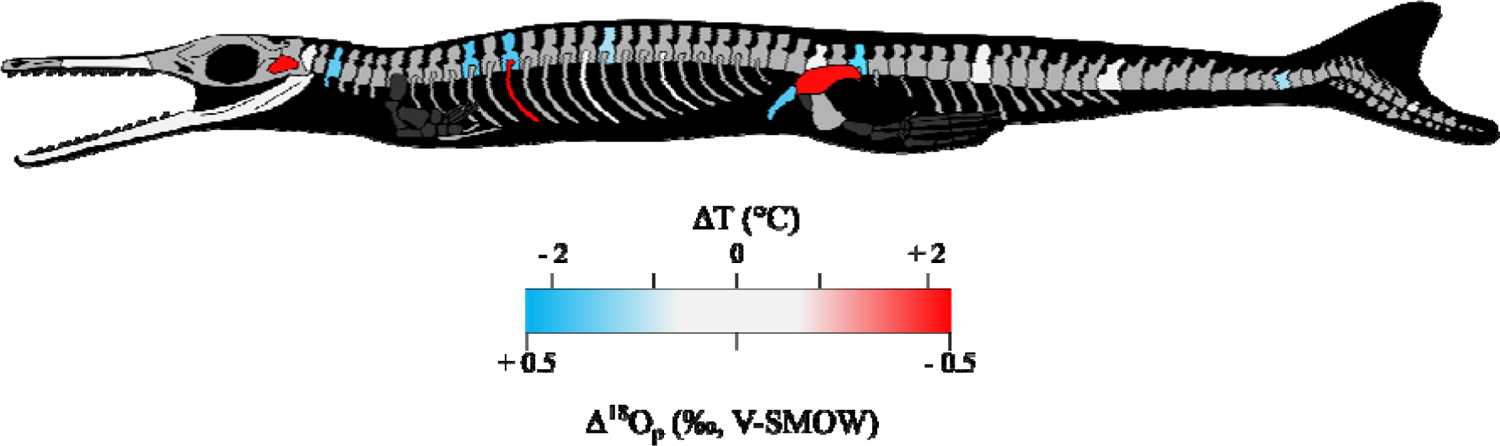
Regional heterothermies in *Metriorhynchus* aff. *superciliosus*. Phosphate oxygen isotope and temperature variability within the skeleton from the specimen (MPV 2010.3.610) of *Metriorhynchus* aff. *superciliosus*. Colour in bones correspond to the δ^18^O_p_ difference between bone and the mid-range value (δ^18^O_p-max_ + δ^18^O_p-min_ / 2) of the skeleton. When both vertebrae centra and neural spine have been measured, the mean value is illustrated. Available skeletal elements are shown in light grey, while unavailable elements are shown in dark grey. The representation of the specimen is not to scale.

Plesiosaur specimens (*Colymbosaurus svalbardensis*, Cryptoclididae indet.) have been recovered in articulation (Vincent et al. 2007; Delsett et al. 2016), consequently, the relation between δ^18^O_p_ values and distance from the trunk has been investigated (**Fig. 5A**). No relationship between δ^18^O_p_ values and position in the limbs (*Colymbosaurus svalbardensis*, Cryptoclididae indet.) as well as for the vertebral position in the neck (Cryptoclididae indet., Elasmosauridae indet.) have been observed (**Fig. 5B**). To sum up, phosphate oxygen isotope compositions reveal variations for all studied specimens but for most of them no significant differences in δ^18^O_p_ from the individual skeletal regions.

**Figure 4:**
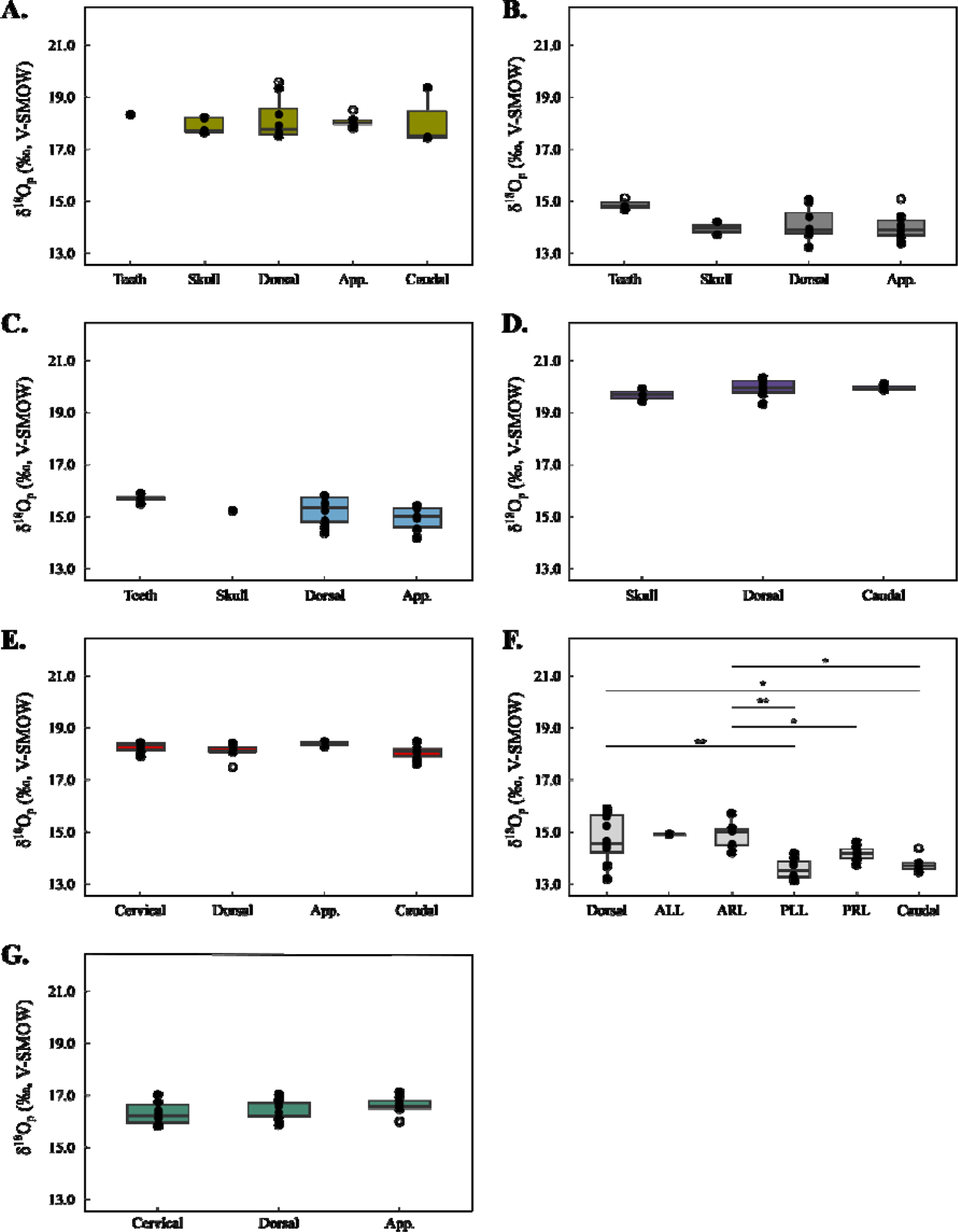
Boxplots showing the δ^18^O_p_ values distribution of bones sets for the specimens of Ichthyosauria (**A.** Ichthyosauria indet, **B.** *Keilhauia* sp., **C.** *Palvennia hoybergeti*), *Metriorhynchus* aff. *superciliosus* (**D.**) and Plesiosauria (**E.** Elasmosauridae indet., **F.** *Colymbosaurus svalbardensis* and **G.** Cryptoclididae indet.). Asterisks indicate the significance of the observed differences between pair of groups: * for *p*-value < 0.05, ** for *p*-value < 0.01 and *** for *p*-value *<*0.001. Outliers are plotted as white circles. The horizontal bars in the boxes correspond to the medians and the whiskers to the minimum and maximum values. The average analytical error for each bone analysed is of the order of 0.3‰. Abbreviations: App. = Appendicular region, ALL = Anterior Left Limb, ARL = Anterior Right Limb, PLL = Posterior Left Limb, PRL = Posterior Right Limb.

**Figure 5:**
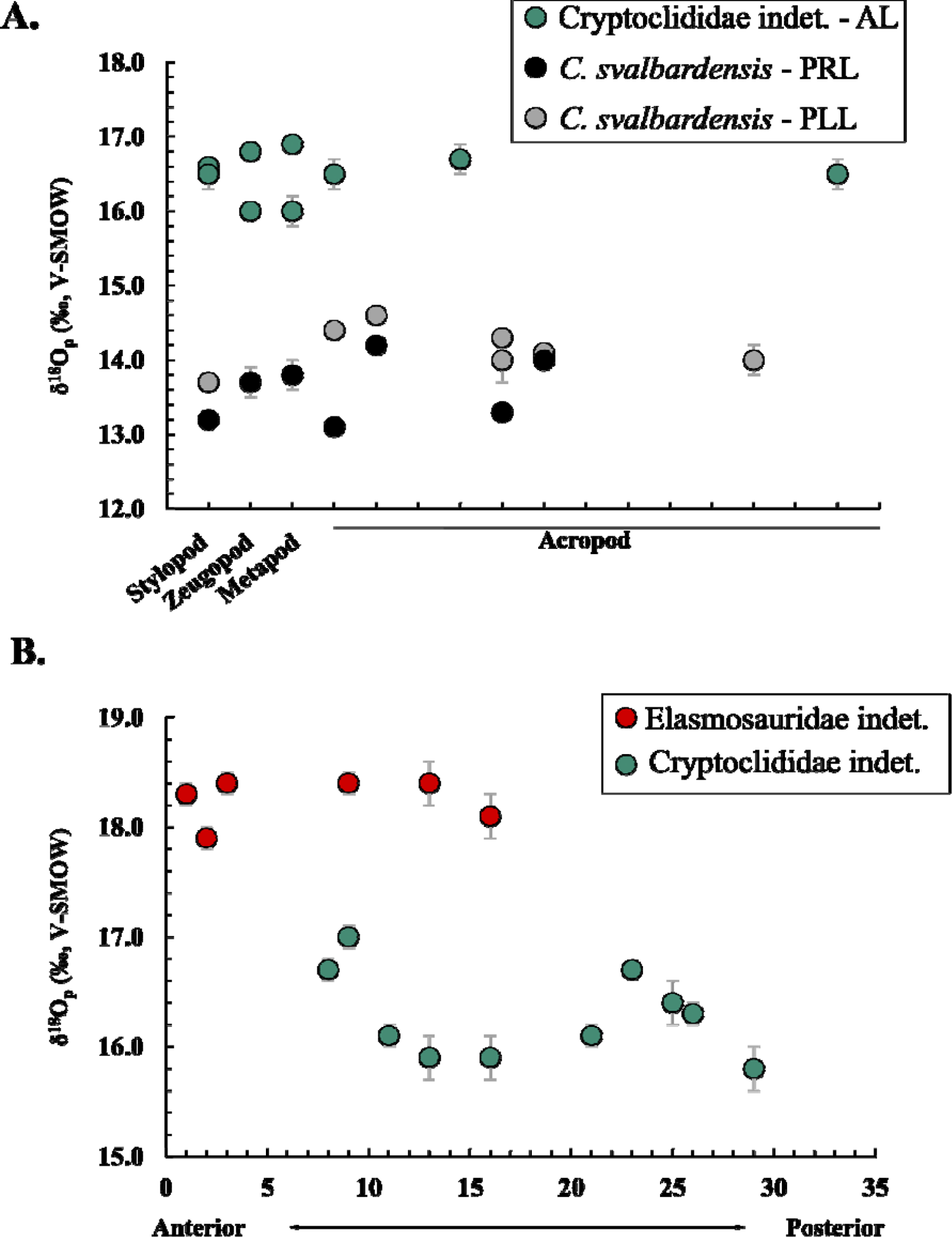
**A.** δ^18^O_p_ values (‰, V-SMOW) according to bone position within the limb for the specimen of Cryptoclididae indet. and the specimen of *Colymbosaurus svalbardensis*. Stylopod set corresponds to the femur or the humerus, Zeugopod set to radius and ulna or tibia and fibula, Metapod set to metacarpals or metatarsals and Acropod set to phalanges. Abbreviation: AL = Anterior limb indet., PRL = Posterior Right Limb and PLL = Posterior Left Limb. **B.** δ^18^O_p_ values (‰, V-SMOW) according to vertebra position within the cervical region for the specimen of Elasmosauridae indet. and the specimen of Cryptoclididae indet.

**Figure 6:**
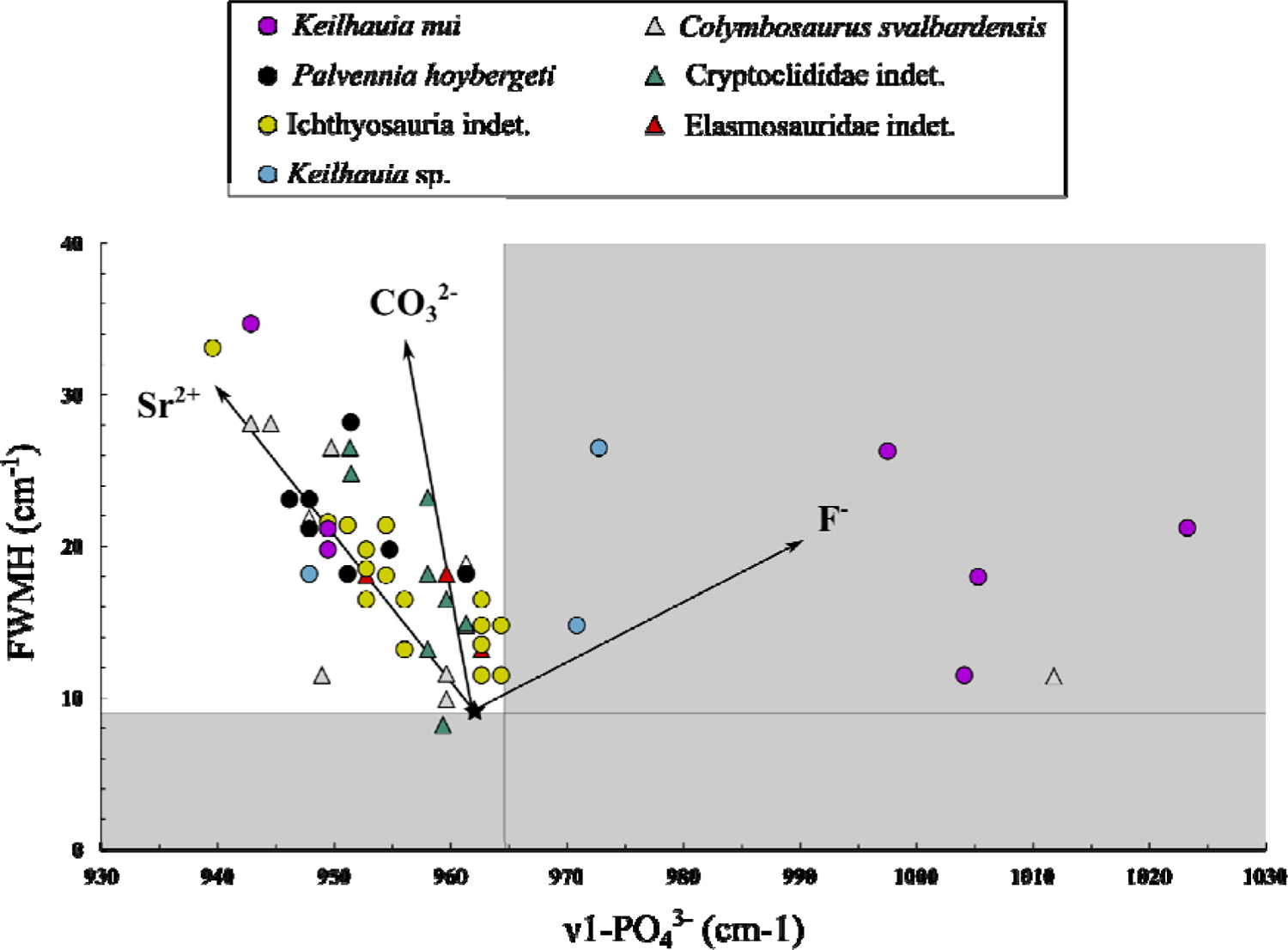
ν1(PO ^3-^) stretching band position and FWMH measured for Ichthyosauria and Plesiosauria bones and teeth (**Supplementary Table 3**). The grey zone corresponds to the combination of parameters which would indicate a mineralogical alteration of the hydroxylapatite of teeth and bones. The black arrows indicate the evolution of the position of the ν1(PO_4_^3-^) stretching band position and the FWMH depending on the type of ionic

### Carbonate hydroxylapatite oxygen and carbon isotope compositions (δ^18^O_c_ and δ^13^C_c_) and carbonate content (wt%)

Oxygen and carbon isotope compositions of the carbonate group of hydroxylapatite as well as carbonate content have been measured to assess the preservation of the oxygen isotope composition of the phosphate group. All the values are reported in **Supplementary Table 1** and a synthesis is provided in **Table 2** for each specimen. The tooth from the pachycormid fish *Hypsocormus* sp. recovered from the same stratigraphical layer as the specimen of Ichthyosauria indet. yielded to a δ^18^O_c_ value equals to 27.30 ± 0.20‰, V-SMOW, a δ^13^C_c_ value equals to −4.27 ± 0.15‰, V-PDB, and a carbonate content of 4%. The δ^18^O_c_ of calcite recrystallisation sampled in the dorsal vertebra of the specimen of Ichthyosauria indet. from “Les Ardilles” is equal to 24.20 ± 0.10‰, V-SMOW and to −0.10 ± 0.10‰, V-PDB for δ^13^C_c_.

### Mineralogical characteristics of bones and teeth from Mesozoic marine reptiles

Raman spectroscopy has been used to assess possible mineralogical changes associated to fossilization and diagenetic processes. The Raman spectral parameters from ichthyosaurian and plesiosaurian bone elements and teeth have been reported in **Supplementary Table 3** and illustrated in **Figure 6**. The values of position of the ν1(PO_4_^3-^) stretching band range from 939.6 cm^-1^ to 1023.4 cm^-1^ whereas FWHM values range from 8.2 to 34.7 cm^-1^ (**Supplementary Table 3**). No significant differences in ν1(PO ^3-^) stretching band values between individuals is observed (Mann-Whitney-Wilcoxon statistical test, *p-* value > 0.05), although some skeletal elements of the specimen of *Keilhauia nui* appear to have higher values. The Raman results indicate that there is little mineralogical change in the skeletal elements, except for some bone elements belonging to the *Keilhauia nui*, the *Keilhauia* sp. and the *Colymbosaurus svalbardensis* specimens.

### Ichthyosauria, Plesiosauria and Metriorhynchidae body temperature

Ichthyosauria, Plesiosauria and Metriorhynchidae body temperature estimates have been compared to environmental paleotemperatures which range from 14 ± 2 (Slottsmøya Member, Spitsberg, 65.5°N) to 34 ± 1°C (“Les Ardilles”, France, 34.5°N; **Supplementary Table 2**). Ichthyosauria, Plesiosauria and Metriorhynchidae body temperature estimates are available in **Supplementary Table 2** and reported against corresponding sea water temperature estimates in **Figure 7**.

**Figure 7:**
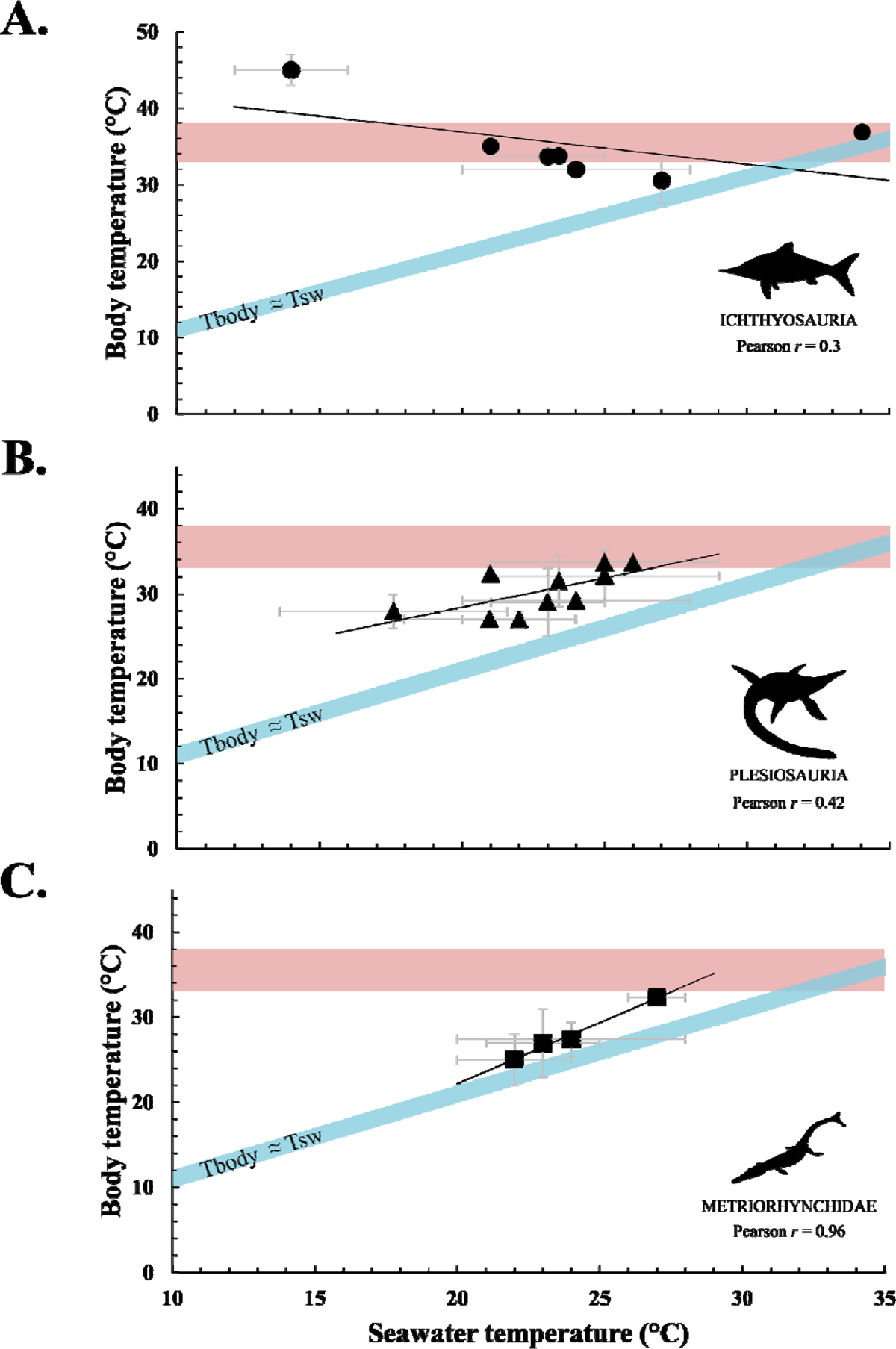
Body temperature estimates of Ichthyosauria (**A.**), Plesiosauria (**B.**) and environment. The body temperature range of extant homeothermic endotherm Cetacea is illustrated in red (Morrison 1962; Hampton et al. 1971; Yeates and Houser 2008), while the body temperature range shown in blue corresponds to the temperature range of a strictly ectothermic poikilothermic model organism whose body temperature is equal to the environmental temperature +2°C. Body and environmental temperatures have been calculated using the equation of Lécuyer et al. (2013) and considering the latitudinal gradient of δ^18^O_sw_ values of Alberti et al. (2020) and an ^18^O-enrichment of body water relative to environmental water of +0.8 ± 0.9% for Mesozoic marine reptiles (Séon et al. 2023). Silhouettes have been made by Gareth Monger for Ichthyosauria and Metriorhynchidae and T. Michael Keesey for Plesiosauria. Images downloaded from PhyloPic and used under a CCBY 3.0 licence: https://creativecommons.org/licenses/by/3.0/.

Our estimates indicate that ichthyosaurs had body temperatures ranging from 31 to 45°C, close to extant Cetacea (Morrison 1962; Hampton et al. 1971; Yeates & Houser 2008, **Fig. 7A**), and systematically higher than those of the sea water in which they lived (between 3 and 31°C higher; **Supplementary Table 2**). No relationship is observed between Ichthyosauria body temperatures and environmental oceanic temperatures (Pearson correlation test, *r* = 0.3; **Fig. 7A**). Plesiosauria body temperature range from 27 to 34°C, these estimates are, as for Ichthyosauria, higher than those of their environment (between 5 and 11°C; **Supplementary Table 2**) and correlate moderately with environmental temperatures (Pearson correlation test, *r* = 0.42; **Fig. 7B**). The estimated body temperatures of Metriorhynchidae, ranging from 25°C to 32°C, are 3 to 5°C higher than (**Supplementary Table 2**) and strongly correlated with environmental temperatures (Pearson correlation test, *r* = 0.96; **Fig. 7C**).

## Discussion

### Preservation of the biological δ^18^O_p_ values

Before discussing the thermophysiological significance of the δ^18^O_p_ values of bones and teeth hydroxylapatite, pristine preservation of the isotope record needs to be assessed. Biotic and abiotic processes leading to the decomposition, burial and fossilization of living organisms may alter the pristine isotope composition of biogenic hydroxylapatite through processes of secondary precipitation, ion adsorption or dissolution–recrystallization of hydroxylapatite (Kolodny et al. 1996; Blake et al. 1997; Trueman et al. 2003, Zazzo et al. 2004a,b). Although no method can definitely demonstrate whether the original isotope compositions have been kept, several ways to assess the preservation state of the oxygen isotope record have been proposed (Iacumin et al. 1996; Kolodny et al. 1996; Fricke et al. 1998; Lécuyer et al. 2003; Pucéat et al. 2004, Zazzo et al. 2004a,*b*; Tütken et al. 2008; Keenan 2016; Lécuyer and Flandrois 2023; Turner-Walker et al. 2023).

In general, enamel is widely considered as more resistant to diagenetic processes compared to bones (e.g. Lowenstam and Weiner 1989; Sillen and LeGeros 1991; Kohn et al. 1999; Lee-Thorp and Sponheimer 2003; Gehler et al. 2011) since it possesses a high crystallinity index, as it is composed of 95% bioapatite crystals. In addition, bioapatite crystals are larger and more compact than those in bone (Pasteris et al. 2008). As a result, the space between the crystals is reduced, limiting fluid circulation, interaction with bacterial and microbial organisms and precipitation of secondary minerals (LeGeros 1981; Driessens and Verbeeck 1990). The precipitation of secondary minerals leads to mineralogical changes observable through mineralogical characterization methods such as Raman spectroscopy. According to Thomas et al. (2007, 2011), bones and teeth altered by diagenetic processes have peak widths (FWHM) of ν1(PO_4_^3-^) at half height of less than 9 cm^-1^ and band positions of ν1(PO_4_^3-^) greater than 964.7 cm^-1^. Among the analysed samples, few skeletal elements have parameters that might suggest that they are mineralogically altered (**Fig. 6**) including different type of bones (skull bones, vertebrae and phalanges) from specimen of *Palvennia hoybergeti*, *Keilhauia nui* and *Colymbosaurus svalbardensis*. The sandy appearance of the remains from the specimen of *Keilhauia nui* could explain why so few silver phosphate crystals were recovered at the end of the chemical preparation. Their low P_2_O_5_ contents compared to the other specimens from the same stratigraphical layer and the mineralogical change observed by Raman spectroscopy would indicate the presence of bone alteration (**Table 2, Supplementary Figure 1B**).

To assess the effect of diagenetic processes on the oxygen isotope composition of the bones and the teeth, we used the difference between oxygen isotope compositions of biogenic hydroxylapatite phosphate and corresponding structural carbonate (δ^18^O_c_ – δ^18^O_p_), the carbonate content (CO_3_^2-^%wt) of hydroxylapatite and the carbon isotope composition of biogenic hydroxylapatite structural carbonate (δ^13^C_c_) (**Fig. 8**). Samples that have either δ^18^O_c_ – δ^18^O_p_ differences higher than +14.7‰ (Iacumin et al. 1996; Pellegrini et al. 2011) or a carbonate content higher than +13.4% are considered doubtful regarding a potential diagenetic alteration (Brudevold and Soremark 1967; Vennemann et al. 2001; McElderry et al. 2013; Wingender et al. 2021). All the samples from Ichthyosauria, Plesiosauria and Metriorhynchidae have δ^18^O_c_ – δ^18^O_p_ differences lower than the critical value of +14.7‰.

**Figure 8:**
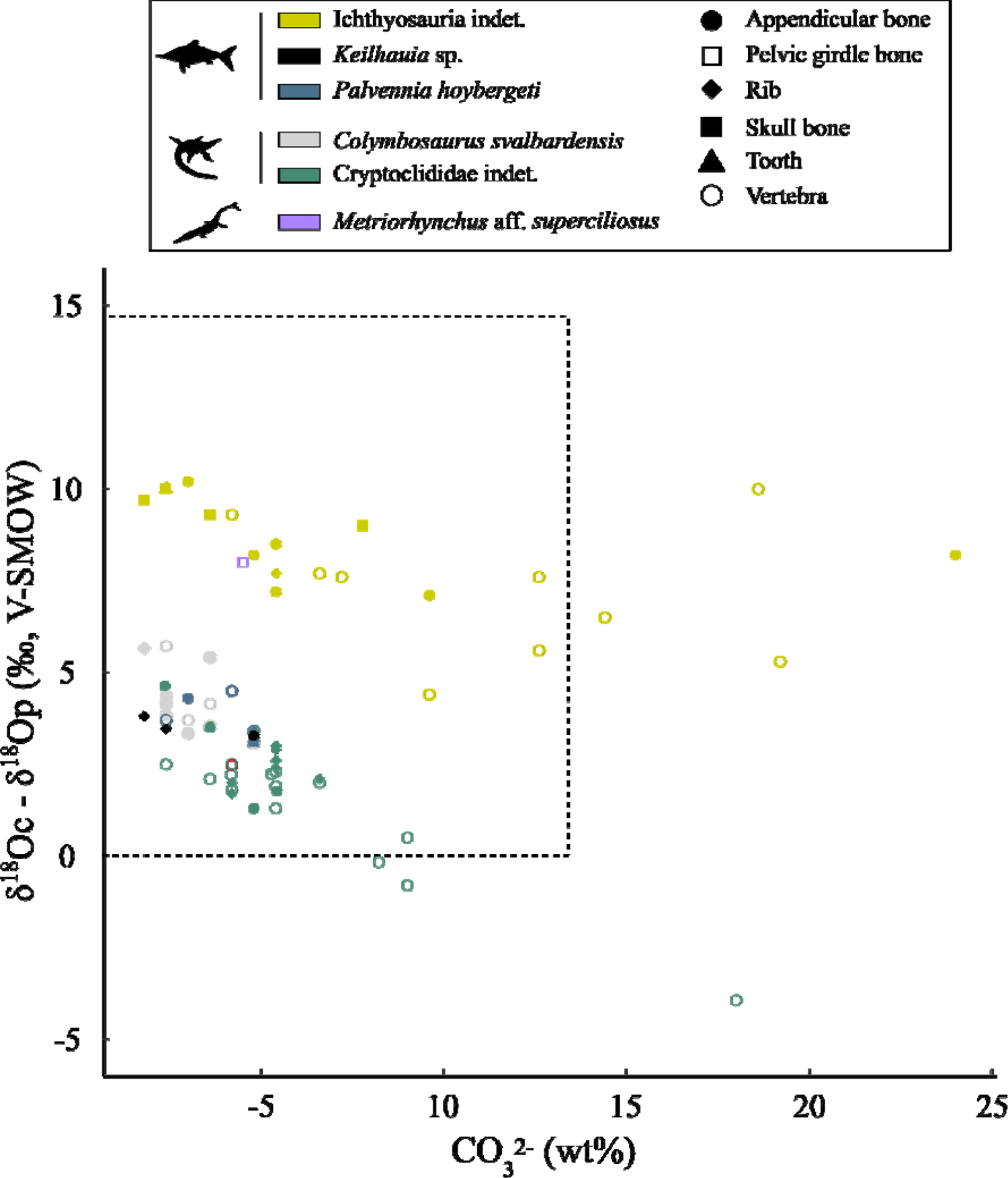
δ^18^O_c_-δ^18^O_p_ differences between teeth and bone types plotted against carbonate content (wt%) of hydroxylapatite. The dashed rectangle zone corresponds to the combination of parameters which would indicate a potential biological preservation of the oxygen isotope composition of the hydroxylapatite of teeth and bones. Silhouettes have been made by Gareth Monger for Ichthyosauria and Metriorhynchidae and T. Michael Keesey for Plesiosauria. Images downloaded from PhyloPic and used under a CCBY 3.0 licence: https://creativecommons.org/licenses/by/3.0/.

Nonetheless, some of them possess carbonate content greater than +13.4% especially from bones belonging to the specimen of Ichthyosauria indet. from “Les Ardilles” (**Fig. 8**), in line with the calcite recrystallizations observed during sampling. Moreover, a trend is observed between carbonate contents and δ^18^O_c_ values considering the tooth δ^18^O_c_ value as the pristine biological signal and that of the calcite recrystallization, which would form the diagenetic endmember (**Supplementary Figure 4**). This relationship also highlights the link between the occurrence of high carbonate content and the ultrastructure (e.g. connected porosity) of the bones, here the vertebrae, since amongst all the skeletal elements analysed in the specimen of Ichthyosauria indet., six (n = 6) are vertebrae and two (n = 2) are limb bones. These skeletal elements are among the bones with the highest porosities within Ichthyosauria (Anderson et al. 2019).

Finally, most of the δ^13^C_c_ values, when plotted against CO_3_ ^2-^ (wt%), conform to two different mixing lines. In the specimen of Cryptoclididae indet. from Spitsbergen, the data point to variable amounts of secondary calcite with δ^13^C_c_ values of −15‰ (**Fig. 9**), which point to a ^13^C-depleted organic carbon source typical of the sulfate reduction zone (Meister and Reyes 2019). In the specimen of Ichthyosauria indet., the data point to variable amounts of more ^13^C-enriched secondary with δ^13^C_c_ values of ∼0‰, in agreement with the value of −0.1‰ measured on intra-bone calcite sampled from the same specimen. These higher isotopic values could either reflect an early precipitation of calcium carbonate close to the sediment-sea water interface with a significant sea water influence or formation below the sulfate-reduction zone close to a ^13^C-enriched methanogenic CO_2_ source (Meister & Reyes 2019). In both specimens, δ^13^C_c_ values point to an almost identical, well-preserved end-member with low CO_3_^2-^ contents (<10%) and δ^13^C_c_ values of −8 to −10‰ V-PDB dominated by structural hydroxylapatite carbonates. The range of δ^13^C_c_ values recorded for these CO_3_ ^2-^-poor samples of marine reptiles are identical to that previously reported for ancient air-breathing vertebrates (Séon et al., 2020), but substantially lower than the δ^13^C_c_ values recorded in the pachycormid Hypscormus sp. teeth measured from “Les Ardilles” and from coeval fossil fishes (−5.47 to +3.15‰ V-PDB; Séon et al., 2020). As expected, air-breathing reptiles have lower δ^13^C_c_ values than fish. In fact, the δ^13^C values of air-breathing reptiles mainly reflect those of their diet, with isotopic fractionation depending on their digestive physiology (Passey et al. 2005). In fish, on the other hand, the δ^13^C values of carbonate come from the diet, but also from the contribution of a large amount of dissolved inorganic carbon in the surrounding water (McConnaughey et al. 1997), which has a high δ^13^C value (Santos et al. 2011). These features demonstrate that, despite the different taphonomical history suggested by the diagenetic δ^13^C_c_ values at the two sites, the geochemical signature of the CO_3_^2-^-poor samples is preserved and hence paleobiologically informative.

**Figure 9:**
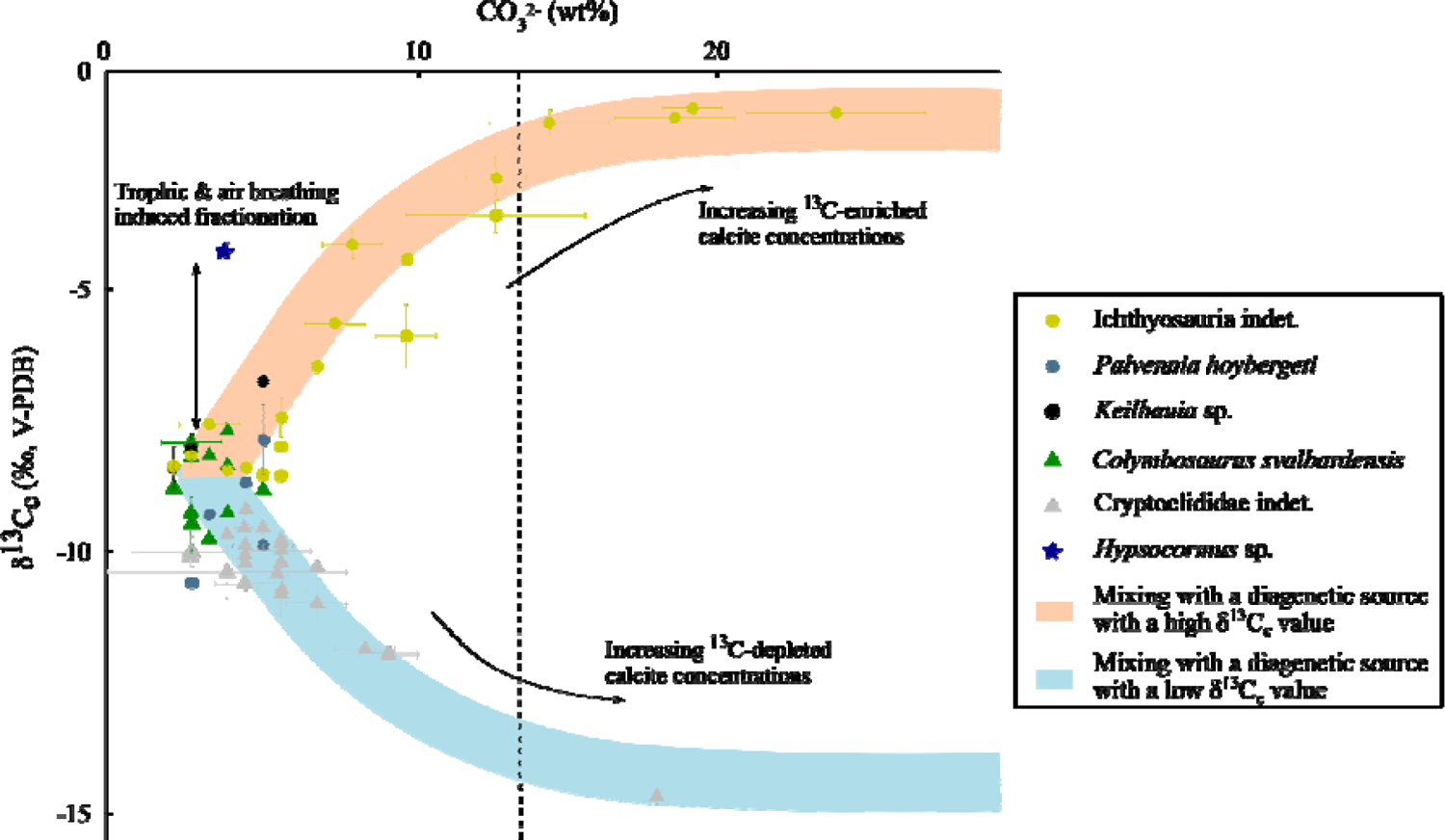
δ^13^C_c_ values (‰, V-PDB) of bone elements and teeth in ichthyosaurs and plesiosaurs as a function of carbonate content (wt%). The values corresponding to the *Hypsocormus* sp. tooth found in the same stratigraphic layer as the Ichthyosauria indet specimen are also plotted. The dotted line marks the boundary between mineralised elements with a carbonate content greater than or less than +13.4%.

From here on and the evidence provided by mineralogical and isotope proxies, we assume that δ^18^O_p_ values from most of the Ichthyosauria, Plesiosauria and Metriorhynchidae samples, except the *Keilhauia nui* specimen (PMO 222.655), have kept at least a significant part of their original oxygen isotope compositions and might be interpreted in terms of thermophysiology. As a precaution, we removed all the skeletal elements for which doubt was expressed regarding the preservation of the original isotopic signal (mineralogical, carbonate content and isotope clues; **Supplementary Table 1**).

### Intra-skeletal body temperature distribution in Ichthyosauria, Plesiosauria and Metriorhynchidae

The oxygen isotope composition of air-breathing aquatic tetrapods is controlled by their body temperatures, the proportion and isotopic composition of metabolic and drinking waters, and the breathing-induced isotope enrichment factor relative to these sources of waters. The possible occurrence of salt glands in at least some taxa of Metriorhynchidae (Fernández and Gasparini 2000, 2008; Gandola et al. 2006; Cowgill et al. 2023), Ichthyosauria (Wahl 2012; Campos et al. 2020; Massare et al. 2021) and Plesiosauria (Buchy et al. 2006; O’Gorman and Gasparini 2013; Páramo-Fonseca et al. 2019) indicates that they would have been able to drink sea water to maintain their water balance. We thus consider that sea water was the main source of water and thus oxygen, as it is the case in extant reptiles with salt glands (Rash and Lillywhite 2019). Since the δ^18^O_sw_ value influencing δ^18^O_p_ values can vary over time and specific regions, the timing of oxygen incorporation into the bones and teeth hydroxylapatite need to be identified to accurately compare the δ^18^O_p_ values of different skeletal elements and identify potential regional heterothermies.

As most reptiles, Ichthyosauria, Plesiosauria and Metriorhynchidae continuously replace their teeth. According to the studies conducted by Kear et al. (2017) and Maxwell et al. (Maxwell et al. 2011a, *b*, 2012), the mineralisation time of the teeth was estimated to be at most 2 to 3 years for plesiosaurs and approximately one year for ichthyosaurs. Although there is currently no available information regarding the tooth renewal time of Metriorhynchidae, it can be assumed that it was comparable to that of extant crocodylomorphs, which is approximately three to fifteen months (Erickson 1996; Finger Jr et al. 2019). In Ichthyosauria and Plesiosauria, bones underwent regular remodelling throughout life, as evidenced by studies conducted by Kolb et al. (2011), Delsett & Hurum (2012), Houssaye et al. (2014), and O’Keefe et al. (2019). For the Metriorhynchidae, cyclical growth marks in some bones indicate no or little bone remodelling (Hua and de Buffrénil 1996), thus the time recorded in the bones is likely to be slightly longer. The δ^18^O_p_ values of the teeth and bones of Ichthyosauria, Plesiosauria, and Metriorhynchidae can be compared since these tissues record a period ranging from one to several years. In addition, annual and seasonal variations in modern δ^18^O_sw_ values of the open ocean are typically only a few tenths of a per mil (∼ ± 0.5 ‰; LeGrande and Schmidt 2006), this alone cannot fully explain the observed δ^18^O_p_ variations in ichthyosaurs and plesiosaurs of more than 1 to 3‰. Thus, we postulate that the δ^18^O_p_ variability recorded in bones and teeth from the Ichthyosauria, Plesiosauria and Metriorhynchidae studied mostly reflects temperature variations.

Assuming that the temporal variation in the oxygen isotope composition of the body water of Ichthyosauria, Plesiosauria and Metriorhynchidae is approximately constant over time, we can infer the range of mineralisation temperatures across their body using the δ^18^O_p_ values of the geochemically best-preserved skeletal elements. Thus, using the equation of Lécuyer et al. (2013), we obtain a mineralisation temperature range of approximately 8 to 12°C for Ichthyosauria (**Fig. 1**), 4 to 12°C for Plesiosauria (**Fig. 2**), and of the order of 4 to 5°C for *Metriorhynchus* aff. *superciliosus* (**Fig. 3**). In comparison, the mineralisation temperature range of skeletal elements estimated from bones δ^18^O_p_ is around 8 to 12°C for extant homeothermic endotherms Cetacea and Pinnipedia, 10°C for poikilothermic endotherm Atlantic bluefin tuna and 15°C for poikilothermic endotherm swordfish (Séon et al., 2022; 2024), while direct measurement of body temperature range for poikilothermic ectotherms such as crocodiles and turtles has been estimated to be ∼10°C (Dunham et al. 1989; Markwick 1998). Note that for specimens of Elasmosauridae indet. and *Metriorhynchus* aff. *superciliosus* we were unable to analyse distal skeletal elements belonging to the appendicular skeleton (not preserved), which may explain why the mineralisation temperature ranges for these specimens are narrower than in the other analysed taxa (**Fig. 2** and **Fig. 3**). For all specimens, the comparison between the different skeletal regions do not allow us, to identify heterothermies in Mesozoic marine reptiles. Indeed, we did not observe as expected colder limbs and tails, and differences in neck temperatures in Plesiosauria. The comparison between the δ^18^O_p_ values of the dorsal region (vertebrae, ribs, girdle bones) and the limbs shows no significant differences except for the specimen of *Colymbosaurus svalbardensis* (**Fig. 4**). However, the differences observed are at the contrary to what we would have expected. This would indicate a higher average temperature of mineralisation of the hindlimbs compared to the skeletal elements of the trunk (vertebrae, ribs, pectoral and pelvic bones). In the specimen of *Colymbosaurus svalbardensis* (PMO 222.663), skeletal elements of the anterior region (dorsal vertebrae, ribs, left and right forelimbs) have significantly higher values than those of the posterior region (left and right hind limbs and caudal vertebrae). These significant differences in δ^18^O_p_ appear unusual in terms of thermoregulatory strategies and could be the result of a preservation difference between the more weathered anterior and posterior skeletal elements of the specimen (Delsett et al., 2016). Regarding the limbs of Plesiosauria, the osteo-histological study carried out by Delsett & Hurum (2012) on two specimens (PMO 216.838, PMO 216.377) of *Colymbosaurus svalbardensis* indicates that the most distal limb bone elements display growth striations perhaps resulting from cyclic growth. In our study, our data reveal no clear relationship between δ^18^O_p_ values and the position of the skeletal element within the limb (**Fig. 5A**). Thus, no body temperature gradient can be identified in the limb of Plesiosauria based on the geochemical approach. An identical conclusion can be drawn regarding the possible presence of a temperature gradient in the neck of Plesiosauria since our detailed mapping of the cervical region shows no relationship between δ^18^O_p_ and vertebral position (**Fig. 5B**). Finally, the only systematic trend recorded by our dataset concerns the teeth that have higher δ^18^O_p_ values than the bones, including skull elements located close to the analysed teeth in specimen PMO 222.667 of *Keilhauia* sp. and *Palvennia hoybergeti* specimen PMO 222.669 (**Fig. 4**). This observation could be explained by a difference in mineralisation temperature at the rostrum of about 3 to 4 °C, but the lower number of values available for each of these observation series means that we must be cautious about such considerations. Indeed, the observed higher values from the teeth could also result from differences in isotopic preservation between teeth enamel and bones.

### Reassement of Ichthyosauria, Plesiosauria and Metriorhynchidae body temperature and thermoregulatory strategies

Previous isotope-based reconstructions of extinct marine reptile body temperatures opted for an ^18^O-enrichment of body water relative to drinking water of +2‰ (Bernard et al. 2010; Séon et al. 2020; Leuzinger et al. 2023), originally estimated by Amiot et al. (2007) from body fluids of young Nile crocodiles (*Crocodylus niloticus*), and Barrick et al., (1999) for turtles (*Chrysemys* sp.). These species have a semi-aquatic ecology, low body mass (1.2 kg to 5 kg) and relatively low metabolic rates. We therefore consider that a +2‰ enrichment of body water relative to environmental water may not be the most appropriate value for Mesozoic fully marine reptiles. We considered instead the +0.8 ± 0.9‰ enrichment value measured on fully aquatic air-breathing vertebrates (*Orcinus orca* and *Tursiops truncatus*; 150 to 3,600 kg) proposed by Séon et al. (2023) which appears more appropriate considering the ecology, the size and the putative thermos-metabolism of the Ichthyosauria, Plesiosauria and Metriorhynchidae studied here.

To reassess Ichthyosauria, Plesiosauria and Metriorhynchidae body temperatures, we have favoured the use of tooth enamel, as its robustness against diagenetic alteration has been demonstrated in previous studies (Lowenstam & Weiner 1989; Sillen & LeGeros 1991; Kohn et al. 1999; Lee-Thorp & Sponheimer 2003; Gehler et al. 2011). A last important change in our work relative to previous studies is the use of a spatially variable δ^18^O_sw_ values calculated using the hemispheric gradient proposed by Alberti et al. (2020). This gradient places δ^18^O_sw_ highest values in subtropical areas subject to high evaporation and lowest values in high-latitudes seas receiving high fluvial input from surrounding large continental areas, consistent with both modelling (Zhou et al. 2008) and empirical evidence (Letulle et al. 2022). We thus consider this scenario as far more realistic than the δ^18^O_sw_ hypothesis used previously (Bernard et al., 2010; Séon et al. 2020). Together, the use of revised values and spatial changes in δ^18^O_sw_ values produce new body temperature estimates of Ichthyosauria, Plesiosauria and Metriorhynchidae that are 3 to 6°C lower than estimated previously (Bernard et al. 2010; Séon et al. 2020, **Supplementary Table 2 and Supplementary Figure 5**).

According to our new estimates, Ichthyosauria maintained their body temperatures at a constant value well above environmental values (**Fig. 7A**) and can be considered as homeothermic endotherms, consistent with previous isotope-based studies (Bernard et al., 2010, Séon et al., 2022). Nevertheless, the body temperature values estimated (45 ± 2°C) for the ichthyosaur of the Slottsmøya Member (*Palvennia hoybergeti* and *Keilhauia* sp.) are higher than those of present-day homeothermic endothermic marine vertebrates, based on a δ^18^O_sw_ value of −1.5‰ from the equation of Alberti et al. (2020). However, this equation is calibrated between 0 and 40°N and it is probable that this δ^18^O_sw_ value of −1.5‰ is overestimated, given that ocean basins such as the Arctic Ocean at the time were landlocked and surrounded by land, and were therefore likely to be subject to freshwater input from precipitation runoff. Estimates from numerical modelling for the middle Cretaceous indicates δ^18^O_sw_ values of −5 ± 2‰ for this region (Zhou et al., 2008). By using this modelled δ^18^O_sw_ value, the estimated body temperature of ichthyosaurs from the Slottsmøya Member would have been 31 ± 4°C, confirming the homeothermic and endothermic nature of Ichthyosauria. At variance with previous studies, however, our results indicate that Plesiosauria, as well as Metriorhynchidae, had body temperature higher than, but covarying with environmental temperatures, and hence should be considered as poikilothermic endotherms. We note that the estimated Metriorhynchidae body temperatures are closer to those of their environment values than Plesiosauria (**Fig. 7B** and **7C**). These thermoregulatory strategies are largely consistent with other indices linked to paleobiogeography, morphology, locomotion and estimated metabolic rates.

Indeed, Ichthyosauria would have been able to live in high latitude environments (Delsett et al. 2016; Rogov et al. 2019) where ocean paleotemperatures were too low for ectothermic turtles and crocodylomorphs to survive, as evidenced by their absence in high latitude deposits from the same stratigraphic levels (Rich et al. 2002; Bardet et al. 2014). This would have been greatly facilitated by their high metabolic rates (de Buffrénil and Mazin 1990; Nakajima et al. 2014; Anderson et al. 2019) and their fusiform morphology that is very effective in limiting heat loss towards the environment (Innes et al. 1990; Gearty et al. 2018) coupled with a subcutaneous layer of adipose tissue reducing thermal conductivity (Lindgren et al. 2018; Delsett et al. 2022). Nevertheless, this could only be the case for the most recent ichthyosaurs in the evolution of the lineage, such as parvipelvian and thunniform ichthyosaurs of the Ophthalmosauridae family. Indeed, primitive Ichthyosauria had more anguiliform morphologies and as far as we know no adipose tissue has been found in these specimens (Motani 2005; Moon and Stubbs 2020).

For Plesiosauria, the characterisation of their thermoregulatory strategy is more ambiguous, and our results interestingly differ from those for ichthyosaurs and to some extent contrast with previous studies (Bernard et al. 2010; Séon et al. 2020; Leuzinger et al. 2023). High metabolic rates close to that of modern endotherms are supported by their paleobiogeographic distribution (Bardet et al. 2014; Delsett et al. 2016; Zverkov et al. 2021), osteo-histological evidence (Wiffen et al. 1995, Kear 2006b; Delsett and Hurum 2012; Fleischle et al. 2018; O’Keefe et al. 2019; Sander and Wintrich 2021) and the identification and quantification of compounds resulting from the degradation of lipids and carbohydrates by infrared spectroscopy (Wiemann et al. 2022). So far, no evidence for a subcutaneous layer of adipose tissue has been reported in Plesiosauria, but this might be a sampling artefact since specimens exhibiting soft-tissue preservation are exceptionally rare (Vincent et al. 2017). On the other hand, our new data show that their body temperature covaried to some extend with environmental temperatures. One possible explanation that may reconcile these apparently contradictory observations is that Plesiosauria produced heat through their locomotor muscles. This heat production strategy is indeed used by sea turtles with the same type of locomotion and enable them to have core body temperatures higher than the rest of the body (Standora et al. 1982; Sato et al. 1994). The heat generated by muscle contraction may then partly conserved by the inertia of their body mass, as in the case of leatherback turtles (Paladino et al. 1990; Sato 2014) whose bone histology studies show evidence of elevated growth rate and high metabolic rate (Nakajima et al. 2014; Wilson 2023). The existence of a comparable heat production system in Plesiosauria may thus partly explain the recorded difference between their body temperature and their environment as well as their inability to maintain their body temperature at a constant level in the thermally least favourable environments (high latitudes). Metriorhynchidae differed from the previous two in terms of their geographical distribution, which was restricted to tropical zones (Bardet et al. 2014) and show osteo-histological features (Hua & de Buffrénil 1996; de Buffrénil et al. 2021) indicative of a poikilothermic ectothermic thermoregulatory strategy. Initially considered as active hunters (Massare 1987; Young and de Andrade 2009; de Andrade et al. 2010; Young et al. 2012), some species could also have been scavengers and opportunists (Hua et al. 2024), a strategy that does not require the production of long and intense efforts, in agreement with the estimates of their movement speed (Massare 1988; Massare et al. 1994; Gutarra et al. 2023). Although their body temperatures have been considered as slightly higher than those of their environment (Séon et al. 2020) our revised estimates point to much lower differences than previously estimated. Interestingly, Gienger et al. (2017) showed that the standard metabolic rate values measured in *Crocodylus porosus* are 36% higher than those of other taxa such as *Alligator mississippiensis* or *Crocodylus johnstoni*. However, this difference in metabolic rate is not visible in the osteo-histological sections taken from these species (de Buffrénil et al. 2021).

Metriorhynchidae may therefore have had a higher metabolic rate, but this difference would not be recorded in bone tissue. We therefore speculate that the production of metabolic heat could have been linked to vital organs such as intestine or liver as well as the locomotor muscles which, as in tuna, would have been internalised and close to the spinal column, and limited heat loss to the surrounding aquatic environment (Graham & Dickson 2004).

## Conclusion

The reassessment of oxygen-isotope-based body temperature estimates of Ichthyosauria, Plesiosauria and Metriorhynchidae have led to a reconsideration of their thermoregulatory strategies. The intra-skeletal δ^18^O_p_ variability in four specimens of Ichthyosauria, three specimens of Plesiosauria and one specimen of *Metriorhynchus* aff. *superciliosus* did not allow us to characterize and locate regional heterothermies. This may be due to the impact of diagenetic processes on the original oxygen isotope composition of bone bioapatite. The newly measured δ^18^O_p_ from the teeth of Ichthyosauria specimens from high paleolatitudes confirm that these organisms had body temperatures higher than those of their living environment and were probably homeothermic endotherms with body temperature ranging between 31 and 37°C. In contrast to Ichthyosauria, it appears that the body temperatures of Plesiosauria and Metriorhynchidae were influenced by changes in environmental temperatures. Therefore, they were most likely poikilothermic endotherms, such as some extant sea turtles and tunas. Nevertheless, the thermoregulatory strategy of Metriorhynchidae remains difficult to characterise because their body temperature estimates are very close to those of their environment and could be defined as either poikilothermic endotherms or poikilothermic ectotherms. Further investigations are needed to clearly define their thermoregulatory strategy. Finally, the independency of Ichthyosauria body temperatures relative to environmental temperatures, confirmed by our study, places these organisms as promising tracers of spatiotemporal changes in δ^18^O_sw_ values that could allow to improve Mesozoic paleoclimate reconstructions.

## Acknowledgments

Stable isotope measurements have been performed at the Plateforme d’Ecologie Isotopique du Laboratoire d’Ecologie des Hydrosystèmes Naturels et Anthropisés (LEHNA, UMR5023, Université Claude Bernard Lyon 1, Lyon, France) and Raman spectroscopy analyses at the Plateforme de Spectroscopie Raman du Laboratoire de Géologie de Lyon (LGL-TPE, UMR5276, Université Claude Bernard Lyon 1). The authors warmly thank the heads and the people responsible of the collection of the Natural History Museum of Oslo, Norway; Michèle Fouché and Sophie Rajaofera from the Muséum d’Histoire Naturelle d’Auxerre, France; Nicolas Morel from the Muséum d’Histoire Naturelle Le Mans and Laurent Picot from the Paléospace museum for accepting to sample specimens in their respective collections. This study was founded by the ANR-OXYMORE (grant no. ANR-18-CE31-0020) and the project 335111 of the Norwegian research council.

## Author Contributions

NS, PV, GS and RA conceived and designed the study. Material preparation and data collection were performed by NS, LLD, AJR and EP. Isotope analysis were performed by NS, RA, FF and CL. Raman spectroscopy data were performed by NS and EP. GS and PV acquired the financial support for the project leading to this publication. PV, SC, CL, GS and RA helped in the supervision of the project. The first draft of the manuscript has been written by NS, RA and PV, and all authors commented on previous versions of the manuscript. All authors read and approved the final manuscript.

## Supplementary information

**Supplementary information 1:** Isotope-based body temperature calculations.

**Table 1:** Summary of information from Ichthyosauria, Plesiosauria, and Metriorhynchidae specimens sampled.

**Table 2:** Synthesis of isotopic values (δ^18^O_p_, δ^18^O_c_, δ^13^C_c_) and CO_3_ ^2-^ and P_2_O_5_ content (wt%) of the bones and teeth hydroxylapatite of the specimens of Ichthyosauria, Plesiosauria and Metriorhynchidae. Abbreviations: ARL = anterior right limb, ALL = anterior left limb, PRL = posterior right limb and PLL = posterior left limb.

## Supplementary Table

**Supplementary Table 1:** Isotopic values (δ^18^O_p_, δ^18^O_c_, δ^13^C_c_) and CO_3_ ^2-^ and P_2_O_5_ content (wt%) of Ichthyosauria, Plesiosauria and Metriorhynchidae bones and teeth phosphate hydroxylapatite. Phosphate oxygen isotope values in bold correspond to data removed from the dataset because of mineralogical alteration and values in italic correspond to data removed from the dataset because of carbonate content (wt%) greater than +13.4%.

**Supplementary Table 2:** Sea water temperature and Ichthyosauria, Plesiosauria and Metriorhynchidae new body temperatures using the equation of Lécuyer et al. (2013) and considering the latitudinal gradient of δ^18^O_sw_ values of Alberti et al. (2020) and an ^18^O-enrichment of body water relative to environmental water of 0.8% for Mesozoic marine reptiles (Séon et al. 2023). Compilation of oxygen isotope compositions of Ichthyosauria, Plesiosauria and Metriorhynchidae teeth phosphate hydroxylapatite according to their location. Difference between old (Anderson et al. 1994, Bernard et al. 2010, Séon et al. 2020) and new body temperature estimates for Ichthyosauria, Plesiosauria and Metriorhynchidae according to palaeolatitude.

**Supplementary Table 3:** Raman spectral parameters of Ichthyosauria, Plesiosauria and Metriorhynchidae bones and teeth samples.

## Supplementary Figures

**Supplementary Figure 1:**
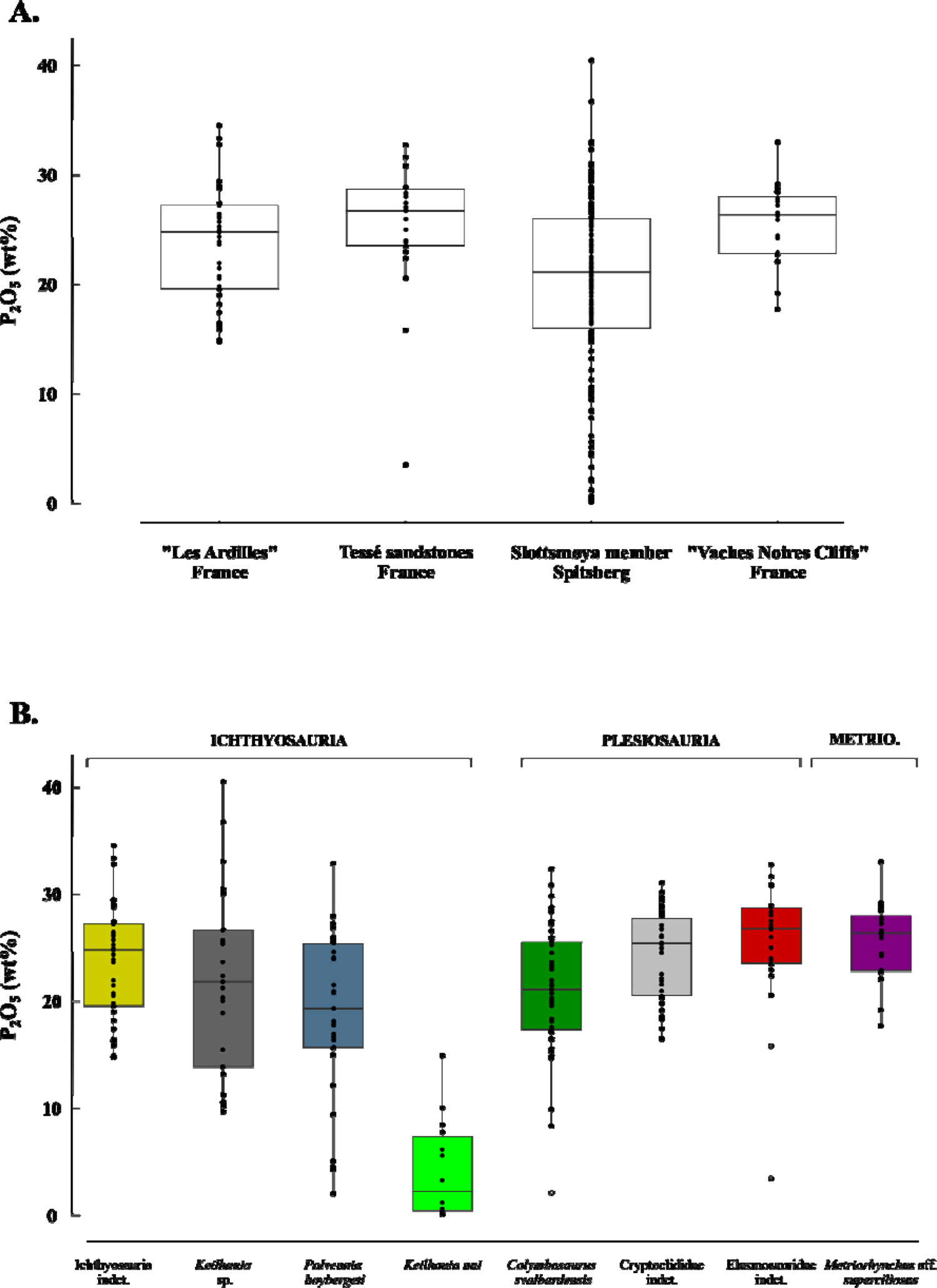
Boxplots showing the P_2_O_5_ content expressed in weight percentage of apatite (wt%) distribution of Mesozoic marine reptile bones and teeth along the fossil deposits (**A.**) and the specimens (**B.**). Outliers are plotted as white circles. The horizontal bars in the boxes correspond to the medians and the whiskers to the minimum and maximum values.

**Supplementary Figure 2:**
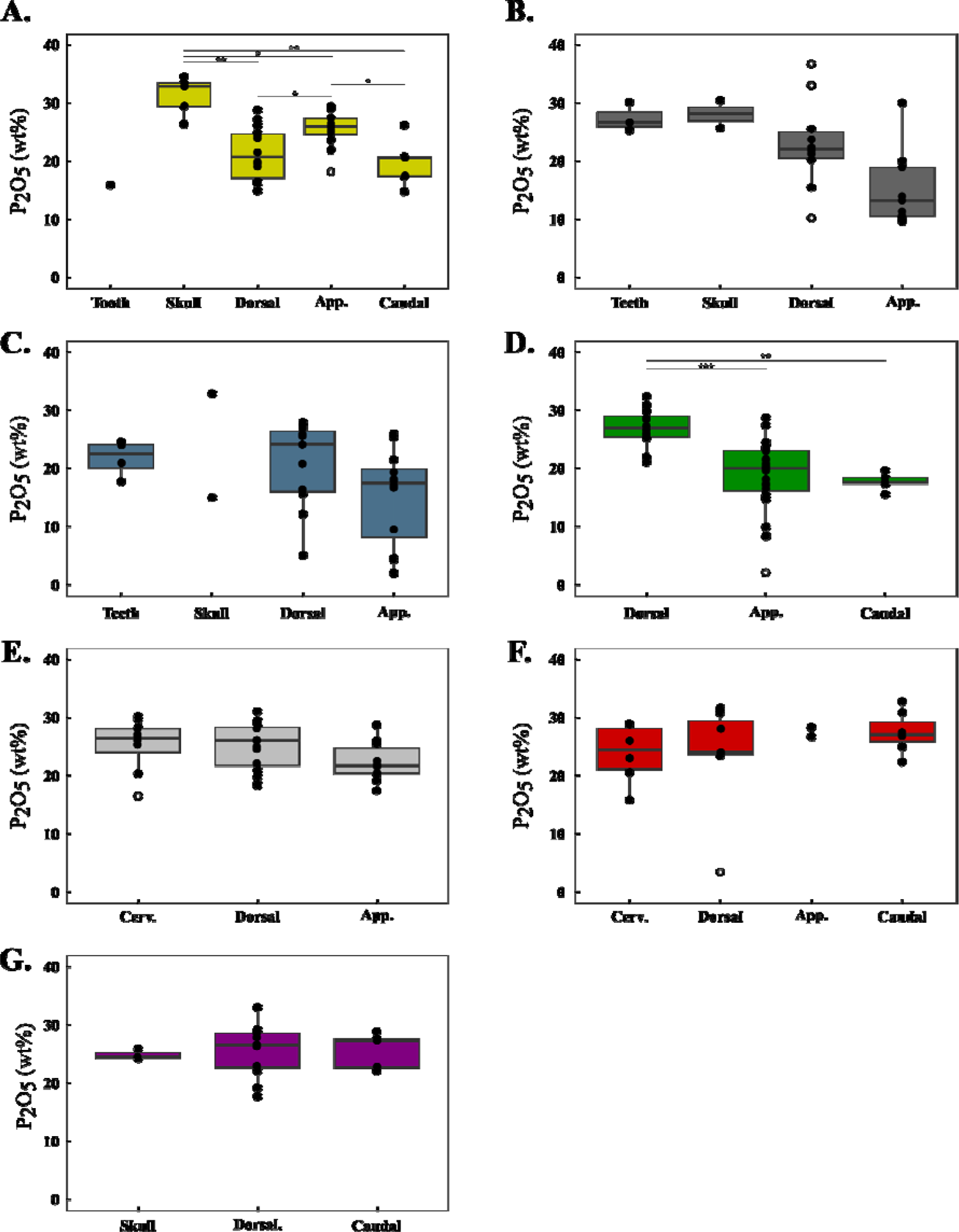
Boxplots showing the P_2_O_5_ content (wt%) distribution of bones sets for the specimens of Ichthyosauria (**A.** Ichthyosauria indet, **B.** *Keilhauia* sp., **C.** *Palvennia hoybergeti*), Plesiosauria (**D.** *Colymbosaurus svalbardensis* and **E.** Cryptoclididae indet., **F.** Elasmosauridae indet.,) and *Metriorhynchus* aff. *superciliosus* (**G.**). Asterisks indicate the significance of the observed differences between pair of groups: * for *p*-value < 0.05, ** for *p*-value < 0.01 and *** for *p*-value *<*0.001. Outliers are plotted as white circles. The horizontal bars in the boxes correspond to the medians and the whiskers to the minimum and maximum values.

**Supplementary Figure 3:**
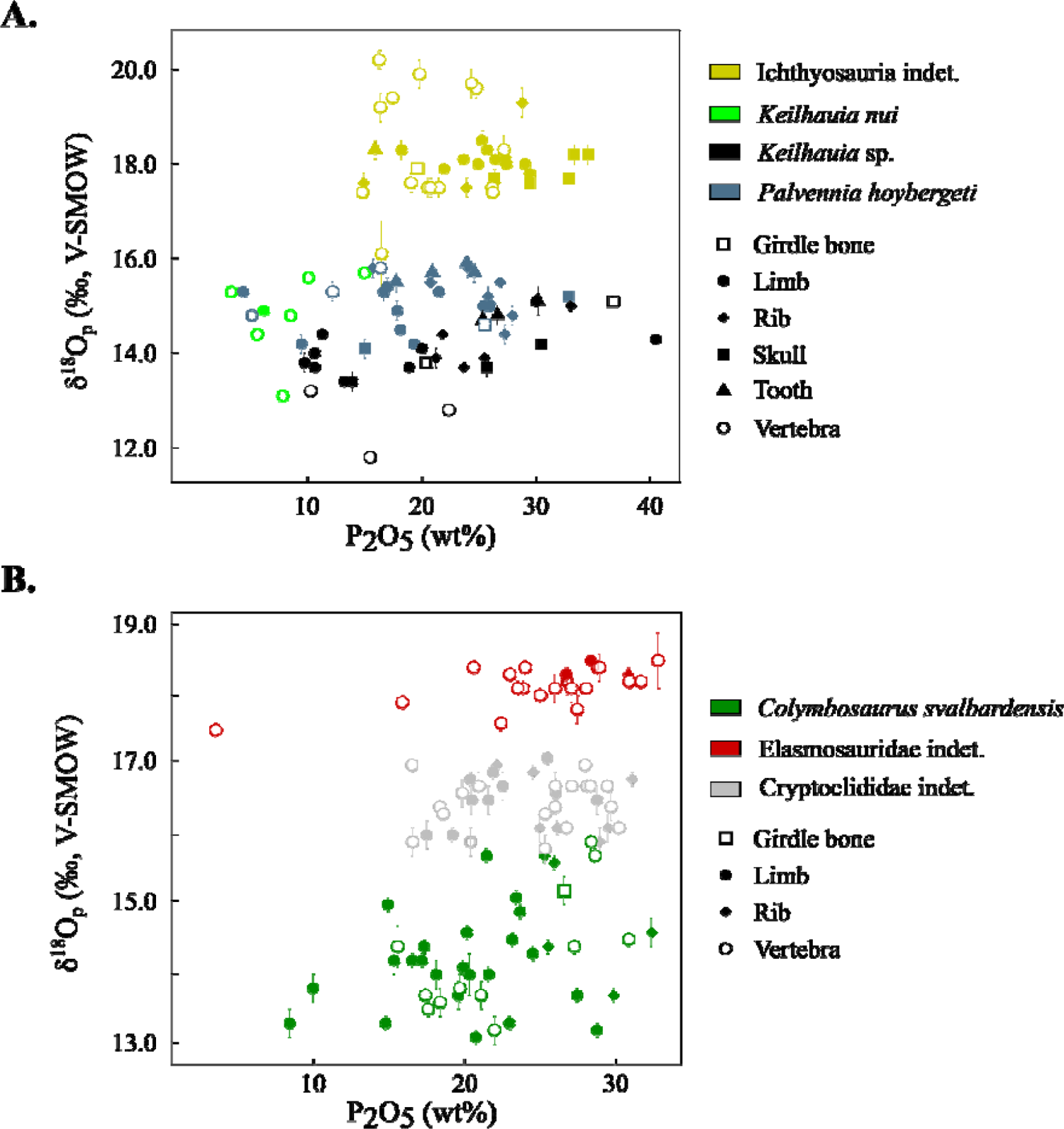
Values of δ^18^O_p_ (‰, V-SMOW) as a function of P_2_O_5_ content (wt%) according to mineralized elements (bones and teeth) in Ichthyosauria (**A.**) and Plesiosauria (**B.**).

**Supplementary Figure 4:**
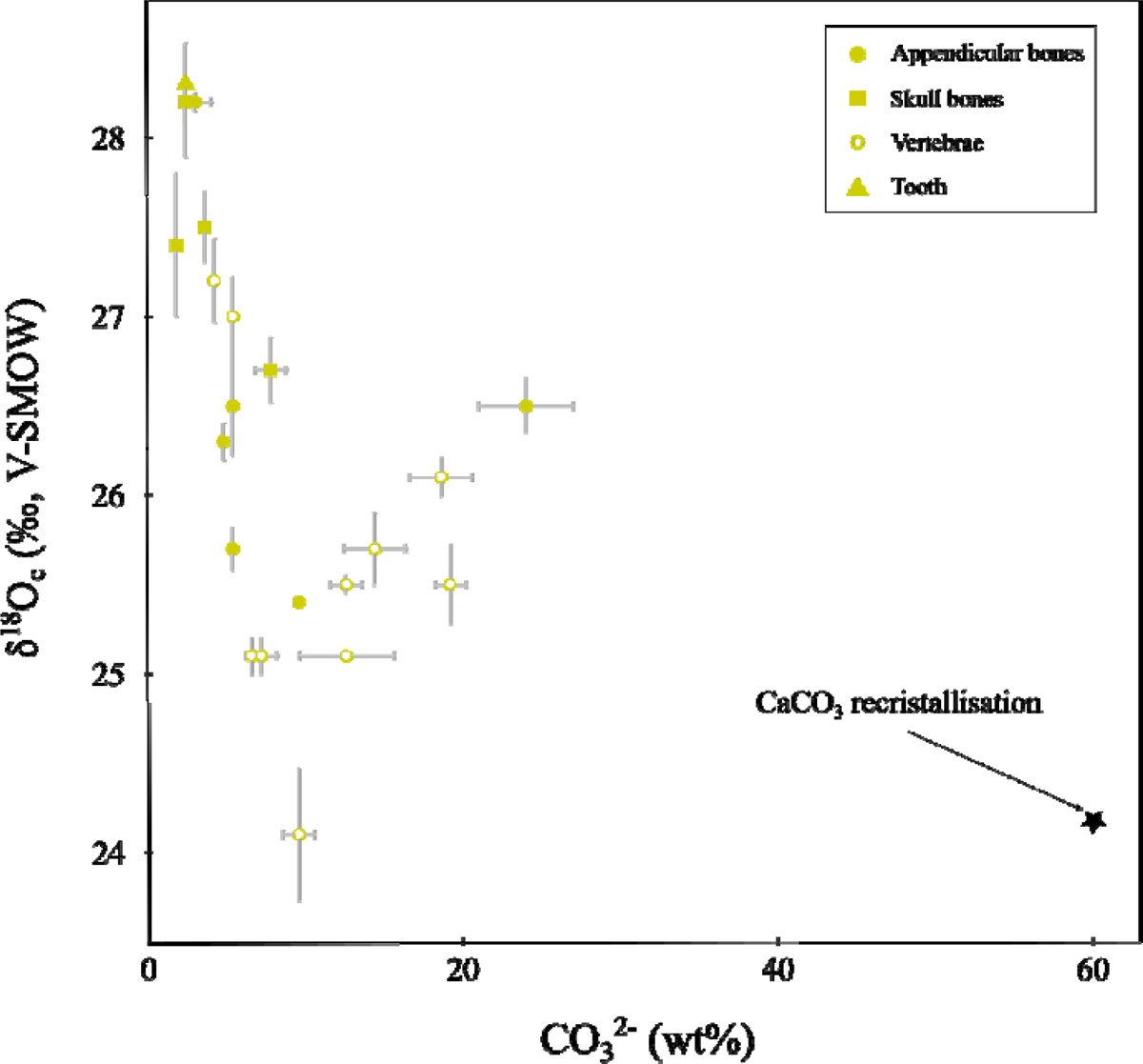
δ^18^O_c_ values (‰, V-SMOW) versus carbonate content of the bones and tooth from the specimen of Ichthyosauria indet.

**Supplementary Figure 5:**
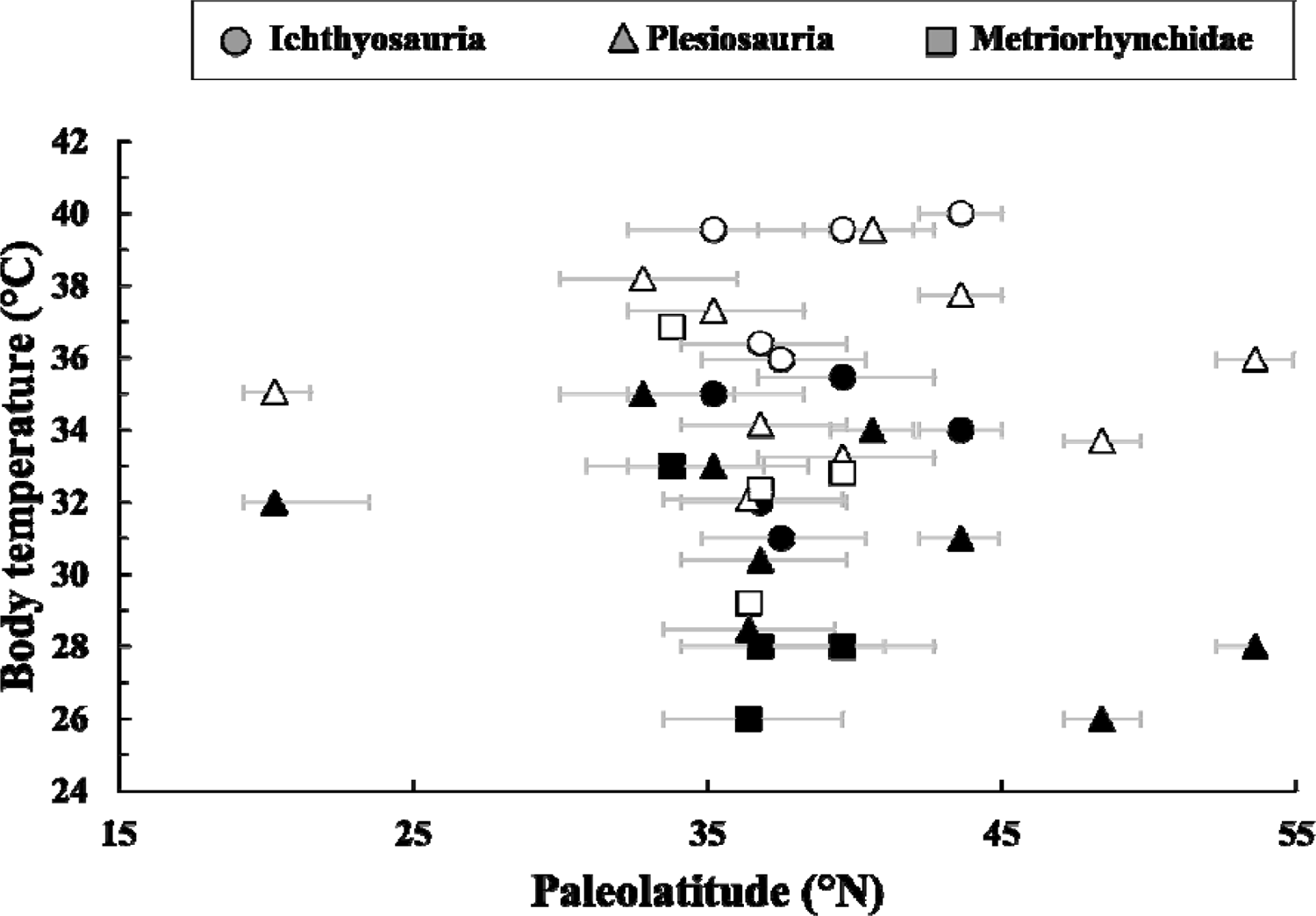
Comparison between previous body temperature in white (Bernard et al. 2010; Séon et al. 2020) and new body temperature estimates in black for Ichthyosauria, Plesiosauria and Metriorhynchidae according to palaeolatitude.

