## Supplementary material for "Reassessment of body temperature and thermoregulation strategies in Mesozoic marine reptiles": Table 1

| | Collection number | Museal institution | Taxonomy | Material | Estimated size (m) | Epoch | Stage | Locality | References |
|---|---|---|---|---|---|---|---|---|---|
| **Ichthyosauria** | - | Musée d'Histoire naturelle d'Auxerre (France) | Ichthyosauria indet. | bones (36) and tooth (1) | 6 to 8 | Late Jurassic | Kimmeridgian | "Les Ardilles", Auxerre, France | Mazin and Pavy (1995) |
| | PMO 222.655 | Natural History Museum of Oslo (Norway) | *Keilhauia nui* | bones (24) | 2 to 3 | Lower Cretaceous | Ryazanian | Slottsmøya Member, Spitsberg, Svalbard, Norway | Delsett et al. (2017) |
| | PMO 222.667 | Natural History Museum of Oslo (Norway) | *Keilhauia* sp. | bones (22) and teeth (3) | ? | Late Jurassic | Volgian | Slottsmøya Member, Spitsberg, Svalbard, Norway | Delsett et al. (2019) |
| | PMO 222.669 | Natural History Museum of Oslo (Norway) | *Palvennia hoybergeti* | bones (25) and teeth (4) | ? | Late Jurassic | Volgian | Slottsmøya Member, Spitsberg, Svalbard, Norway | Delsett et al. (2018) |
| **Plesiosauria** | PMO 212.662 | Natural History Museum of Oslo (Norway) | Cryptoclididae indet. | bones (38) | ? | Late Jurassic | Volgian | Slottsmøya Member, Spitsberg, Svalbard, Norway | Under study |
| | PMO 222.663 | Natural History Museum of Oslo (Norway) | *Colymbosaurus svalbardensis* | bones (41) | ? | Late Jurassic | Volgian | Slottsmøya Member, Spitsberg, Svalbard, Norway | Roberts et al. (2017) |
| | MHNLM.2005.16.1 | Musée d'histoire Naturelle Le Mans (France) | Elasmosauridae indet. | bones (22) | 3.5 - 4 | Middle Jurassic | Aalenian | Tessé Sandstones, Saint-Rémy du Val commune, Sarthe Department, France | Vincent et al. (2007) |
| **Metriorhynchidae** | MPV 2010.3.610 | Paléospace Museum (France) | *Metriorhynchus* aff. *superciliosus* | bones (20) | 2 | Middle Jurassic | Late Callovian | Marnes de Dives, "Vaches Noires Cliffs", Villers-sur-Mer, France | Le Mort et al. (2022) |
| | - | Paléospace Museum (France) | Metriorhynchidae indet. | teeth (2) from probably 2 specimens | ? | Middle Jurassic | Late Callovian | Marnes de Dives, "Vaches Noires Cliffs", Villers-sur-Mer, France | This study |
