## Supplementary material for "Reassessment of body temperature and thermoregulation strategies in Mesozoic marine reptiles": Table 2

| | | $\delta^{18}O_p$ (‰, V-SMOW) | | | | $\delta^{18}O_c$ (‰, V-SMOW) | | | | $\delta^{18}O_c$ (‰, V-PDB) | | | | $\delta^{13}C_c$ (‰, V-PDB) | | | | $CO_3^{2-}$ (wt%) | | | | $P_2O_5$ (wt%) | | | |
|---|---|---|---|---|---|---|---|---|---|---|---|---|---|---|---|---|---|---|---|---|---|---|---|---|---|
| | | **Range** | **Mean** | **SEM** | **n** | **Range** | **Mean** | **SEM** | **n** | **Range** | **Mean** | **SEM** | **n** | **Range** | **Mean** | **SEM** | **n** | **Range** | **Mean** | **SEM** | **n** | **Range** | **Mean** | **SEM** | **n** |
| Ichthyosauria indet. | Global | 16.1 to 20.2 | 18.2 | 0.8 | 36 | 24.10 to 28.30 | 26.34 | 1.17 | 21 | -6.58 to -2.56 | -4.45 | 1.13 | 21 | -8.56 to -1.56 | -5.79 | 2.68 | 21 | 2 to 24 | 9 | 6 | 21 | 14.8 to 34.6 | 23.7 | 5.3 | 37 |
| | Teeth | 18.3 | - | - | 1 | 28.3 | | | 1 | -2.56 | - | - | 1 | -8.18 | - | - | 1 | 2 | - | - | 1 | 15.9 | - | - | 1 |
| | Skull | 17.6 to 18.2 | 17.9 | 0.3 | 5 | 26.70 to 28.20 | 27.4 | 0.53 | 4 | -4.08 to -2.61 | -3.36 | 0.52 | 4 | -8.46 to -4.16 | -7.3 | 1.82 | 4 | 2 to 8 | 4 | 2 | 4 | 26.4 to 34.6 | 31.3 | 3 | 5 |
| | Dorsal region | 16.1 to 20.2 | 18.4 | 1.2 | 14 | 24.10 to 27.20 | 25.78 | 0.94 | 8 | -6.58 to -3.62 | -4.99 | 0.89 | 8 | -8.56 to -1.56 | -4.57 | 2.76 | 8 | 4 to 19 | 11 | 5 | 8 | 14.9 to 28.8 | 21.4 | 4.4 | 14 |
| | Caudal region | 17.4 to 19.4 | 17.8 | 0.8 | 5 | 25.1 | 25.1 | 0 | 2 | -5.65 | -5.65 | 0 | 2 | -6.48 to -3.59 | -5.04 | 1.45 | 2 | 7 to 13 | 10 | 3 | 2 | 14.8 to 26.2 | 19.9 | 3.8 | 5 |
| | Appendicular | 17.8 to 18.5 | 18.1 | 0.2 | 11 | 25.4 to 28.2 | 26.43 | 0.89 | 6 | -5.38 to -2.63 | -4.38 | 0.87 | 6 | -8.54 to -1.65 | -6.28 | 2.45 | 6 | 3 to 24 | 9 | 7 | 6 | 18.2 to 29.5 | 25.5 | 3 | 12 |
| *Palvennia hoybergeti* | Global | 14.1 to 15.9 | 15.1 | 0.5 | 27 | 18.20 to 19.10 | 18.62 | 0.41 | 5 | -12.37 to -11.43 | -11.93 | 0.42 | 5 | -10.6 to -7.88 | -9.28 | 1.05 | 5 | 2 to 5 | 4 | 1 | 5 | 2.0 to 32.9 | 18.8 | 7.9 | 29 |
| (PMO 222.669) | Teeth | 15.5 to 15.9 | 15.7 | 0.1 | 4 | - | - | - | - | | | | | | | | | | | | | 17.7 to 24.6 | 21.8 | 2.7 | 4 |
| | Skull | 14.1 to 15.2 | 14.7 | 0.6 | 2 | 18.30 | | | 1 | -12.27 | | | 1 | -9.89 | | | 1 | 8 | | | 1 | 15.0 to 32.9 | 23.9 | 8 | 2 |
| | Dorsal region | 14.4 to 15.8 | 15.2 | 0.5 | 11 | 18.20 to 19.10 | 18.77 | 0.4 | 3 | -12.37 to -11.43 | -11.79 | 0.42 | 3 | -10.60 to -7.88 | -9.06 | 1.14 | 3 | 2 to 5 | 4 | 1 | 3 | 5.1 to 28.0 | 20.7 | 7.1 | 11 |
| | Appendicular | 14.2 to 15.4 | 14.9 | 0.4 | 10 | 18.5 | | | 1 | -12.03 | | | 1 | -9.31 | | | 1 | 5 | | | 1 | 2.0 to 26.0 | 15.2 | 7.8 | 12 |
| *Keilhauia nui* | Global | 13.1 to 15.7 | 14.8 | 0.9 | 7 | | | | | | | | | | | | | | | | | 3.3 to 15.0 | 8 | 3.8 | 7 |
| (PMO 222.655) | Dorsal region | 13.1 to 15.7 | 14.8 | 1.0 | 5 | | | | | | | | | | | | | | | | | 3.3 to 15.0 | 8.3 | 4 | 5 |
| | Caudal region | 14.8 | - | - | 1 | | | | | | | | | | | | | | | | | 8.5 | - | - | 1 |
| | Appendicular | 14.9 | - | - | 1 | | | | | | | | | | | | | | | | | 6.1 | - | - | 1 |
| *Keilhauia* sp. | Global | 11.8 to 15.1 | 14.0 | 0.8 | 25 | 17.51 to 18.37 | 17.92 | 0.35 | 3 | -13.01 to -12.17 | -12.61 | 0.34 | 3 | -8.40 to -6.77 | -7.74 | 0.7 | 3 | 2 to 5 | 3 | 1 | 3 | 9.7 to 40.5 | 21.9 | 8.5 | 25 |
| (PMO 222.667) | Teeth | 14.7 to 15.1 | 14.9 | 0.2 | 3 | | | | | | | | | | | | | | | | | 25.3 to 30.2 | 27.4 | 2.0 | 3 |
| | Skull | 13.7 to 14.2 | 14.0 | 0.3 | 2 | | | | | | | | | | | | | | | | | 25.7 to 30.5 | 28.1 | 2.4 | 2 |
| | Dorsal region | 11.8 to 15.1 | 13.8 | 0.9 | 10 | 17.51 to 17.87 | 17.69 | 0.18 | 2 | -13.01 to -12.66 | -12.84 | 0.18 | 2 | -8.40 to -8.05 | -8.23 | 0.18 | 2 | 3 to 4 | 2 | 0 | 2 | 10.3 to 36.8 | 23 | 7.3 | 10 |
| | Appendicular | 13.4 to 15.1 | 14.0 | 0.5 | 10 | 18.37 | - | - | 1 | -12.17 | - | - | 1 | -6.77 | - | - | 1 | 5 | - | - | 1 | 9.7 to 40.5 | 17.9 | 9.6 | 10 |
| *Colymbosaurus svalbardensis* | Global | 13.1 to 15.9 | 14.3 | 0.8 | 40 | 17.83 to 20.06 | 18.62 | 0.74 | 13 | -12.7 to -10.48 | -11.92 | 0.73 | 13 | -10.01 to -7.71 | -8.76 | 0.75 | 13 | 2 to 5 | 3 | 1 | 13 | 8.4 to 32.4 | 21.5 | 5.5 | 40 |
| (PMO 222.663) | Dorsal region | 13.2 to 15.9 | 14.7 | 0.9 | 12 | 17.84 to 20.06 | 18.7 | 0.9 | 5 | -12.69 to -10.48 | -11.9 | 0.9 | 5 | -10.01 to -8.80 | -9.30 | 0.5 | 5 | 2 to 5 | 3 | 1 | 5 | 21.1 to 32.4 | 27 | 3.4 | 12 |
| | Caudal region | 13.5 to 14.4 | 13.8 | 0.4 | 5 | 19.32 | - | - | 1 | -11.25 | - | - | 1 | -9.27 | - | - | 1 | 2 | - | - | 1 | 15.5 to 19.7 | 17.7 | 1.5 | 5 |
| | ALL | 14.9 | - | - | 1 | 18.44 | - | - | 1 | -12.05 | - | - | 1 | -7.71 | - | - | 1 | 4 | - | - | 1 | 23.7 | - | - | 1 |
| | ARL | 14.2 to 15.7 | 14.9 | 0.6 | 5 | 19.36 | - | - | 1 | -11.16 | - | - | 1 | -8.22 | - | - | 1 | 2 | - | - | 1 | 14.9 to 23.4 | 19.9 | 3.9 | 5 |
| | PLL | 13.1 to 14.2 | 13.5 | 0.4 | 9 | | | | | | | | | | | | | | | | | 8.4 to 28.8 | 18.2 | 6.4 | 9 |
| | PRL | 13.7 to 14.6 | 14.2 | 0.3 | 8 | 17.83 to 19.51 | 18.3 | 0.7 | 5 | -12.70 to -11.07 | -12.20 | 0.7 | 5 | -9.49 to -7.92 | -8.4 | 0.6 | 5 | 2 to 4 | 3 | 1 | 5 | 15.3 to 27.4 | 20.4 | 3.9 | 8 |
| Cryptoclididae indet. | Global | 15.8 to 17.1 | 16.5 | 0.4 | 38 | 12.47 to 20.63 | 18.28 | 1.47 | 27 | -17.83 to -9.92 | -12.25 | 1.42 | 27 | -14.68 to -9.21 | -10.50 | 1.09 | 27 | 2 to 18 | 6 | 3 | 27 | 16.5 to 31.1 | 24.3 | 4.2 | 38 |
| (PMO 212.662) | Cervical region | 15.8 to 17.0 | 16.4 | 0.4 | 12 | 16.2 to 19.2 | 18.1 | 0.9 | 11 | -14.27 to -11.39 | -12.4 | 0.8 | 11 | -11.99 to -9.56 | -10.50 | 0.8 | 11 | 2 to 9 | 5 | 2 | 11 | 16.5 to 30.2 | 25 | 4.7 | 12 |
| | Dorsal region | 15.9 to 17.0 | 16.5 | 0.3 | 16 | 12.47 to 19.1 | 18.07 | 1.9 | 12 | -17.83 to -11.43 | -12.40 | 1.8 | 12 | -14.68 to -9.21 | -10.60 | 1.4 | 12 | 4 to 18 | 6 | 3 | 12 | 18.3 to 31.1 | 25 | 4 | 16 |
| | Appendicular | 16.0 to 17.1 | 16.6 | 0.4 | 10 | 18.00 to 20.63 | 19.3 | 1.3 | 4 | -12.57 to -9.92 | -11.30 | 1.2 | 4 | -10.73 to -9.56 | -10.02 | 0.5 | 4 | 2 to 5 | 4 | 1 | 4 | 17.5 to 28.8 | 22.4 | 3.4 | 10 |
| Elasmosauridae indet. | Global | 17.5 to 18.5 | 18.2 | 0.3 | 22 | | | | | | | | | | | | | | | | | 3.5 to 32.8 | 25.3 | 6.1 | 22 |
| (MHNLM.2005.16.1) | Cervical region | 17.9 to 18.4 | 18.3 | 0.2 | 6 | | | | | | | | | | | | | | | | | 15.8 to 28.9 | 23.8 | 4.7 | 6 |
| | Dorsal region | 17.5 to 18.1 | 18.1 | 0.3 | 7 | | | | | | | | | | | | | | | | | 3.5 to 31.7 | 23.6 | 8.8 | 7 |
| | Caudal region | 17.6 to 18.5 | 18.1 | 0.3 | 7 | | | | | | | | | | | | | | | | | 22.4 to 32.8 | 27.5 | 3.2 | 7 |
| | Appendicular | 18.3 to 18.5 | 18.4 | 0.1 | 2 | | | | | | | | | | | | | | | | | 26.7 to 28.3 | 27.5 | 0.8 | 2 |
| *Metriorhynchus* aff. *superciliosus* | Global | 19.3 to 20.3 | 19.9 | 0.3 | 20 | | | | | | | | | | | | | | | | | 17.7 to 33.0 | 25.6 | 3.7 | 20 |
| (MPV 2010.3.610) | Skull | 19.4 to 19.9 | 19.7 | 0.2 | 3 | | | | | | | | | | | | | | | | | 24.2 to 25.9 | 24.9 | 0.9 | 3 |
| | Dorsal region | 19.3 to 20.3 | 19.9 | 0.3 | 12 | 27.3 | - | - | 1 | -3.53 | - | - | 1 | -8.22 | - | - | 1 | 5 | - | - | 1 | 17.7 to 33.0 | 25.7 | 4.4 | 12 |
| | Caudal region | 19.9 to 20.1 | 20.0 | 0.1 | 5 | | | | | | | | | | | | | | | | | 22.1 to 28.9 | 25.7 | 3.1 | 5 |
