## Supplementary Table 1 for "Reassessment of body temperature and thermoregulation strategies in Mesozoic marine reptiles"

| Taxa | Coll. N° | #Sample | Skeletal element | Additionnal information | Skeletal region | $\delta^{18}O_p$ (‰, V-SMOW) | | | $\delta^{18}O_c$ (‰, V-SMOW) | | | $\delta^{18}O_c$ (‰, V-PDB) | | | $\delta^{13}C_c$ (‰, V-PDB) | | | wt% $CO_3^{2-}$ | | | wt% $P_2O_5$ | |
|---|---|---|---|---|---|---|---|---|---|---|---|---|---|---|---|---|---|---|---|---|---|---|
| | | | | | | Mean | SD | N | Mean | SD | N | Mean | SD | N | Mean | SD | N | Mean | SD | N | | |
| Ichthyosauria indet. | | ICK22 | Left humerus | | Appendicular | 17.8 | 0.2 | 4 | | | | | | | | | | | | | 29.5 | |
| | | ICK23 | Left radius | | Appendicular | 18.1 | 0.1 | 5 | 26.3 | 0.1 | 2 | -4.51 | 0.1 | 2 | -8.54 | 0.04 | 2 | 5 | 0 | 2 | 26.4 | |
| | | ICK24 | Left ulna | | Appendicular | 18.1 | 0.1 | 4 | | | | | | | | | | | | | 23.6 | |
| | | ICK25 | Phalange | Proximal | Appendicular | | | | | | | | | | | | | | | | 26.1 | |
| | | ICK26 | Phalange | Proximal | Appendicular | 18 | 0.1 | 3 | | | | | | | | | | | | | 27.4 | |
| | | ICK27 | Phalange | Intermediate | Appendicular | 17.9 | 0.1 | 4 | | | | | | | | | | | | | 21.9 | |
| | | ICK28 | Phalange | Intermediate | Appendicular | 18 | 0.1 | 5 | 26.5 | 0.28 | 3 | -4.33 | 0.28 | 3 | -8 | 0.05 | 3 | 5 | 0 | 3 | 24.9 | |
| | | ICK29 | Phalange | Distal | Appendicular | 18.1 | 0.1 | 5 | | | | | | | | | | | | | 27.3 | |
| | | ICK30 | Phalange | Distal | Appendicular | 18.5 | 0.2 | 5 | 25.7 | 0.12 | 3 | -5.09 | 0.12 | 3 | -7.45 | 0.37 | 3 | 5 | 0 | 3 | 25.3 | |
| | | ICK31 | Phalange? | Intermediate? | Appendicular | **18.3** | 0.2 | 5 | 26.5 | 0.15 | 3 | -4.31 | 0.15 | 3 | -1.65 | 0.14 | 3 | 24 | 3 | 3 | 18.2 | |
| | | ICK32 | Phalange? | Intermediate? | Appendicular | 18.3 | 0.1 | 5 | 25.4 | - | 1 | -5.38 | - | 1 | -4.43 | - | 1 | 10 | - | 1 | 25.7 | |
| | | ICK33 | Phalange? | Intermediate? | Appendicular | 18 | 0.1 | 4 | 28.2 | 0.05 | 2 | -2.63 | 0.05 | 2 | -7.59 | 0.04 | 2 | 3 | 1 | 2 | 29.0 | |
| | | | | | *Min.* | 17.8 | | | 25.40 | | | -5.38 | | | -8.54 | | | 3 | | | 18.2 | |
| | | | | | *Max.* | 18.5 | | | 28.20 | | | -2.63 | | | -1.65 | | | 24 | | | 29.5 | |
| | | | | | *Mean / SEM* | 18.1 | 0.2 | | 26.43 | 0.89 | | -4.38 | 0.87 | | -6.28 | 2.45 | | 9 | 7 | | 25.5 | 3.0 |
| | 302 | ICK2 | Caudal vertebra | Anterior | Caudal | 19.4 | 0.1 | 3 | | | | | | | | | | | | | 17.4 | |
| | 313 | ICK9 | Caudal vertebra | Anterior | Caudal | 17.5 | 0.2 | 5 | | | | | | | | | | | | | 20.5 | |
| | 341 | ICK13 | Caudal vertebra | | Caudal | 17.5 | 0.2 | 5 | 25.10 | 0.01 | 2 | -5.65 | 0.01 | 2 | -3.59 | 0.32 | 2 | 13 | 3 | 2 | 20.7 | |
| | 343 | ICK14 | Caudal vertebra | More distal than ICK13 | Caudal | 17.4 | 0.1 | 3 | 25.10 | 0.10 | 2 | -5.65 | 0.10 | 2 | -6.48 | 0.01 | 2 | 7 | 0 | 3 | 26.2 | |
| | 344 | ICK15 | Caudal vertebra | More distal than ICK14 | Caudal | 17.4 | 0.1 | 4 | | | | | | | | | | | | | 14.8 | |
| | | | | | *Min.* | 17.4 | | | 25.10 | | | -5.65 | | | -6.48 | | | 7 | | | 14.8 | |
| | | | | | *Max.* | 19.4 | | | 25.10 | | | -5.65 | | | -3.59 | | | 13 | | | 26.2 | |
| | | | | | *Mean / SEM* | 17.84 | 0.8 | | 25.10 | 0.00 | | -5.65 | 0.00 | | -5.04 | 1.45 | | 10 | 3 | | 19.9 | 3.8 |
| | 299 | ICK1 | Dorsal vertebra | Posterior | Dorsal | 18.3 | 0.3 | 3 | | | | | | | | | | | | | 27.2 | |
| | 305 | ICK3 | Dorsal vertebra | Anterior | Dorsal | 19.6 | 0.2 | 4 | | | | | | | | | | | | | 24.8 | |
| | 307 | ICK4 | Dorsal vertebra | Posterior | Dorsal | **19.2** | 0.3 | 3 | 25.7 | 0.2 | 3 | -5.03 | 0.2 | 3 | -1.84 | 0.26 | 3 | 14 | 2 | 3 | 16.4 | |
| | 310 | ICK5 | Dorsal vertebra | Anterior | Dorsal | 19.7 | 0.3 | 4 | 24.1 | 0.37 | 3 | -6.58 | 0.37 | 3 | -5.89 | 0.6 | 3 | 10 | 1 | 3 | 24.4 | |
| | 311 | ICK6 | Dorsal vertebra | Anterior | Dorsal | 19.9 | 0.3 | 4 | 25.5 | 0.05 | 3 | -5.22 | 0.05 | 3 | -2.89 | 0.42 | 3 | 13 | 1 | 3 | 19.8 | |
| | 312 | ICK7 | Dorsal vertebra | Mid-dorsale | Dorsal | **20.2** | 0.2 | 4 | 25.5 | 0.22 | 3 | -5.24 | 0.22 | 3 | -1.56 | 0.04 | 3 | 19 | 1 | 3 | 16.3 | |
| | 315 | ICK8 | Dorsal vertebra | Mid-dorsale | Dorsal | **16.1** | 0.7 | 4 | 26.1 | 0.11 | 2 | -4.72 | 0.11 | 2 | -1.74 | 0.07 | 2 | 19 | 2 | 2 | 16.5 | |
| | 333 | ICK10 | Dorsal vertebra | Posterior | Dorsal | 17.6 | 0.2 | 5 | | | | | | | | | | | | | 19.0 | |
| | 338 | ICK11 | Dorsal vertebra | Transition cervicale-dorsale | Dorsal | 17.5 | 0.1 | 5 | 25.1 | 0.1 | 3 | -5.65 | 0.1 | 3 | -5.66 | 0.09 | 3 | 7 | 1 | 3 | 26.2 | |
| | 339 | ICK12 | Dorsal vertebra | Transition cervicale-dorsale | Dorsal | 17.5 | 0.2 | 4 | | | | | | | | | | | | | 21.5 | |
| | | ICK16 | Rib | | Dorsal | 17.6 | 0.2 | 5 | | | | | | | | | | | | | 14.9 | |
| | | ICK17 | Rib | | Dorsal | 19.3 | 0.3 | 3 | 27 | 0.22 | 3 | -3.83 | 0.22 | 3 | -8.56 | 0.01 | 3 | 5 | 0 | 3 | 28.8 | |
| | | ICK18 | Rib | | Dorsal | 17.5 | 0.2 | 3 | | | | | | | | | | | | | 23.9 | |
| | | ICK21 | Left coracoid | | Dorsal | 17.9 | 0.1 | 5 | 27.2 | 0.23 | 3 | -3.62 | 0.23 | 3 | -8.41 | 0.02 | 3 | 4 | 0 | 3 | 19.6 | |
| | | | | | *Min.* | 16.1 | | | 24.10 | | | -6.58 | | | -8.56 | | | 4 | | | 14.9 | |
| | | | | | *Max.* | 20.2 | | | 27.20 | | | -3.62 | | | -1.56 | | | 19 | | | 28.8 | |
| | | | | | *Mean / SEM* | 18.4 | 1.2 | | 25.78 | 0.94 | | -4.99 | 0.89 | | -4.57 | 2.76 | | 11 | 5 | | 21.4 | 4.4 |
| | | ICK19 | Right nasal | Proximal part | Skull | 17.7 | 0.1 | 5 | 27.4 | 0.4 | 3 | -3.43 | 0.4 | 3 | -8.39 | 0.14 | 3 | 2 | 0 | 3 | 32.8 | |
| | | ICK20 | Right dentary | Proximal part | Skull | 17.6 | 0.1 | 5 | | | | | | | | | | | | | 29.5 | |
| | | ICK34 | Left angular | Proximal part | Skull | 17.7 | 0.2 | 5 | 26.7 | 0.18 | 3 | -4.08 | 0.18 | 3 | -4.16 | 0.28 | 3 | 8 | 1 | 3 | 26.4 | |
| | | ICK35 | Left dentary | Distal part | Skull | 18.2 | 0.2 | 5 | 27.5 | 0.2 | 3 | -3.33 | 0.2 | 3 | -8.46 | 0.01 | 3 | 4 | 0 | 3 | 34.6 | |
| | | ICK38 | Left premaxilla | | Skull | 18.2 | 0.2 | 4 | 28.2 | 0.31 | 3 | -2.61 | 0.31 | 3 | -8.19 | 0.15 | 3 | 2 | 0 | 3 | 33.4 | |
| | | | | | *Min.* | 17.6 | | | 26.70 | | | -4.08 | | | -8.46 | | | 2 | | | 26.4 | |
| | | | | | *Max.* | 18.2 | | | 28.20 | | | -2.61 | | | -4.16 | | | 8 | | | 34.6 | |
| | | | | | *Mean / SEM* | 17.9 | 0.3 | | 27.45 | 0.53 | | -3.36 | 0.52 | | -7.30 | 1.82 | | 4 | 2 | | 31.3 | 3.0 |
| | | ICK36 | Tooth | On the premaxilla | Tooth | 18.3 | 0.2 | 4 | 28.3 | 0.23 | 3 | -2.56 | 0.23 | 3 | -8.18 | 0.26 | 3 | 2 | 0 | 3 | 15.9 | |
| | | | | | **Global Min.** | 16.1 | | | 24.10 | | | -6.58 | | | -8.56 | | | 2 | | | 14.8 | |
| | | | | | **Global Max.** | 20.2 | | | 28.30 | | | -2.56 | | | -1.56 | | | 24 | | | 34.6 | |
| | | | | | **Global Mean / SEM** | 18.2 | 0.8 | | 26.34 | 1.17 | | -4.45 | 1.13 | | -5.79 | 2.68 | | 9 | 6 | | 23.7 | 5.3 |
| | | | | | **Global Δ** | 4.1 | | | 4.20 | | | 4.02 | | | 7.00 | | | 22 | | | 19.8 | |
| | | | | | **Global Mid-range** | 18.2 | | | 26.20 | | | -4.57 | | | -5.06 | | | 13 | | | 24.7 | |

underlined: Data removed because mineralogy alt underlined: Data removed because mineralogy altered

**in bold: Data removed because %wt CO3 > 13.4% in bold: Data removed because %wt CO3 > 13.4%**

| Taxa | Coll. N° | #Sample | Skeletal element | Additionnal information | Skeletal region | $\delta^{18}O_p$ (‰, V-SMOW) | | | $\delta^{18}O_c$ (‰, V-SMOW) | | | $\delta^{18}O_c$ (‰, V-PDB) | | | $\delta^{13}C_c$ (‰, V-PDB) | | | wt% $CO_3^{2-}$ | | | wt% $P_2O_5$ | |
|---|---|---|---|---|---|---|---|---|---|---|---|---|---|---|---|---|---|---|---|---|---|---|
| | | | | | | Mean | SD | N | Mean | SD | N | Mean | SD | N | Mean | SD | N | Mean | SD | N | | |
| *Palvennia hoybergeti* | PMO 222.66 | PH5 | Left humerus | | Appendicular | 15.4 | 0.2 | 5 | | | | | | | | | | | | | 16.9 | |
| | PMO 222.66 | PH6 | Preaccessory element | | Appendicular | 14.2 | 0.2 | 3 | | | | | | | | | | | | | 9.5 | |
| | PMO 222.66 | PH7 | Ulna | | Appendicular | | | | | | | | | | | | | | | | 4.6 | |
| | PMO 222.66 | PH8 | Limb bone | | Appendicular | 15.3 | 0.1 | 2 | | | | | | | | | | | | | 4.3 | |
| | PMO 222.66 | PH17 | Right humerus | | Appendicular | 15.3 | 0 | 5 | | | | | | | | | | | | | 21.5 | |
| | PMO 222.66 | PH18 | Right radius | | Appendicular | 15.3 | 0.2 | 5 | | | | | | | | | | | | | 16.6 | |
| | PMO 222.66 | PH19 | Phalange ? | | Appendicular | | | | | | | | | | | | | | | | 2.0 | |
| | PMO 222.66 | PH20 | Pisiform | | Appendicular | 15 | 0.2 | 5 | | | | | | | | | | | | | 25.4 | |
| | PMO 222.66 | PH21 | Limb bone | | Appendicular | 14.2 | 0.1 | 5 | 18.50 | 0.17 | 3 | -12.03 | 0.17 | 3 | -9.31 | 0.08 | 3 | 3 | 0 | 3 | 19.3 | |
| | PMO 222.66 | PH22 | Limb bone | | Appendicular | 14.5 | 0.1 | 5 | | | | | | | | | | | | | 18.1 | |
| | PMO 222.66 | PH23 | Carpal distal IV | | Appendicular | 14.9 | 0.2 | 5 | | | | | | | | | | | | | 17.8 | |
| | PMO 222.66 | PH24 | Ulnare | | Appendicular | 15 | 0.1 | 5 | | | | | | | | | | | | | 26.0 | |
| | | | | | *Min.* | 14.2 | | | | | | | | | | | | | | | 2.0 | |
| | | | | | *Max.* | 15.4 | | | | | | | | | | | | | | | 26.0 | |
| | | | | | *Mean / SEM* | 14.9 | 0.4 | | | | | | | | | | | | | | 15.2 | 7.8 |
| | PMO 222.66 | PH9 | Rib fragment | | Dorsal | 15.8 | 0.2 | 5 | | | | | | | | | | | | | 15.7 | |
| | PMO 222.66 | PH10 | Rib fragment | | Dorsal | 15.5 | 0.1 | 5 | | | | | | | | | | | | | 26.9 | |
| | PMO 222.66 | PH11 | Vertebra | | Dorsal | 15.8 | 0.1 | 5 | | | | | | | | | | | | | 16.4 | |
| | PMO 222.66 | PH12 | Vertebra | | Dorsal | 15.3 | 0.2 | 5 | 19.00 | 0.16 | 3 | -11.56 | 0.16 | 3 | -10.60 | 0.1 | 3 | 2 | 0 | 3 | 12.2 | |
| | PMO 222.66 | PH13 | Rib fragment | | Dorsal | 15.8 | 0.2 | 5 | | | | | | | | | | | | | 24.1 | |
| | PMO 222.66 | PH16 | Atlas-axis | | Dorsal | 14.8 | 0 | 2 | | | | | | | | | | | | | 5.1 | |
| | PMO 222.66 | PH25 | Coracoid | | Dorsal | 14.6 | 0.1 | 5 | 19.10 | 0.15 | 3 | -11.43 | 0.15 | 3 | -8.70 | 0.07 | 3 | 4 | 0 | 3 | 25.5 | |

| | | | | | | | | | | | | | | | | | | | | | | |
|---|---|---|---|---|---|---|---|---|---|---|---|---|---|---|---|---|---|---|---|---|---|---|
| | PMO 222.66 | PH26 | Rib | | Dorsal | 14.8 | 0.2 | 5 | 18.20 | 0.33 | 3 | -12.37 | 0.33 | 3 | -7.88 | 0.66 | 3 | 5 | 0 | 3 | 28.0 | |
| | PMO 222.66 | PH27 | Rib | | Dorsal | 15.2 | 0.2 | 5 | | | | | | | | | | | | | 25.8 | |
| | PMO 222.66 | PH28 | Rib | | Dorsal | 14.4 | 0.2 | 5 | | | | | | | | | | | | | 27.3 | |
| | PMO 222.66 | PH29 | Rib | | Dorsal | 15.5 | 0.1 | 5 | | | | | | | | | | | | | 20.8 | |
| | | | | | *Min.* | 14.4 | | | 18.20 | | | -12.37 | | | -10.60 | | | 2 | | | 5.1 | |
| | | | | | *Max.* | 15.8 | | | 19.10 | | | -11.43 | | | -7.88 | | | 5 | | | 28.0 | |
| | | | | | *Mean / SEM* | 15.2 | 0.5 | | 18.77 | 0.40 | | -11.79 | 0.42 | | -9.06 | 1.14 | | 4 | 1 | | 20.7 | 7.1 |
| | PMO 222.66 | PH14 | Left dentary | | Skull | 15.2 | 0.1 | 5 | 18.30 | 0.30 | 3 | -12.27 | 0.3 | 3 | -9.89 | 0.12 | 3 | 5 | 1 | 3 | 32.9 | |
| | PMO 222.66 | PH15 | Left premaxilla | | Skull | 14.1 | 0.2 | 4 | | | | | | | | | | | | | 15.0 | |
| | | | | | *Min.* | 14.1 | | | | | | | | | | | | | | | 15.0 | |
| | | | | | *Max.* | 15.2 | | | | | | | | | | | | | | | 32.9 | |
| | | | | | *Mean / SEM* | 14.7 | 0.6 | | | | | | | | | | | | | | 23.9 | 8.9 |
| | - | PH1 | Tooth | | Teeth | 15.9 | 0.1 | 5 | | | | | | | | | | | | | 23.9 | |
| | - | PH2 | Tooth | | Teeth | 15.7 | 0.2 | 5 | | | | | | | | | | | | | 20.9 | |
| | - | PH3 | Tooth | | Teeth | 15.5 | 0.2 | 5 | | | | | | | | | | | | | 17.7 | |
| | - | PH4 | Tooth | | Teeth | 15.7 | 0.2 | 5 | | | | | | | | | | | | | 24.6 | |
| | | | | | *Min.* | 15.5 | | | | | | | | | | | | | | | 17.7 | |
| | | | | | *Max.* | 15.9 | | | | | | | | | | | | | | | 24.6 | |
| | | | | | *Mean / SEM* | 15.7 | 0.1 | | | | | | | | | | | | | | 21.8 | 2.7 |
| | | | | | **Global Min.** | **14.1** | | | **18.20** | | | **-12.37** | | | **-10.60** | | | **2** | | | **2.0** | |
| | | | | | **Global Max.** | **15.9** | | | **19.10** | | | **-11.43** | | | **-7.88** | | | **5** | | | **32.9** | |
| | | | | | **Global Mean / SEM** | **15.1** | **0.5** | | **18.62** | **0.41** | | **-11.93** | **0.42** | | **-9.28** | **1.05** | | **4** | **1** | | **18.8** | **7.9** |
| | | | | | **Global Δ** | **1.8** | | | **0.90** | | | **0.94** | | | **2.72** | | | **3** | | | **30.8** | |
| | | | | | **Global Mid-range** | **15** | | | **18.65** | | | **-11.9** | | | **-9.24** | | | **3.5** | | | **17.5** | |

underlined: Data removed because mineralogy altered

**in bold: Data removed because %wt CO3 > 13.4%**

| Taxa | Coll. N° | #Sample | Skeletal element | Additionnal information | Skeletal region | $\delta^{18}O_p$ (‰, V-SMOW) | | | $\delta^{18}O_c$ (‰, V-SMOW) | | | $\delta^{18}O_c$ (‰, V-PDB) | | | $\delta^{13}C_c$ (‰, V-PDB) | | | wt% $CO_3^{2-}$ | | | wt% $P_2O_5$ | |
|---|---|---|---|---|---|---|---|---|---|---|---|---|---|---|---|---|---|---|---|---|---|---|
| | | | | | | Mean | SD | N | Mean | SD | N | Mean | SD | N | Mean | SD | N | Mean | SD | N | | |
| *Keilhauia* sp. | PMO 222.66 | K8 | Right humerus | | Appendicular | 15.1 | 0.1 | 5 | 18.37 | 0.2 | 3 | -12.17 | 0.2 | 3 | -6.77 | 0.05 | 3 | 5 | 0 | 3 | 30.0 | |
| | PMO 222.66 | K9 | Limb bone | | Appendicular | 13.4 | 0.2 | 5 | | | | | | | | | | | | | 13.9 | |
| | PMO 222.66 | K10 | Limb bone | | Appendicular | 13.7 | 0.1 | 5 | | | | | | | | | | | | | 18.9 | |
| | PMO 222.66 | K11 | Limb bone | | Appendicular | 14.3 | 0 | 1 | | | | | | | | | | | | | 40.5 | |
| | PMO 222.66 | K12 | Phalange ? | | Appendicular | 13.8 | 0.2 | 5 | | | | | | | | | | | | | 9.7 | |
| | PMO 222.66 | K13 | Phalange ? | | Appendicular | 14.4 | 0.1 | 5 | | | | | | | | | | | | | 11.3 | |
| | PMO 222.66 | K14 | Phalange ? | | Appendicular | 13.4 | 0.1 | 5 | | | | | | | | | | | | | 13.2 | |
| | PMO 222.66 | K15 | Distal phalange | | Appendicular | 13.7 | 0.1 | 5 | | | | | | | | | | | | | 10.6 | |
| | PMO 222.66 | K16 | Proximal phalange | | Appendicular | 14 | 0.1 | 5 | | | | | | | | | | | | | 10.6 | |
| | PMO 222.66 | K17 | Right radius? | | Appendicular | 14.1 | 0.1 | 5 | | | | | | | | | | | | | 20.0 | |
| | | | | | *Min.* | 13.4 | | | | | | | | | | | | | | | 9.7 | |
| | | | | | *Max.* | 15.1 | | | | | | | | | | | | | | | 40.5 | |
| | | | | | *Mean / SEM* | 14.0 | 0.5 | | | | | | | | | | | | | | 17.9 | 9.6 |
| | PMO 222.66 | K6 | Atlas-axis | | Dorsal | 12.8 | 0.1 | 5 | | | | | | | | | | | | | 22.3 | |
| | PMO 222.66 | K7 | Right scapula | | Dorsal | 13.8 | 0.1 | 5 | | | | | | | | | | | | | 20.3 | |
| | PMO 222.66 | K18 | Preflexural vertebra | | Dorsal | 13.2 | 0.1 | 5 | | | | | | | | | | | | | 10.3 | |
| | PMO 222.66 | K19 | Preflexural vertebra | | Dorsal | 11.8 | 0.1 | 5 | | | | | | | | | | | | | 15.5 | |
| | - | K20 | Rib fragment | | Dorsal | 13.9 | 0.1 | 5 | | | | | | | | | | | | | 25.5 | |
| | - | K21 | Rib fragment | | Dorsal | 15 | 0.1 | 5 | | | | | | | | | | | | | 33.1 | |
| | - | K22 | Rib fragment | | Dorsal | 13.7 | 0.1 | 5 | 17.51 | 0.01 | 2 | -13.01 | 0.01 | 2 | -8.4 | 0.40 | 2 | 2 | 0 | 3 | 23.7 | |
| | - | K23 | Rib fragment | | Dorsal | 14.4 | 0.1 | 5 | 17.87 | 0.1 | 3 | -12.66 | 0.10 | 3 | -8.05 | 0.29 | 3 | 2 | 0 | 3 | 21.8 | |
| | - | K24 | Rib fragment | | Dorsal | 13.9 | 0.2 | 5 | | | | | | | | | | | | | 21.2 | |
| | PMO 222.66 | K25 | Right coracoid | | Dorsal | 15.1 | 0.1 | 5 | | | | | | | | | | | | | 36.8 | |
| | | | | | *Min.* | 11.8 | | | 17.51 | | | -13.01 | | | -8.40 | | | 2 | | | 10.3 | |
| | | | | | *Max.* | 15.1 | | | 17.87 | | | -12.66 | | | -8.05 | | | 2 | | | 36.8 | |
| | | | | | *Mean / SEM* | 13.8 | 0.9 | | 17.69 | 0.18 | | -12.84 | 0.18 | | -8.23 | 0.18 | | 2 | 0 | | 23.0 | 7.3 |
| | PMO 222.66 | K4 | Dentary | | Skull | 14.2 | 0.1 | 5 | | | | | | | | | | | | | 30.5 | |
| | PMO 222.66 | K5 | Left quadrate | | Skull | 13.7 | 0.2 | 3 | | | | | | | | | | | | | 25.7 | |
| | | | | | *Min.* | 13.7 | | | | | | | | | | | | | | | 25.7 | |
| | | | | | *Max.* | 14.2 | | | | | | | | | | | | | | | 30.5 | |
| | | | | | *Mean / SEM* | 14.0 | 0.3 | | | | | | | | | | | | | | 28.1 | 2.4 |
| | PMO 222.66 | K1 | Tooth | | Teeth | 14.8 | 0.2 | 5 | | | | | | | | | | | | | 26.6 | |
| | PMO 222.66 | K2 | Tooth | | Teeth | 15.1 | 0.3 | 4 | | | | | | | | | | | | | 30.2 | |
| | PMO 222.66 | K3 | Tooth | | Teeth | 14.7 | 0.2 | 2 | | | | | | | | | | | | | 25.3 | |
| | | | | | *Min.* | 14.7 | | | | | | | | | | | | | | | 25.3 | |
| | | | | | *Max.* | 15.1 | | | | | | | | | | | | | | | 30.2 | |
| | | | | | *Mean / SEM* | 14.9 | 0.2 | | | | | | | | | | | | | | 27.4 | 2.0 |
| | | | | | **Global Min.** | **11.8** | | | **17.51** | | | **-13.01** | | | **-8.4** | | | **2** | | | **9.7** | |
| | | | | | **Global Max.** | **15.1** | | | **18.37** | | | **-12.17** | | | **-6.77** | | | **5** | | | **40.5** | |
| | | | | | **Global Mean / SEM** | **14.0** | **0.8** | | **17.92** | **0.35** | | **-12.61** | **0.34** | | **-7.74** | **0.70** | | **3** | **1** | | **21.9** | **8.5** |
| | | | | | **Global Δ** | **3.3** | | | **0.86** | | | **0.84** | | | **1.63** | | | **3** | | | **30.8** | |
| | | | | | **Global Mid-range** | **13.5** | | | **17.9** | | | **-12.6** | | | **-7.6** | | | **4** | | | **25.1** | |

underlined: Data removed because mineralogy altered

**in bold: Data removed because %wt CO3 > 13.4%**

| Taxa | Coll. N° | #Sample | Skeletal element | Additional information | Skeletal region | $\delta^{18}O_p$ (‰, V-SMOW) | | | $\delta^{18}O_c$ (‰, V-SMOW) | | | $\delta^{18}O_c$ (‰, V-PDB) | | | $\delta^{13}C_c$ (‰, V-PDB) | | | wt% $CO_3^{2-}$ | | | wt% $P_2O_5$ |
|---|---|---|---|---|---|---|---|---|---|---|---|---|---|---|---|---|---|---|---|---|---|
| | | | | | | Mean | SD | N | Mean | SD | N | Mean | SD | N | Mean | SD | N | Mean | SD | N | |
| *Keilhauia nui* | | Mi1 | Presacral vertebra | Approximately the 5th | Dorsal | 13.1 | 0,3 | 5 | | | | | | | | | | | | | 7.8 |
| | | Mi2 | Presacral vertebra | Approximately the 6th | Dorsal | | | | | | | | | | | | | | | | 0.1 |
| | | Mi3 | Presacral vertebra | Approximately the 7th | Dorsal | 15.6 | 0,1 | 4 | | | | | | | | | | | | | 10.1 |
| | | Mi4 | Presacral vertebra | Approximately the 8th | Dorsal | 15.7 | 0,2 | 5 | | | | | | | | | | | | | 15.0 |
| | | Mi5 | Presacral vertebra | Approximately the 16th | Dorsal | 15.3 | 0,1 | 2 | | | | | | | | | | | | | 3.3 |
| | | Mi6 | Presacral vertebra | Approximately the 17th | Dorsal | | | | | | | | | | | | | | | | 0.3 |
| | | Mi7 | Presacral vertebra | Approximately the 18th | Dorsal | | | | | | | | | | | | | | | | 0.6 |
| | | Mi8 | Presacral vertebra | Approximately the 29th | Dorsal | 14.4 | 0,0 | 2 | | | | | | | | | | | | | 5.6 |

| | | | | | | | | | | | | | | | | | | | | | | |
|---|---|---|---|---|---|---|---|---|---|---|---|---|---|---|---|---|---|---|---|---|---|---|
| | | Mi9 | Presacral vertebra | Approximately the 39th | Dorsal | | | | | | | | | | | | | | | | 1.2 | |
| | | Mi23 | Rib fragment | | Dorsal | | | | | | | | | | | | | | | | | |
| | | Mi24 | Rib fragment | | Dorsal | | | | | | | | | | | | | | | | | |
| | | | | | *Min.* | 13.1 | | | | | | | | | | | | | | | 0.1 | |
| | | | | | *Max.* | 15.7 | | | | | | | | | | | | | | | 15.0 | |
| | | | | | *Mean / SEM* | 14.8 | 1.0 | | | | | | | | | | | | | | 4.9 | 5.0 |
| | | Mi10 | Right humerus | | Appendicular | | | | | | | | | | | | | | | | | |
| | | Mi11 | Radius | | Appendicular | 14.9 | 0,1 | 5 | | | | | | | | | | | | | 6.1 | |
| | | Mi12 | Limb bone | | Appendicular | | | | | | | | | | | | | | | | 0.5 | |
| | | Mi13 | Limb bone | | Appendicular | | | | | | | | | | | | | | | | 0.6 | |
| | | Mi14 | Caudal vertebra | Near flexural region | Caudal | | | | | | | | | | | | | | | | 0.4 | |
| | | Mi15 | Caudal vertebra | Approximately the 54th | Caudal | 14.8 | 0,1 | 4 | | | | | | | | | | | | | 8.5 | |
| | | Mi16 | Caudal vertebra | Approximately the 55th | Caudal | | | | | | | | | | | | | | | | | |
| | | Mi17 | Caudal vertebra | Approximately the 56th | Caudal | | | | | | | | | | | | | | | | | |
| | | Mi18 | Caudal vertebra | ›ximately the 71th, probably post-fle | Caudal | | | | | | | | | | | | | | | | | |
| | | Mi19 | Caudal vertebra | ›ximately the 70th, probably post-fle | Caudal | | | | | | | | | | | | | | | | | |
| | | Mi20 | Caudal vertebra | ›ximately the 68th, probably post-fle | Caudal | | | | | | | | | | | | | | | | | |
| | | Mi21 | Caudal vertebra | ›ximately the 64th, probably post-fle | Caudal | | | | | | | | | | | | | | | | | |
| | | Mi22 | Dentary | | Skull | | | | | | | | | | | | | | | | | |
| | | | | | **Global Min.** | **13.1** | | | | | | | | | | | | | | | **0.1** | |
| | | | | | **Global Max.** | **15.7** | | | | | | | | | | | | | | | **15.0** | |
| | | | | | **Global Mean / SEM** | **14.8** | **0.9** | | | | | | | | | | | | | | **5.7** | **4.8** |
| | | | | | **Global Δ** | **2.6** | | | | | | | | | | | | | | | **14.8** | |
| | | | | | **Global Mid-range** | **14.4** | | | | | | | | | | | | | | | **7.5** | |

underlined: Data removed because mineralogy altered

**in bold: Data removed because %wt CO3 > 13.4%**

| Taxa | Coll. N° | #Sample | Skeletal element | Additionnal information | Skeletal region | $\delta^{18}O_p$ (‰, V-SMOW) | | | $\delta^{18}O_c$ (‰, V-SMOW) | | | $\delta^{18}O_c$ (‰, V-PDB) | | | $\delta^{13}C_c$ (‰, V-PDB) | | | wt% $CO_3^{2-}$ | | | wt% $P_2O_5$ | |
|---|---|---|---|---|---|---|---|---|---|---|---|---|---|---|---|---|---|---|---|---|---|---|
| | | | | | | Mean | SD | N | Mean | SD | N | Mean | SD | N | Mean | SD | N | Mean | SD | N | | |
| *Colymbosaurus svalbardensis* | - | G22 | Left humerus | | ALL | 14.9 | 0.1 | 5 | 18.44 | 0.1 | 3 | -12.05 | 0.1 | 3 | -7.71 | 0.04 | 3 | 4 | 0 | | 23.66 | |
| | PMO 222.663/1 | G23 | Right humerus | | ARL | 15 | 0.1 | 5 | 19.36 | 0.33 | 3 | -11.16 | 0.33 | 3 | -8.22 | 0.04 | 3 | 2 | 0 | 3 | 14.9 | |
| | PMO 222.663/1 | G26 | Right radius | | ARL | 14.2 | 0.1 | 5 | | | | | | | | | | | | | 16.5 | |
| | PMO 222.663/1 | G27 | Post-axial accessory bone | | ARL | 15.1 | 0.1 | 3 | | | | | | | | | | | | | 23.4 | |
| | PMO 222.663/1 | G28 | Intermedium | | ARL | 14.5 | 0.1 | 5 | | | | | | | | | | | | | 23.1 | |
| | PMO 222.663/1 | G29 | Distal carpal finger II | | ARL | 15.7 | 0.1 | 5 | | | | | | | | | | | | | 21.4 | |
| | | | | | *Min.* | 14.2 | | | | | | | | | | | | | | | 14.9 | |
| | | | | | *Max.* | 15.7 | | | | | | | | | | | | | | | 23.4 | |
| | | | | | *Mean / SEM* | 14.9 | 0.6 | | | | | | | | | | | | | | 19.9 | 3.9 |
| | PMO 222.663/2 | G17 | Caudal vertebra | | Caudal | 14.4 | 0.3 | 5 | | | | | | | | | | | | | 15.5 | |
| | - | G18 | Caudal vertebra | Last one | Caudal | 13.7 | 0.1 | 5 | | | | | | | | | | | | | 17.4 | |
| | - | G38 | Caudal vertebra | Anterior to G39 | Caudal | 13.6 | 0.2 | 5 | 19.32 | 0.01 | 2 | -11.25 | 0.01 | 2 | -9.27 | 0.03 | 2 | 2 | 0 | 2 | 18.4 | |
| | - | G39 | Caudal vertebra | Anterior to G40 | Caudal | 13.5 | 0.1 | 5 | | | | | | | | | | | | | 17.6 | |
| | - | G40 | Caudal vertebra | | Caudal | 13.8 | 0.2 | 4 | | | | | | | | | | | | | 19.7 | |
| | | | | | *Min.* | 13.5 | | | | | | | | | | | | | | | 15.5 | |
| | | | | | *Max.* | 14.4 | | | | | | | | | | | | | | | 19.7 | |
| | | | | | *Mean / SEM* | 13.8 | 0.4 | | | | | | | | | | | | | | 17.7 | 1.5 |
| | PMO 222.663/1 | G19 | Sacral vertebra | | Dorsal | 13.2 | 0.2 | 5 | | | | | | | | | | | | | 21.9 | |
| | - | G20 | Dorsal vertebra | | Dorsal | 14.4 | 0.1 | 4 | 18.1 | 0.23 | 3 | -12.44 | 0.23 | 3 | -9.76 | 0.08 | 3 | 3 | 0 | 3 | 27.2 | |
| | PMO 222.663/1 | G21 | Sacral vertebra | | Dorsal | 13.7 | 0.2 | 5 | | | | | | | | | | | | | 21.1 | |
| | PMO 222.663/1 | G30 | Dorsal vertebra | | Dorsal | 15.7 | 0.1 | 5 | 18.76 | 0.08 | 3 | -11.74 | 0.08 | 3 | -8.85 | 0.06 | 3 | 5 | 0 | 3 | 28.6 | |
| | PMO 222.663/1 | G31 | Dorsal vertebra | | Dorsal | 15.9 | 0.1 | 5 | | | | | | | | | | | | | 28.4 | |
| | PMO 222.663/1 | G32 | Dorsal vertebra | | Dorsal | 14.5 | 0.1 | 4 | 18.65 | 0.01 | 3 | -11.9 | 0.01 | 3 | -9.27 | 0.05 | 3 | 4 | 0 | 3 | 30.9 | |
| | - | G33 | Rib fragment | | Dorsal | 15.6 | 0.1 | 5 | | | | | | | | | | | | | 25.9 | |
| | - | G34 | Rib fragment | | Dorsal | 15.7 | 0.1 | 5 | | | | | | | | | | | | | 25.3 | |
| | PMO 222.663/1 | G35 | Rib | | Dorsal | 13.7 | 0.1 | 4 | 17.84 | 0.08 | 3 | -12.69 | 0.08 | 3 | -10.01 | 0.07 | 3 | 2 | 0 | 3 | 29.9 | |
| | PMO 222.663/1 | G36 | Rib | | Dorsal | 14.6 | 0.2 | 4 | | | | | | | | | | | | | 32.4 | |
| | - | G37 | Clavicle-interclavicle | | Dorsal | 15.2 | 0.2 | 3 | | | | | | | | | | | | | 26.6 | |
| | - | G41 | Rib | | Dorsal | 14.4 | 0.1 | 5 | 20.06 | 0.08 | 2 | -10.48 | 0.06 | 2 | -8.8 | 0.01 | 2 | 2 | 0 | 2 | 25.5 | |
| | | | | | *Min.* | 13.2 | | | 17.84 | | | -12.69 | | | -10.01 | | | 2 | | | 21.1 | |
| | | | | | *Max.* | 15.9 | | | 20.06 | | | -10.48 | | | -8.8 | | | 5 | | | 32.4 | |
| | | | | | *Mean / SEM* | 14.7 | 0.9 | | 18.7 | 0.9 | | -11.9 | 0.9 | | -9.3 | 0.5 | | 3 | 1 | | 27.0 | 3.4 |
| | PMO 222.663/1 | G9 | Left fibula | | PLL | 13.7 | 0.2 | 5 | | | | | | | | | | | | | 19.6 | |
| | PMO 222.663/ | G10 | Metatarsal of the finger I | | PLL | 13.8 | 0.2 | 4 | | | | | | | | | | | | | 9.9 | |
| | PMO 222.663/1 | G11 | V,1 | | PLL | 13.1 | 0.1 | 5 | | | | | | | | | | | | | 20.7 | |
| | PMO 222.663/ | G12 | III,2 | | PLL | 14.2 | 0.1 | 5 | | | | | | | | | | | | | 17.2 | |
| | PMO 222.663/ | G13 | II, 5 | | PLL | 13.3 | 0.1 | 5 | | | | | | | | | | | | | 22.9 | |
| | PMO 222.663/ | G14 | V, 5 | | PLL | 13.3 | 0.1 | 5 | | | | | | | | | | | | | 14.7 | |
| | PMO 222.663/ | G15 | IV, 6 | | PLL | 14 | 0.1 | 5 | | | | | | | | | | | | | 21.6 | |
| | PMO 222.663/ | G16 | III, 13 | | PLL | 13.3 | 0.2 | 5 | | | | | | | | | | | | | 8.4 | |
| | PMO 222.663/1 | G24 | Left femur | | PLL | 13.2 | 0.1 | 5 | | | | | | | | | | | | | 28.8 | |
| | | | | | *Min.* | 13.1 | | | | | | | | | | | | | | | 8.4 | |
| | | | | | *Max.* | 14.2 | | | | | | | | | | | | | | | 28.8 | |
| | | | | | *Mean / SEM* | 13.5 | 0.4 | | | | | | | | | | | | | | 18.2 | 6.4 |
| | PMO 222.663/ | G1 | Right fibula | | PRL | 14.2 | 0.2 | 5 | 18.38 | 0.16 | 3 | -12.16 | 0.16 | 3 | -8.04 | 0.11 | 3 | 2 | 0 | 3 | 15.3 | |
| | PMO 222.663/ | G2 | Metatarsal of the finger I | | PRL | | | | | | | | | | | | | | | | | |
| | PMO 222.663/1 | G3 | V,1 | | PRL | 14.4 | 0.1 | 4 | | | | | | | | | | | | | 17.3 | |
| | PMO 222.663/ | G4 | III,2 | | PRL | 14.6 | 0.1 | 4 | 17.94 | 0.08 | 3 | -12.59 | 0.08 | 3 | -8.18 | 0.01 | 3 | 3 | 0 | 3 | 20.1 | |
| | PMO 222.663/ | G5 | II, 5 | | PRL | 14.3 | 0.1 | 5 | | | | | | | | | | | | | 24.5 | |
| | PMO 222.663/ | G6 | V, 5 | | PRL | 14 | 0.3 | 5 | | | | | | | | | | | | | 20.3 | |
| | PMO 222.663/ | G7 | IV, 6 | | PRL | 14.1 | 0.1 | 5 | 19.51 | 0.01 | 3 | -11.07 | 0.01 | 3 | -8.37 | 0.03 | 3 | 4 | 0 | 3 | 19.9 | |
| | PMO 222.663/ | G8 | III, 11 | | PRL | 14 | 0.2 | 5 | 17.84 | 0.25 | 2 | -12.69 | 0.25 | 2 | -7.92 | 0.02 | 2 | 2 | 1 | 2 | 18.1 | |
| | PMO 222.663/ | G25 | Right femur | | PRL | 13.7 | 0.1 | 5 | 17.83 | 0.07 | 3 | -12.7 | 0.07 | 3 | -9.49 | 0.51 | 3 | 2 | 0 | 3 | 27.4 | |
| | | | | | *Min.* | 13.7 | | | 17.83 | | | -12.7 | | | -9.49 | | | 2 | | | 15.3 | |
| | | | | | *Max.* | 14.6 | | | 19.51 | | | -11.07 | | | -7.92 | | | 4 | | | 27.4 | |

| | | | | | | | | | | | | | | | | | | | | | | |
|---|---|---|---|---|---|---|---|---|---|---|---|---|---|---|---|---|---|---|---|---|---|---|
| | | | | | *Mean / SEM* | 14.2 | 0.3 | | 18.3 | 0.7 | | -12.2 | 0.7 | | -8.4 | 0.6 | | 3 | 1 | | 20.4 | 3.9 |
| | | | | | **Global Min.** | **13.1** | | | **17.83** | | | **-12.70** | | | **-10.01** | | | **2** | | | **8.4** | |
| | | | | | **Global Max.** | **15.9** | | | **20.06** | | | **-10.48** | | | **-7.71** | | | **5** | | | **32.4** | |
| | | | | | **Global Mean / SEM** | **14.3** | **0.8** | | **18.62** | **0.74** | | **-11.92** | **0.73** | | **-8.76** | **0.75** | | **3** | **1** | | **21.5** | **5.5** |
| | | | | | **Global Δ** | **2.8** | | | **2.23** | | | **2.22** | | | **2.30** | | | **3** | | | **24.0** | |
| | | | | | **Global Mid-range** | **14.5** | | | **18.945** | | | **-11.59** | | | **-8.86** | | | **4** | | | **20.4** | |

| Taxa | Coll. N° | #Sample | Skeletal element | Additional information | Skeletal region | $\delta^{18}O_p$ (‰, V-SMOW) | | | $\delta^{18}O_c$ (‰, V-SMOW) | | | $\delta^{18}O_c$ (‰, V-PDB) | | | $\delta^{13}C_c$ (‰, V-PDB) | | | wt% $CO_3^{2-}$ | | | wt% $P_2O_5$ | |
|---|---|---|---|---|---|---|---|---|---|---|---|---|---|---|---|---|---|---|---|---|---|---|
| | | | | | | Mean | SD | N | Mean | SD | N | Mean | SD | N | Mean | SD | N | Mean | SD | N | | |
| Cryptoclididae indet. | | SS9 | Propodial fragment | | Appendicular | 16.6 | 0.1 | 5 | 18.37 | 0.29 | 3 | -12.11 | 0.29 | 3 | -10.73 | 0.13 | 3 | 5 | 1 | 3 | 26.0 | |
| | | SS30 | Left humerus | | Appendicular | 16.5 | 0.2 | 5 | | | | | | | | | | | | | 21.5 | |
| | PMO 212.662/0 | SS31 | Left radius | | Appendicular | 16.8 | 0.1 | 5 | | | | | | | | | | | | | 20.3 | |
| | PMO 212.662/0 | SS32 | Left ulna | | Appendicular | 16 | 0.1 | 5 | | | | | | | | | | | | | 19.2 | |
| | PMO 212.662/0 | SS33 | Left intermedium | | Appendicular | 16.9 | 0.1 | 4 | | | | | | | | | | | | | 21.9 | |
| | PMO 212.662/0 | SS34 | Metacarpal #4 | | Appendicular | 16 | 0.2 | 4 | 20.63 | 0.41 | 2 | -9.92 | 0.41 | 3 | -10.10 | 0.04 | 3 | 2 | 0 | 2 | 17.5 | |
| | PMO 212.662/0 | SS35 | I,1 | | Appendicular | 16.5 | 0.2 | 5 | | | | | | | | | | | | | 20.4 | |
| | PMO 212.662/0 | SS36 | III, 2 | | Appendicular | 17.1 | 0.1 | 5 | | | | | | | | | | | | | 25.4 | |
| | PMO 212.662/0 | SS37 | II,4 | | Appendicular | 16.7 | 0.2 | 5 | 18.00 | 0.04 | 3 | -12.57 | 0.04 | 3 | -9.56 | 0.13 | 3 | 5 | 0 | 3 | 22.5 | |
| | PMO 212.662/0 | SS38 | Distal phalange | | Appendicular | 16.5 | 0.2 | 5 | 20.00 | 0.01 | 2 | -10.61 | 0.01 | 2 | -9.69 | 0.03 | 2 | 4 | 0 | 2 | 28.8 | |
| | | | | | *Min.* | 16 | | | 18.00 | | | -12.57 | | | -10.73 | | | 2 | | | 17.5 | |
| | | | | | *Max.* | 17.1 | | | 20.63 | | | -9.92 | | | -9.56 | | | 5 | | | 28.8 | |
| | | | | | *Mean / SEM* | 16.6 | 0.4 | | 19.3 | 1.3 | | -11.3 | 1.2 | | -10.02 | 0.5 | | 4 | 1 | | 22.4 | 3.4 |
| | PMO 212.662/0 | SS12 | Cervical vertebra | The most anterior | Cervical | 17 | 0.1 | 5 | 16.20 | 0.08 | 3 | -14.27 | 0.08 | 3 | -11.99 | 0.05 | 3 | 9 | 1 | 3 | 16.5 | |
| | PMO 212.662/0 | SS13 | Cervical vertebra | Between SS12 et SS14 | Cervical | 16.4 | 0.1 | 5 | 16.90 | 0.16 | 3 | -13.55 | 0.16 | 3 | -11.95 | 0.07 | 3 | 9 | 1 | 3 | 26.0 | |
| | PMO 212.662/0 | SS14 | Cervical vertebra | Between SS13 and SS15 | Cervical | 16.7 | 0.1 | 5 | | | | | | | | | | | | | 28.3 | |
| | PMO 212.662/0 | SS15 | Cervical vertebra | Between SS14 and SS16 | Cervical | 17 | 0.1 | 5 | 18.30 | 0.08 | 3 | -12.24 | 0.08 | 3 | -10.22 | 0.1 | 3 | 5 | 0 | 3 | 28.0 | |
| | PMO 212.662/0 | SS16 | Cervical vertebra | Between SS15 and SS17 | Cervical | 16.1 | 0.1 | 5 | 18.10 | 0.42 | 2 | -12.46 | 0.42 | 2 | -11 | 0.48 | 2 | 7 | 1 | 2 | 26.7 | |
| | PMO 212.662/0 | SS17 | Cervical vertebra | Between SS16 and SS18 | Cervical | 15.9 | 0.2 | 5 | 18.40 | 0.05 | 3 | -12.11 | 0.05 | 3 | -10.23 | 0.07 | 3 | 4 | 0 | 3 | 20.4 | |
| | PMO 212.662/0 | SS18 | Cervical vertebra | Between SS17 and SS19 | Cervical | 15.9 | 0.2 | 5 | 18.40 | 0.08 | 3 | -12.14 | 0.08 | 3 | -10.01 | 0.28 | 3 | 2 | 2 | 3 | 16.5 | |
| | PMO 212.662/0 | SS19 | Cervical vertebra | Between SS18 and SS20 | Cervical | 16.1 | 0.1 | 5 | 18.40 | 0.06 | 3 | -12.14 | 0.06 | 3 | -10.82 | 0.42 | 3 | 5 | 0 | 3 | 30.2 | |
| | PMO 212.662/0 | SS20 | Cervical vertebra | Between SS19 and SS22 | Cervical | 16.7 | 0 | 5 | 18.92 | 0.04 | 3 | -11.59 | 0.03 | 3 | -9.56 | 0.06 | 3 | 4 | 0 | 3 | 27.1 | |
| | PMO 212.662/0 | SS21 | Cervical vertebra | Between SS20 and SS22 | Cervical | 16.4 | 0.2 | 5 | 18.64 | 0.1 | 3 | -11.86 | 0.1 | 3 | -10.44 | 0.03 | 3 | 5 | 0 | 3 | 29.7 | |
| | PMO 212.662/0 | SS22 | Cervical vertebra | Between SS21 and SS23 | Cervical | 16.3 | 0.1 | 5 | 18.10 | 0.08 | 3 | -12.47 | 0.08 | 3 | -9.89 | 0.03 | 3 | 4 | 0 | 3 | 25.3 | |
| | PMO 212.662/0 | SS23 | Cervical vertebra | The most posterior | Cervical | 15.8 | 0.2 | 5 | 19.20 | 0.13 | 2 | -11.39 | 0.13 | 2 | -9.91 | 0.06 | 2 | 5 | 0 | 2 | 25.3 | |
| | | | | | *Min.* | 15.8 | | | 16.20 | | | -14.27 | | | -11.99 | | | 2 | | | 16.5 | |
| | | | | | *Max.* | 17 | | | 19.20 | | | -11.39 | | | -9.56 | | | 9 | | | 30.2 | |
| | | | | | *Mean / SEM* | 16.4 | 0.4 | | 18.1 | 0.9 | | -12.4 | 0.8 | | -10.5 | 0.8 | | 5 | 2 | | 25.0 | 4.7 |
| | | SS1 | Rib fragment | | Dorsal | 15.9 | 0.2 | 5 | 18.90 | 0.13 | 3 | -11.68 | 0.13 | 3 | -9.79 | 0.04 | 3 | 5 | 0 | 3 | 28.9 | |
| | | SS2 | Rib fragment | | Dorsal | 16.1 | 0.1 | 4 | 18.10 | 0.07 | 3 | -12.44 | 0.07 | 3 | -10.08 | 0.07 | 3 | 4 | 0 | 3 | 26.1 | |
| | | SS3 | Dorsal vertebra | Transversal process | Dorsal | 16.7 | 0.2 | 5 | 18.80 | 0.12 | 2 | -11.76 | 0.12 | 2 | -10.4 | 0.5 | 2 | 4 | 4 | 2 | 26.0 | |
| | | SS4 | Vertebra | | Dorsal | 16.6 | 0.2 | 5 | 19.08 | 0.15 | 3 | -11.43 | 0.15 | 3 | -10.62 | 0.12 | 3 | 4 | 1 | 3 | 19.8 | |
| | | SS5 | Vertebra | | Dorsal | 16.7 | 0.2 | 5 | 16.53 | 0.06 | 3 | -13.9 | 0.06 | 3 | -11.87 | 0.07 | 3 | 8 | 1 | 3 | 20.9 | |
| | | SS6 | Vertebra | | Dorsal | 16.7 | 0.1 | 5 | 18.60 | 0.2 | 3 | -11.99 | 0.2 | 3 | -10 | 0.12 | 3 | 5 | 1 | 3 | 28.1 | |
| | | SS7 | Rib fragment | | Dorsal | 16.7 | 0.1 | 5 | 19.10 | 0.21 | 3 | -11.45 | 0.21 | 3 | -9.85 | 0.03 | 3 | 5 | 0 | 3 | 26.1 | |
| | | SS8 | Rib fragment | | Dorsal | 16.9 | 0.1 | 5 | | | | | | | | | | | | | 24.5 | |
| | | SS10 | Dorsal vertebra | | Dorsal | 16.7 | 0.1 | 5 | | | | | | | | | | | | | 29.4 | |
| | | SS11 | Rib fragment | | Dorsal | 17 | 0.1 | 5 | 18.70 | 0.2 | 3 | -11.85 | 0.2 | 3 | -9.21 | 0.1 | 3 | 4 | 0 | 3 | 22.1 | |
| | | SS24 | Dorsal rib | Distal part | Dorsal | 16.8 | 0.1 | 5 | 18.90 | 0.03 | 3 | -11.61 | 0.03 | 3 | -10.32 | 0.04 | 3 | 7 | 0 | 3 | 31.1 | |
| | | SS25 | Dorsal rib | | Dorsal | 16.6 | 0.2 | 5 | | | | | | | | | | | | | 26.0 | |
| | | SS26 | Dorsal vertebra | | Dorsal | **16.4** | 0.1 | 5 | 12.47 | 0.15 | 3 | -17.83 | -14.68 | 3 | -14.68 | 0.14 | 3 | 18 | 0 | 3 | 18.3 | |
| | | SS27 | Dorsal vertebra | | Dorsal | 16.3 | 0.1 | 4 | | | | | | | | | | | | | 18.6 | |
| | | SS28 | Dorsal rib | Proximal part | Dorsal | 16.1 | 0.2 | 5 | 18.70 | 0.23 | 3 | -11.84 | 0.23 | 3 | -10.23 | 0.1 | 3 | 5 | 0 | 3 | 29.5 | |
| | | SS29 | Dorsal rib | Medial part | Dorsal | 16.1 | 0.2 | 5 | 19.00 | 0 | 3 | -11.6 | 0 | 3 | -10.22 | 0.03 | 3 | 5 | 0 | 3 | 25.0 | |
| | | | | | *Min.* | 15.9 | | | 12.47 | | | -17.83 | | | -14.68 | | | 4 | | | 18.3 | |
| | | | | | *Max.* | 17 | | | 19.10 | | | -11.43 | | | -9.21 | | | 18 | | | 31.1 | |
| | | | | | *Mean / SEM* | 16.5 | 0.3 | | 18.07 | 1.9 | | -12.4 | 1.8 | | -10.6 | 1.4 | | 6 | 4 | | 25.0 | 4.0 |
| | | | | | **Global Min.** | **15.8** | | | **12.47** | | | **-17.83** | | | **-14.68** | | | **2** | | | **16.5** | |
| | | | | | **Global Max.** | **17.1** | | | **20.63** | | | **-9.92** | | | **-9.21** | | | **18** | | | **31.1** | |
| | | | | | **Global Mean / SEM** | **16.5** | **0.4** | | **18.28** | **1.47** | | **-12.25** | **1.42** | | **-10.50** | **1.09** | | **6** | **3** | | **24.3** | **4.2** |
| | | | | | **Global Δ** | **1.3** | | | **8.16** | | | **7.91** | | | **5.47** | | | **16** | | | **14.6** | |
| | | | | | **Global Mid-range** | **16.5** | | | **16.55** | | | **-13.88** | | | **-11.95** | | | **10** | | | **23.8** | |

| Taxa | Coll. N° | #Sample | Skeletal element | Additionnal information | Skeletal region | $\delta^{18}O_p$ (‰, V-SMOW) | | | $\delta^{18}Oc$ (‰, V-SMOW) | | | $\delta^{18}Oc$ (‰, V-PDB) | | | $\delta^{13}C_c$ (‰, V-PDB) | | | wt% $CO_3^{2-}$ | | | wt% $P_2O_5$ | |
|---|---|---|---|---|---|---|---|---|---|---|---|---|---|---|---|---|---|---|---|---|---|---|
| | | | | | | Mean | SD | N | Mean | SD | N | Mean | SD | N | Mean | SD | N | Mean | SD | N | | |
| Elasmosauridae indet. | | EL2 | Humerus | | Appendicular | 18.5 | 0 | 5 | | | | | | | | | | | | | 28.3 | |
| | | EL3 | Humerus | | Appendicular | 18.3 | 0.1 | 4 | | | | | | | | | | | | | 26.7 | |
| | | | | | *Min.* | 18.3 | | | | | | | | | | | | | | | 26.7 | |
| | | | | | *Max.* | 18.5 | | | | | | | | | | | | | | | 28.3 | |
| | | | | | *Mean / SEM* | 18.4 | 0.1 | | | | | | | | | | | | | | 27.5 | 0.8 |
| | | EL16 | « 5th » caudal vertebra | | Caudal | 18.2 | 0.2 | 5 | | | | | | | | | | | | | 26.8 | |
| | | EL17 | « 6th » caudal vertebra | | Caudal | 18.2 | 0.1 | 5 | | | | | | | | | | | | | 30.9 | |
| | | EL18 | « 7th » caudal vertebra | | Caudal | 17.8 | 0.2 | 5 | | | | | | | | | | | | | 27.5 | |
| | | EL19 | « 8th » caudal vertebra | | Caudal | 18.1 | 0.2 | 4 | | | | | | | | | | | | | 27.1 | |
| | | EL20 | « 9th » caudal vertebra | | Caudal | 18 | 0.1 | 3 | | | | | | | | | | | | | 25.0 | |
| | | EL21 | « 10th » caudal vertebra | | Caudal | 17.6 | 0.1 | 4 | | | | | | | | | | | | | 22.4 | |
| | | EL22 | « 11th » caudal vertebra | | Caudal | 18.5 | 0.4 | 4 | | | | | | | | | | | | | 32.8 | |
| | | | | | *Min.* | 17.6 | | | | | | | | | | | | | | | 22.4 | |
| | | | | | *Max.* | 18.5 | | | | | | | | | | | | | | | 32.8 | |
| | | | | | *Mean / SEM* | 18.1 | 0.3 | | | | | | | | | | | | | | 27.5 | 3.2 |
| | | EL4 | « 1st » cervical vertebra | | Cervical | 18.3 | 0.1 | 5 | | | | | | | | | | | | | 23.0 | |
| | | EL5 | « 2nd » cervical vertebra | | Cervical | 17.9 | 0.1 | 5 | | | | | | | | | | | | | 15.8 | |
| | | EL6 | « 3th » cervical vertebra | | Cervical | 18.4 | 0.1 | 5 | | | | | | | | | | | | | 28.8 | |

| | | | | | | | | | | | | | | | | | | | | | | |
|---|---|---|---|---|---|---|---|---|---|---|---|---|---|---|---|---|---|---|---|---|---|---|
| | | EL7 | « 9th » cervical vertebra | | Cervical | 18.4 | 0.1 | 5 | | | | | | | | | | | | | 20.6 | |
| | | EL8 | « 13th » cervical vertebra | | Cervical | 18.4 | 0.2 | 5 | | | | | | | | | | | | | 28.9 | |
| | | EL9 | « 16th » cervical vertebra | | Cervical | 18.1 | 0.2 | 5 | | | | | | | | | | | | | 26.0 | |
| | | | | | *Min.* | 17.9 | | | | | | | | | | | | | | | 15.8 | |
| | | | | | *Max.* | 18.4 | | | | | | | | | | | | | | | 28.9 | |
| | | | | | *Mean / SEM* | 18.3 | 0.2 | | | | | | | | | | | | | | 23.8 | 4.7 |
| | | EL1 | Rib | | Dorsal | 18.3 | 0.1 | 5 | | | | | | | | | | | | | 30.8 | |
| | | EL10 | « 2nd » pectoral vertebra | | Dorsal | 18.2 | 0.1 | 5 | | | | | | | | | | | | | 31.7 | |
| | | EL11 | « 3th » dorsal vertebra | | Dorsal | 17.5 | 0.1 | 2 | | | | | | | | | | | | | 3.5 | |
| | | EL12 | « 6th » dorsal vertebra | | Dorsal | 18.1 | 0.1 | 5 | | | | | | | | | | | | | 23.8 | |
| | | EL13 | « 9th » dorsal vertebra | | Dorsal | 18.4 | 0.1 | 5 | | | | | | | | | | | | | 24.0 | |
| | | EL14 | « 14th » dorsal vertebra | | Dorsal | 18.1 | 0.0 | 5 | | | | | | | | | | | | | 28.0 | |
| | | EL15 | « 2nd » sacral vertebra | | Dorsal | 18.1 | 0.1 | 5 | | | | | | | | | | | | | 23.5 | |
| | | | | | *Min.* | 17.5 | | | | | | | | | | | | | | | 3.5 | |
| | | | | | *Max.* | 18.4 | | | | | | | | | | | | | | | 31.7 | |
| | | | | | *Mean / SEM* | 18.1 | 0.3 | | | | | | | | | | | | | | 23.6 | 8.8 |
| | | | | | **Global Min.** | **17.5** | | | | | | | | | | | | | | | **3.5** | |
| | | | | | **Global Max.** | **18.5** | | | | | | | | | | | | | | | **32.8** | |
| | | | | | **Global Mean / SEM** | **18.2** | **0.3** | | | | | | | | | | | | | | **25.3** | **6.1** |
| | | | | | **Global Δ** | **1.0** | | | | | | | | | | | | | | | **29.3** | |
| | | | | | **Global Mid-range** | **18.0** | | | | | | | | | | | | | | | **18.1** | |

underlined: Data removed because mineralogy altered

**in bold: Data removed because %wt CO3 > 13.4%**

| | Coll. N° | #Sample | Skeletal element | Additionnal information | Skeletal region | $\delta^{18}O_p$ (‰, V-SMOW) | | | $\delta^{18}Oc$ (‰, V-SMOW) | | | $\delta^{18}Oc$ (‰, V-PDB) | | | $\delta^{13}C_c$ (‰, V-PDB) | | | wt% $CO_3^{2-}$ | | | wt% $P_2O_5$ | |
|---|---|---|---|---|---|---|---|---|---|---|---|---|---|---|---|---|---|---|---|---|---|---|
| | | | | | | Mean | SD | N | Mean | SD | N | Mean | SD | N | Mean | SD | N | Mean | SD | N | | |
| *Metriorhynchus* aff. *superciliosus* | | M13 | 1st caudal vertebra | | Caudal | 19.9 | 0.3 | 5 | | | | | | | | | | | | | 27.2 | |
| | | M14 | 7th caudal vertebra | | Caudal | 19.9 | 0.1 | 5 | | | | | | | | | | | | | 22.1 | |
| | | M15 | 14th caudal vertebra | | Caudal | 20 | 0.2 | 5 | | | | | | | | | | | | | 28.9 | |
| | | M16 | 23th caudal vertebra | | Caudal | 20.1 | 0.1 | 5 | | | | | | | | | | | | | 22.7 | |
| | | M17 | caudal vertebra distale | | Caudal | 19.9 | 0.1 | 5 | | | | | | | | | | | | | 27.6 | |
| | | | | | *Min.* | 19.9 | | | | | | | | | | | | | | | 22.1 | |
| | | | | | *Max.* | 20.1 | | | | | | | | | | | | | | | 28.9 | |
| | | | | | *Moy. / SEM* | 20.0 | 0.1 | | | | | | | | | | | | | | 25.7 | 3.1 |
| | | M4 | Rib indt. | | Dorsal | 19.3 | 0.2 | 4 | | | | | | | | | | | | | 19.2 | |
| | | M5 | Dorsal rib | | Dorsal | 19.9 | 0.2 | 4 | | | | | | | | | | | | | 29.2 | |
| | | M6 | Dorsal rib | | Dorsal | 19.9 | 0.1 | 5 | | | | | | | | | | | | | 22.9 | |
| | | M7 | Right ischium | | Dorsal | 20.2 | 0.3 | 5 | | | | | | | | | | | | | 26.4 | |
| | | M8 | Axis | | Dorsal | 20 | 0.2 | 4 | | | | | | | | | | | | | 33.0 | |
| | | M9 | «2nd» dorsal vertebra | | Dorsal | 20.2 | 0.1 | 5 | | | | | | | | | | | | | 26.3 | |
| | | M10 | «5th» dorsal vertebra | | Dorsal | 20.2 | 0.2 | 3 | | | | | | | | | | | | | 28.5 | |
| | | M11 | «13th» dorsal vertebra | | Dorsal | 20.1 | 0.3 | 5 | | | | | | | | | | | | | 26.6 | |
| | | M12 | 1st sacral vertebra | | Dorsal | 19.8 | 0.2 | 5 | | | | | | | | | | | | | 22.1 | |
| | | M18 | 3th cervical vertebra | | Dorsal | 19.7 | 0.1 | 4 | | | | | | | | | | | | | 28.6 | |
| | | M19 | 3th cervical vertebra | | Dorsal | 20.3 | 0.2 | 5 | | | | | | | | | | | | | 27.8 | |
| | | M21 | Bones of the pelvic girdle indt. | | Dorsal | 19.3 | 0.1 | 5 | 27.3 | 0.2 | 3 | | | | -8.22 | 0.16 | - | 5 | 1 | - | 17.7 | |
| | | | | | *Min.* | 19.3 | | | | | | | | | | | | | | | 17.7 | |
| | | | | | *Max.* | 20.3 | | | | | | | | | | | | | | | 33.0 | |
| | | | | | *Moy. / SEM* | 19.9 | 0.3 | | | | | | | | | | | | | | 25.7 | 4.4 |
| | | M1 | Maxillary | | Skull | 19.7 | 0.2 | 4 | | | | | | | | | | | | | 24.4 | |
| | | M2 | Mandible | | Skull | 19.9 | 0.1 | 5 | | | | | | | | | | | | | 24.2 | |
| | | M3 | Squamosal | | Skull | 19.4 | 0.2 | 4 | | | | | | | | | | | | | 25.9 | |
| | | | | | *Min.* | 19.4 | | | | | | | | | | | | | | | 24.2 | |
| | | | | | *Max.* | 19.9 | | | | | | | | | | | | | | | 25.9 | |
| | | | | | *Moy. / SEM* | 19.7 | 0.3 | | | | | | | | | | | | | | 24.9 | 0.9 |
| | | | | | **Global Min.** | **19.3** | | | | | | | | | | | | | | | **17.7** | |
| | | | | | **Global Max.** | **20.3** | | | | | | | | | | | | | | | **33.0** | |
| | | | | | **Global Mean / SEM** | **19.9** | **0.3** | | | | | | | | | | | | | | **25.6** | **3.7** |
| | | | | | **Global Δ** | **1.0** | | | | | | | | | | | | | | | **15.3** | |
| | | | | | **Global Mid-range** | **19.8** | | | | | | | | | | | | | | | **25.4** | |
