## Supplementary Table 2 for "Reassessment of body temperature and thermoregulation strategies in Mesozoic marine reptiles"

| Locality | Stage | Paleolatitude °N (van Hinsbergen et al. 2015) | $\delta^{18}O_{fish}$ (‰, V-SMOW) | | | $\delta^{18}O_{seawater}$ estimates from Alberti et al. 2020 (‰ V-SMOW) | $T_{seawater}$ (°C) | |
|---|---|---|---|---|---|---|---|---|
| | | | N | Mean | SEM | | Mean | SEM |
| Smallmouth Sands, England | Kimmeridgian | 36.8 [34.1 - 39.7] | 21 | 20.3 | 0.8 | -0.4 | 24 | 4 |
| Peterborough, England | Late Callovian - Early Oxfordian | 39.6 [36.7 - 41.7] | 12 | 20.4 | 0.5 | -0.6 | 23 | 2 |
| "Vache Noires Cliffs", France | Callovian | 36.4 [33.5 - 39.6] | 5 | 20.8 | 0.4 | -0.4 | 22 | 2 |
| Quarries “Les Lourdines”, France | Middle Callovian | 33.8 [30.9 - 36.9] | 3 | 19.9 | 0.5 | -0.3 | 27 | 1 |
| Oulad abdoun, Morocco | Maastrichtian | 20.3 [19.2 - 21.5] | 4 | 20.4 | 0.9 | 0.0 | 25 | 4 |
| Cambridge, England | Late Albian | 43.6 [42.2 - 45.2] | 2 | 20.2 | 0.1 | -0.7 | 23 | 0 |
| Westbury, England | Early Kimmeridgian | 37.5 [34.8 - 40.4] | 4 | 19.5 | 0.1 | -0.5 | 27 | 0 |
| Sorel, France | Sinemurian | 40.6 [39.2 - 42.0] | 1 | 19.7 | - | -0.6 | 26 | - |
| Crussol, France | Middle Oxfordian | 32.8 [30.0 - 36.0] | 2 | 20.3 | 1 | -0.3 | 25 | 4 |
| Bourgogne, France | Early Oxfordian | 35.2 [32.3 - 38.3] | 1 | 21.1 | - | -0.4 | 21 | - |
| Asen, Sweden | Early Campanian | 53.6 [52.3 - 54.9] | 4 | 21.1 | 0.9 | -1.1 | 18 | 4 |
| Ullstorp, Sweden | Early Campanian | 48.4 [47.1 - 49.7] | 4 | 20.4 | 0.6 | -0.9 | 21 | 3 |
| "Les Ardilles", France | Kimmeridgian | 34.5 [31.8 - 37.4] | 1 | 18.1 | - | -0.4 | 34 | - |
| Slottsmøya Member, Spitsbergen | Volgian | 65.5 [62.8 - 68.4] | | | | -1.4 | 14 | 2 |

| Locality | **Metriorhynchidae** | | | | | | **Plesiosauria** | | | | | | **Ichthyosauria** | | | | | |
|---|---|---|---|---|---|---|---|---|---|---|---|---|---|---|---|---|---|---|
| | $\delta^{18}O_p$ (‰ V-SMOW) | | | $T_{body}$ (°C) | | $T_{body} - T_{seawater}$ | $\delta^{18}O_p$ (‰ V-SMOW) | | | $T_{body}$ (°C) | | $T_{body} - T_{seawater}$ | $\delta^{18}O_p$ (‰ V-SMOW) | | | $T_{body}$ (°C) | | $T_{body} - T_{seawater}$ |
| | N | Mean | SD | Mean | SEM | | N | Mean | SEM | Mean | SEM | | N | Mean | SEM | Mean | SEM | |
| Smallmouth Sands, England | 8 | 20.4 | 0.4 | 27 | 2 | 3 | 1 | 20 | - | 29 | - | 5 | 3 | 19.5 | 0.1 | 32 | 0 | 8 |
| Peterborough, England | 11 | 20.3 | 0.6 | 27 | 4 | 4 | 4 | 20.2 | 0.6 | 29 | 4 | 6 | 6 | 18.8 | 0.2 | 34 | 1 | 11 |
| "Vache Noires Cliffs", France | 7 | 20.9 | 0.5 | 25 | 3 | 3 | 2 | 20.5 | 0.2 | 27 | 1 | 5 | | | | | | |
| Quarries “Les Lourdines”, France | 2 | 19.4 | 0.2 | 32 | 1 | 5 | | | | | | | | | | | | |
| Oulad abdoun, Morocco | | | | | | | 3 | 19.8 | 0.6 | 32 | 3 | 7 | | | | | | |
| Cambridge, England | | | | | | | 2 | 19.2 | 0.6 | 31 | 3 | 8 | 3 | 18.7 | 0.2 | 34 | 1 | 10 |
| Westbury, England | | | | | | | | | | | | | 2 | 19.6 | 0.6 | 31 | 3 | 4 |
| Sorel, France | | | | | | | 1 | 18.8 | - | 34 | - | 8 | | | | | | |
| Crussol, France | | | | | | | 1 | 19.1 | - | 34 | - | 9 | | | | | | |
| Bourgogne, France | | | | | | | 1 | 19.3 | - | 32 | - | 11 | 1 | 18.8 | - | 35 | - | 14 |
| Asen, Sweden | | | | | | | 2 | 19.6 | 0.4 | 28 | 2 | 10 | | | | | | |
| Ullstorp, Sweden | | | | | | | 2 | 20.1 | 0.5 | 27 | 0 | 6 | | | | | | |
| "Les Ardilles", France | | | | | | | | | | | | | 1 | 18.3 | - | 37 | | 3 |
| Slottsmøya Member, Spitsbergen | | | | | | | | | | | | | 7 | 15.3 | 0.4 | 45 | 2 | 31 |
