## Supplementary Table 3 for "Reassessment of body temperature and thermoregulation strategies in Mesozoic marine reptiles"

| Specimen | #Sample | Type of material | Position ν1-$PO_4^{3-}$ ($cm^{-1}$) | ν1-$PO_4^{3-}$ FWHM ($cm^{-1}$) |
|---|---|---|---|---|
| | Hydroxylapatite | | 962.1 | 9.2 |
| Elasmosauridae indet. (MHNLM.2005.16) | EL4 | Cervical vertebra | 959.7 | 18.2 |
| | EL10 | Pectoral vertebra | 962.7 | 13.2 |
| | EL12 | Dorsal vertebra | 942.9 | 28.1 |
| | EL15 | Sacral vertebra | 952.8 | 18.1 |
| | EL16 | Caudal vertebra | 962.7 | 13.2 |
| | EL20 | Caudal vertebra | 962.7 | 13.2 |
| | EL22 | Caudale vertebra | 961.4 | 14.9 |
| | | Mean | 957.8 | 17.0 |
| | | SEM | 6.9 | 5.0 |
| Ichthyosauria indet. | ICK1 | Dorsal vertebra | 954.5 | 18.1 |
| | ICK3 | Dorsal vertebra | 954.5 | 21.4 |
| | ICK4 | Dorsal vertebra | 962.7 | 16.5 |
| | ICK5 | Dorsal vertebra | 952.8 | 19.8 |
| | ICK6 | Dorsal vertebra | 952.8 | 19.8 |
| | ICK7 | Dorsal vertebra | 964.4 | 11.5 |
| | ICK8 | Dorsal vertebra | 952.8 | 16.5 |
| | ICK9 | Caudal vertebra | 951.2 | 18.2 |
| | ICK10 | Dorsal vertebra | 964.4 | 14.8 |
| | ICK11 | Dorsal vertebra | 949.5 | 21.6 |
| | ICK12 | Dorsal vertebra | 964.4 | 14.8 |
| | ICK13 | Caudal vertebra | 956.1 | 13.2 |
| | ICK14 | Caudal vertebra | 951.2 | 21.4 |
| | ICK18 | Rib | 962.7 | 13.2 |
| | ICK19 | Right nasal | 956.1 | 16.5 |
| | ICK22 | Left humerus | 952.8 | 16.5 |
| | ICK23 | Left radius | 952.8 | 18.2 |
| | ICK28 | Phalange | 939.6 | 33.1 |
| | ICK31 | Phalange? | 962.7 | 14.8 |
| | ICK34 | Left angular | 962.7 | 14.8 |
| | ICK36 | Tooth | 962.7 | 11.5 |
| | ICK38 | Left premaxilla | 962.7 | 13.2 |
| | | Mean | 956.6 | 17.2 |
| | | SEM | 6.3 | 4.6 |
| *Keilhauia nui* (PMO 222.655) | Mi2 | Dorsal vertebra | 997.6 | 26.3 |
| | Mi3 | Presacral vertebra | 949.5 | 21.5 |
| | Mi8 | Presacral vertebra | 949.5 | 19.8 |
| | Mi9 | Presacral vertebra | 1023.4 | 21.2 |
| | Mi14 | Caudal vertebra | 942.9 | 34.7 |
| | Mi16 | Caudal vertebra | 1005.4 | 18 |
| | Mi23 | Rib | 1004.2 | 11.5 |
| | | Mean | 981.8 | 21.9 |
| | | SEM | 30.8 | 6.7 |
| *Palvennia hoybergeti* (PMO 222.669) | PH12 | Dorsal vertebra | 970.9 | 14.8 |
| | PH14 | Left dentary | 947.9 | 18.2 |
| | PH15 | Premaxilla | 972.8 | 26.5 |
| | | Mean | 963.9 | 19.8 |
| | | SEM | 11.3 | 4.9 |
| *Keilhauia* sp. (PMO 222.667) | K1 | Tooth | 947.9 | 21.5 |
| | K2 | Tooth | 951.2 | 18.2 |
| | K3 | Tooth | 961.4 | 18.2 |
| | K10 | Limb bone | 946.2 | 23.1 |
| | K19 | Preflexural vertebra | 947.9 | 23.1 |
| | K24 | Rib | 954.8 | 19.8 |

| | | | | |
|---|---|---|---|---|
| | K25 | Coracoid | 951.47 | 28.2 |
| | | Mean | 951.6 | 21.7 |
| | | SEM | 4.8 | 3.3 |
| *Colymbosaurus svalbardensis* (PMO 222.663) | G1 | Right fibula | 961.4 | 14.8 |
| | G12 | III,2 | 944.6 | 28.1 |
| | G16 | III, 13 | 1011.9 | 11.4 |
| | G22 | Left humerus | 949 | 11.5 |
| | G23 | Right humerus | 959.7 | 9.9 |
| | G33 | Rib fragment | 942.9 | 28.1 |
| | G38 | Caudal vertebra | 959.7 | 11.6 |
| | G41 | Rib | 949.8 | 26.5 |
| | | Mean | 959.9 | 17.7 |
| | | SEM | 20.8 | 7.7 |
| Cryptoclididae indet. (PMO 212.662) | SS9 | Propodial fragment | 951.5 | 24.8 |
| | SS11 | Rib | 959.7 | 16.5 |
| | SS16 | Cervical vertebra | 947.9 | 21.5 |
| | SS20 | Cervical vertebra | 947.9 | 21.5 |
| | SS21 | Cervical vertebra | 959.4 | 8.2 |
| | SS23 | Cervical vertebra | 958.1 | 23.2 |
| | SS23 | Cervical vertebra | 961.4 | 18.2 |
| | SS24 | Rib | 958.1 | 18.2 |
| | SS26 | Dorsal vertebra | 958.1 | 13.2 |
| | SS28 | Rib | 961.4 | 14.9 |
| | SS29 | Dorsal rib | 959.4 | 8.2 |
| | SS31 | Left radius | 951.4 | 26.5 |
| | | Mean | 956.2 | 17.9 |
| | | SEM | 4.8 | 5.8 |
